# Detection of Cell-Cell Interactions via Photocatalytic Cell Tagging

**DOI:** 10.1101/2021.10.04.463039

**Authors:** Rob C. Oslund, Tamara Reyes-Robles, Cory H. White, Jake H. Tomlinson, Kelly A. Crotty, Edward P. Bowman, Dan Chang, Vanessa M. Peterson, Lixia Li, Silvia Frutos, Miquel Vila-Perelló, David Vlerick, Karen Cromie, David H. Perlman, Samantha D. O’Hara, Lee R. Roberts, Grazia Piizzi, Erik C. Hett, Daria J. Hazuda, Olugbeminiyi O. Fadeyi

**Affiliations:** Merck Exploratory Science Center, Merck & Co., Inc., Cambridge, Massachusetts, USA; Discovery Research, Merck & Co., Inc., San Francisco, California, USA; Genetics and Pharmacogenomics, Merck & Co., Inc., Boston, Massachusetts, USA; SpliceBio S.L., Barcelona, Spain; Ablynx, A Sanofi Company, Zwijnaarde, East Flanders, Belgium; Infectious Diseases and Vaccine Research, Merck & Co., Inc., West Point, Pennsylvania, USA

## Abstract

Cell-cell interactions drive essential biological processes critical to cell and tissue development, function, pathology, and disease outcome. The growing appreciation of immune cell interactions within disease environments has led to significant efforts to develop protein- and cell-based therapeutic strategies. A better understanding of these cell-cell interactions will enable the development of effective immunotherapies. However, characterizing these complex cellular interactions at molecular resolution in their native biological contexts remains challenging. To address this, we introduce photocatalytic cell tagging (PhoTag), a modality agnostic platform for profiling cell-cell interactions. Using photoactivatable flavin-based cofactors, we generate phenoxy radical tags for targeted labeling at the cell surface. Through various targeting modalities (e.g. MHC-Multimer, antibody, single domain antibody (VHH)) we deliver a flavin photocatalyst for cell tagging within monoculture, co-culture, and peripheral blood mononuclear cells. PhoTag enables highly selective tagging of the immune synapse between an immune cell and an antigen-presenting cell through targeted labeling at the cell-cell junction. This allowed for the ability to profile gene expression-level differences between interacting and bystander cell populations. Given the modality agnostic and spatio-temporal nature of PhoTag, we envision its broad utilization to detect and profile intercellular interactions within an immune synapse and other confined cellular regions for any biological system.

## Introduction

Direct cell-cell interactions play an essential role in the development and biological function of all multicellular organisms^1^. These interactions occur through protein-, glycan-, and/or lipid-mediated physical contact at the plasma membrane to form cellular junctions, adhesions, or synapses in either stable or transient fashion^1,2^. It is these critical interactions that drive the organization of cells into tissues and other complex cellular environments with important consequences on human health. A key example is the tumor microenvironment where interactions between cancer, endothelial, stromal, and immune cells dictate tumor progression^3,4^. Within this context, T cells form immunological synapses with antigen-presenting cells (APCs) through binding events between the T Cell Receptor (TCR) and the peptide-MHC complex (pMHC), adhesion molecules, and costimulatory/checkpoint receptor-ligand pairs on T cell and APC cell surfaces^5,6^. The precise spatial and temporal organization of these surface proteins at the T cell-APC interface governs critical immunological signaling events to determine tumor immune response outcomes^5^. The growing appreciation for the role of T cells in driving tumor immune responses has spurred significant efforts in developing checkpoint inhibitor and adoptive T cell transfer therapies^7^. This has consequently led to an increased need to gain insight into cellular contacts formed between immune cells and their cellular targets.

However, probing interactions within a cellular synapse is a difficult task. The spatially restricted distance of the synapse and the plethora of cell types and proteins involved places a significant burden on identifying suitable technologies that can adequately profile within this environment. A recent approach to explore cell-cell interactions uses enzyme-based cell tagging methods to capture interacting cells and their protein environments^8-12^. While this is an attractive strategy for studying interacting cells, the enzyme size and strict cell engineering requirement for enzyme expression can limit its application to cells compatible with recombinant protein expression and non-sterically restricted junctions. We recently disclosed a novel microenvironment-mapping platform^13^ that exploits photocatalytic carbene generation for antibody target identification and mapping protein environments; however, the short half-life of the carbene species limits efficient tagging and isolation of interacting cells for downstream analysis. We sought to develop a small molecule photocatalyst to generate a longer-lived reactive species for intercellular tagging. This new approach will be compatible with any targeting modality to access within confined cellular regions such as a synaptic junction (**Figure 1a**).

**Figure 1.**
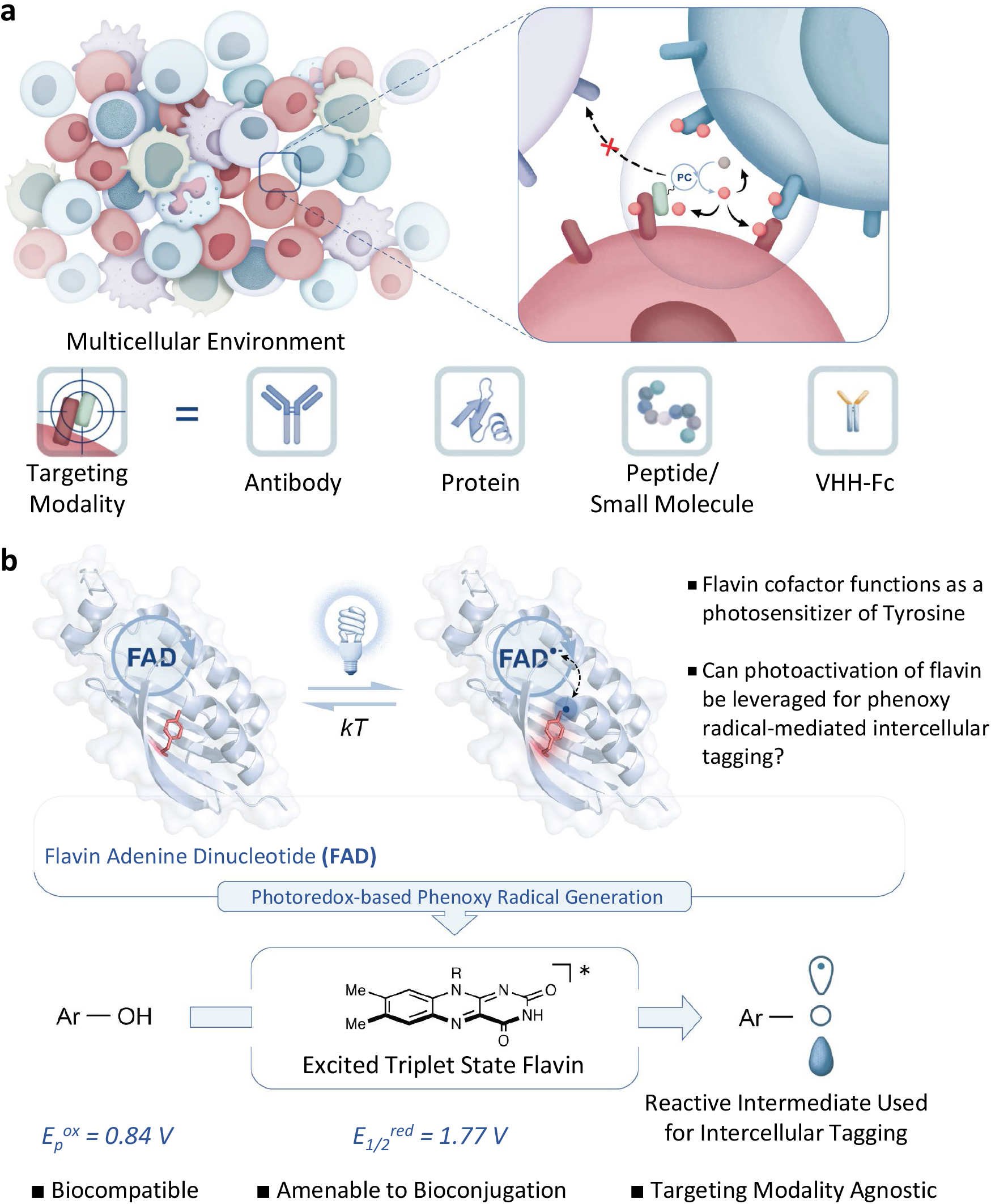
Profiling Multicellular Interaction Environments via Photocatalytic Cell Tagging (PhoTag). a) A small molecule photocatalyst (PC) is targeted to cell surface regions, such as a cellular synapse for light-activated covalent cellular tagging. The small molecule nature of the photocatalyst makes it compatible with diverse targeting modalities. b) Overview of design concept using visible-light activated flavin to generate phenoxy radical tags. Blue light photoreceptors use flavin as a cofactor to oxidize electron rich amino acids such as tyrosines upon light activation. In this work, a photocatalyst system is developed to generate reactive phenoxy radical labeling species for targeted protein labeling within intercellular environments.

Protein electron transfer reactions play crucial roles in biological and bioenergetic redox processes^14-16^. Among the amino acids involved, tyrosine serves as an efficient electron transfer intermediate^17-21^. Indeed, the intrinsic reactivity of tyrosyl radicals generated from tyrosine oxidation has been effectively harnessed in peroxidase-based proximity labeling technology, where tyramide is enzymatically oxidized by heme-based peroxidases to promote labeling of proteins in proximity to the peroxidase-tagged protein of interest^22-27^. The reactivity of the radical is an important factor controlling the labeling radius^28^. However, the extremely rapid generation^29^ of the phenol radical and the large size of the peroxidase enzyme limits its utility in tight cellular junctions. On the other hand, flavin cofactors in the form of flavin mononucleotide (FMN) and flavin adenine dinucleotide (FAD) are ubiquitously used within a wide range of cellular contexts as photosensitizers to oxidize electron-rich amino acid side chains, including the single electron oxidation of tyrosine residues (**Figure 1b**)^30^.

We therefore questioned whether a reactive tyramide radical intermediate could be generated via flavin activation in a light-controlled manner. Our design plan would exploit the excited states of the flavin cofactor (E_1/2_^red^ = 1.77 V versus saturated calomel electrode (SCE)) that is capable of oxidizing phenol substrates (E_ox_ = 0.84 V) via single electron transfer (SET) to generate the reactive phenoxy radical cell tagging intermediate in close proximity to the cell-cell interface of interest (**Figure 1b**)^31-33^. Herein we report the development of a novel, temporally-controlled, flavin-based photocatalyst system that we call **Pho**tocatalytic Cell **Tag**ging (PhoTag) (**Figure 1a**). Using PhoTag, we demonstrate antibody-based targeted labeling of cell surfaces as well as the use of MHC-Multimers for antigen-specific tagging on cells isolated from whole blood. We further demonstrate the ability to achieve highly selective synaptic tagging within a two-cell synapse using a VHH-based flavin conjugate. This selective biotinylation approach enabled the ability to specifically label and isolate uniquely interacting cells from a cell suspension to profile gene expression level differences.

## Results

### Identification of Flavins as Protein Labeling Photocatalysts

In order to explore the feasibility of using PhoTag, we first examined whether the reaction between capped tyrosine and a biotin phenol substrate could be accomplished in the presence of a riboflavin photocatalyst. In a biocompatible solvent (PBS) and with blue LED irradiation, we observed the formation of the desired coupled product in moderate conversion (**Supporting Figure 1**). Based on this promising result, we proceeded to test riboflavin and other suitable photocatalysts for labeling at the protein level (**Figure 2a**), by monitoring the efficiency of various photocatalysts to induce labeling of bovine serum albumin (BSA) with biotin tyramide under visible light irradiation (455 nm) (**See Supporting Information, Bio-photoreactor**). From an initial screening of various catalytic flavins and other oxidizing photocatalysts by western blot, we found that lumiflavin and riboflavin tetraacetate (RFT) yielded the highest levels of protein biotinylation (**Supporting Figures 2 and 3**). Notably, Ru(bpy)_3_, known to induce photooxidation/crosslinking of proteins^34-36^, required the presence of a strong oxidant (ammonium persulfate) to induce moderate protein biotinylation (**Supporting Figure 2**). We next expanded the labeling approach into other proteins and observed light-dependent protein biotinylation in the presence of RFT and biotin tyramide (**Figure 2b**). Peptide LC-MS/MS analysis of RFT-labeled proteins revealed predominant labeling at solvent-exposed tyrosine residues (**Supporting Figures 4-8**).

**Figure 2.**
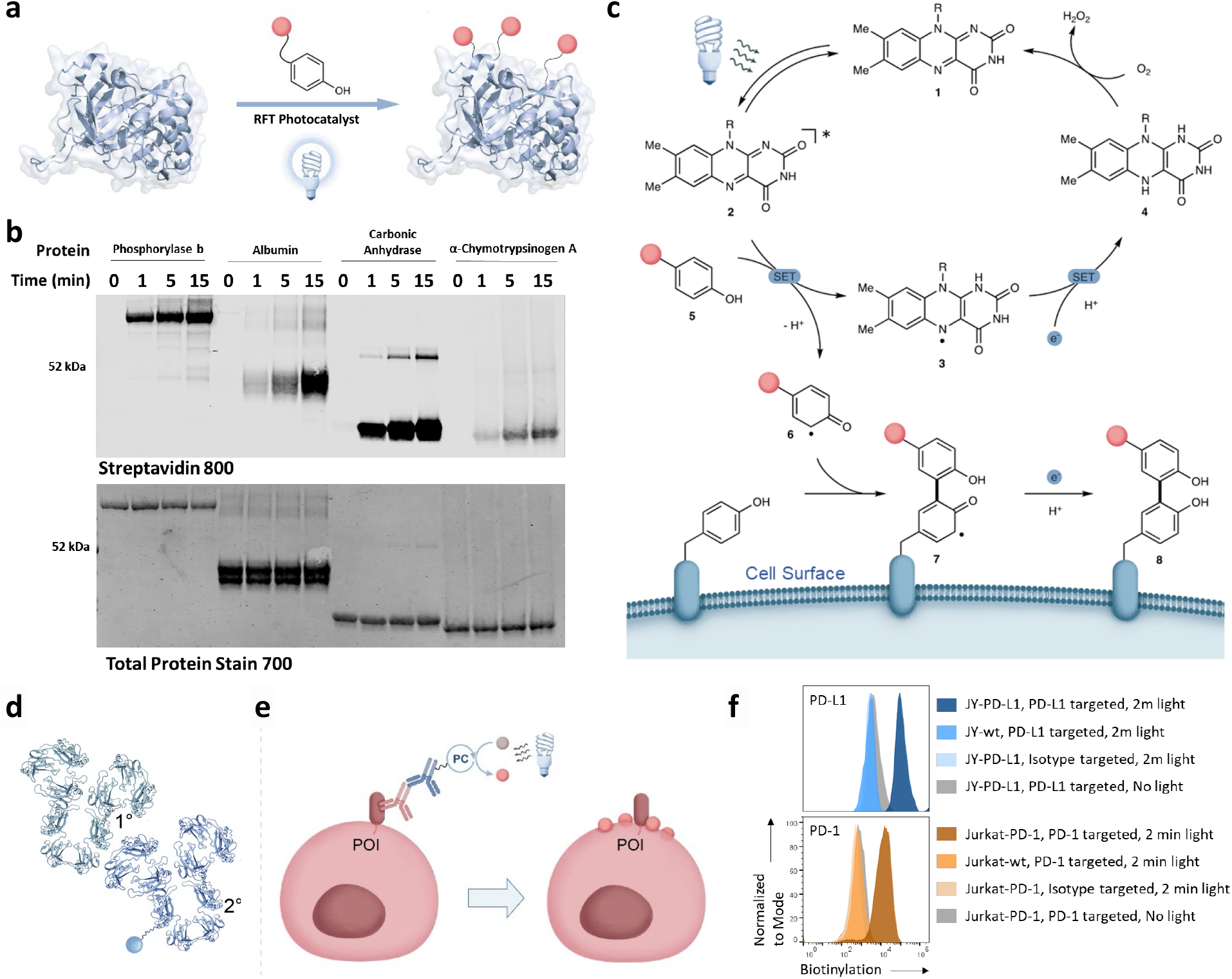
Use of flavins for photocatalytic protein labeling. a) Schematic depicting the use of riboflavin tetraacetate (RFT) and visible light to activate a phenol tag for protein labeling. b) Western blot analysis of light- and time-dependent biotinylation on Phosphorylase B, Albumin, Carbonic anhydrase, and Chymotrypsinogen A using RFT and biotin tyramide. c) Proposed reaction mechanism for flavin-based protein labeling. d) Ab-PhoTag system consisting of a primary antibody (1°) and secondary antibody (2°) labeled with RFT photocatalyst (blue circle) for cell surface targeted labeling. e) Schematic depicting cell surface biotinylation through targeted labeling on a protein of interest (POI) using a primary/secondary antibody flavin conjugate system. f) Flow cytometry analysis of targeted labeling of PD-L1 on JY cells or PD-1 on Jurkat cells. Surface biotinylation occurs when the protein is expressed on the cell surface and in the presence of visible light.

A unique feature of this approach is the dependency of the catalyst on light for temporal control of photoactivation. Indeed, we found that the extent of protein biotinylation can be finely tuned by the catalyst concentration, light duration, and intensity unlike the peroxidase-based enzyme method (**Supporting Figure 9**). Furthermore, protein labeling through a light ON/OFF experiment clearly demonstrates that constant blue LED illumination is required for labeling to proceed (**Supporting Figure 10**). We also found this method was compatible with phenol probes containing bioorthogonal alkyne, azide, DBCO, and fluorophore groups for protein labeling (**Supporting Figure 11**).

A proposed mechanism for flavin-based photo-protein labeling is shown in **Figure 2c**. Photoexcitation of the flavin catalyst **1** under blue LED illumination results in the formation of the flavin singlet excited state, that rapidly generates a triplet-excited flavin **2** via intersystem crossing (ISC)^37^. Next, a single electron transfer from biotin tyramide **5** to the strongly oxidizing triplet-excited flavin species should generate the corresponding phenoxy radical **6** upon deprotonation of the radical cation. Nucleophilic addition of mainly tyrosine residues into the phenoxy radical intermediate **6** should provide the cross-coupled phenol adduct **8** followed by regeneration of the ground state flavin photocatalyst **1** (**Supporting Figures 12 and 13**).

To test whether PhoTag could be used in a targeted fashion, we prepared an antibody flavin conjugate to drive localization of RFT to a protein of interest (**Supporting Figure 14**). Using a primary/secondary antibody flavin conjugate system (**Figure 2d**; hereafter, Ab-PhoTag) where the primary antibody binds the protein of interest and a secondary antibody flavin conjugate recognizes the primary antibody for targeted delivery of the RFT photocatalyst, we demonstrated targeted biotinylation on purified proteins (CD45 or PD-L1) (**Supporting Figure 15**). We next performed Ab-PhoTag on live cells (**Figure 2e**) through targeted labeling of PD-L1 on JY cells overexpressing PD-L1 or PD-1 on Jurkat cells overexpressing PD-1. Targeted labeling of these receptors led to increased cell surface biotinylation, in stark contrast to the background levels observed in the absence of light irradiation, targeting with an isotype antibody, or lack of PD-L1 or PD-1 expression (**Figure 2f**). A similar increase in cell surface biotinylation over isotype was achieved through Ab-PhoTag-targeted labeling of CD86 on JY cells or CD45 on Jurkat cells (**Supporting Figure 16**).

### Antigen-specific T Cell Tagging Using PhoTag System

To further explore the compatibility of the PhoTag system with other modalities, we investigated the use of MHC-Multimers^38^ as a means to direct the flavin photocatalyst to the T cell surface for selective tagging of antigen-specific T cells within a population of human peripheral blood mononuclear cells (PBMCs). CD8+ T cells were targeted using MHC-Multimers displaying HLA A02 and an antigen peptide derived from cytomegalovirus (CMV) 65 kDa phosphoprotein (pp65), NLVPMVATV, or a negative control peptide that is not recognized by PBMCs. These MHC-Multimers were pre-labeled with biotin-RFT resulting in an MHC Multimer-RFT complex (MHC-PhoTag) that can selectively target the flavin photocatalyst to CMV pp65 specific CD8+ T cells for selective biotinylation (**Figure 3a and Supporting Information, Labeling of MHC Multimer with RFT-Biotin**). The MHC-PhoTag system was then tested on purified HLA A02 restricted CD8+ T cells enriched for CMV antigen-specificity resulting in selective biotinylation of CMV-specific CD8+ T cells (**Supporting Figure 17**). This system was next used to biotinylate CMV-specific T cells from human donor PBMCs. Efficient biotinylation occurred in a time and visible light-dependent manner on a small subset of CD8+ T cells that required targeting with the pp65-derived HLA*0201-restricted peptide (**Figure 3b**). Encouraged by this result, we next selected two human PBMC donors and performed MHC-PhoTag on CMV specific CD8+ T cells. After targeted labeling of cells (**Figure 3c**), viable biotinylated CD8+ T cells were then sorted from the larger PBMC population and analyzed by single cell T cell receptor (TCR) sequencing (**Figure 3d**). The paired alpha/beta TCR clonotypes obtained from single cell TCR sequencing revealed a unique set of TCR clonotypes for each donor with high predicted affinity for the pp65 peptide antigen comparable to results obtained using MHC-Multimers labeled with Phycoerythrin (PE) dye (**Figure 3d**). Collectively, these results demonstrate the utility of MHC-PhoTag technology for targeted tagging of antigen-specific T cells and suggest potential use of this technology more broadly to study or manipulate clinical samples *ex vivo* via selective covalent installation of any label.

**Figure 3.**
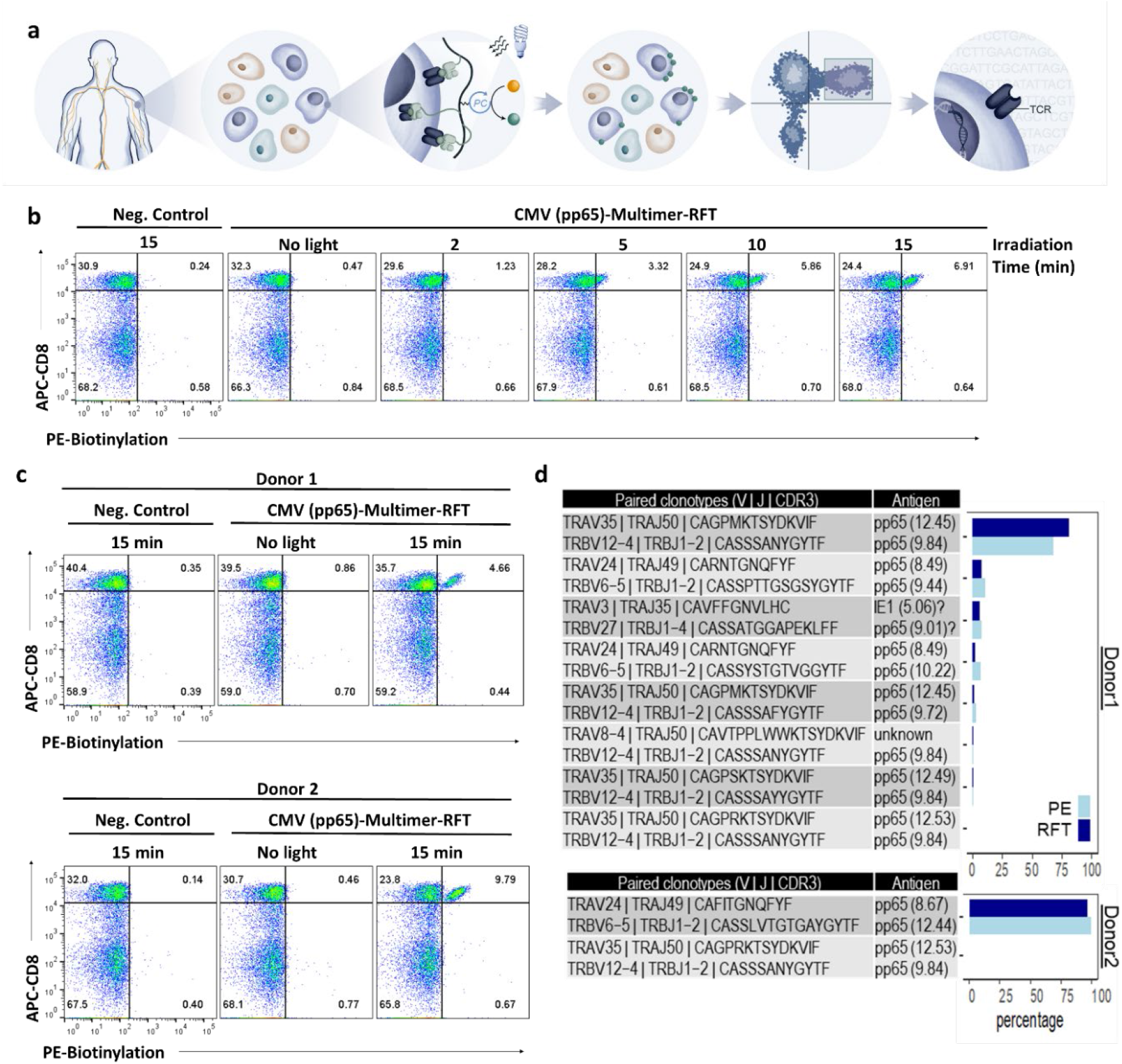
Selective covalent tagging of antigen-specific CD8+ T cell populations using MHC-PhoTag. a) Schematic showing the use of MHC-Multimers labeled with RFT (MHC-PhoTag) for selective labeling of unique CD8+ T cell populations within PBMCs. The selectively labeled cells are isolated by cell sorting and subjected to TCR sequencing analysis. b) Flow cytometry analysis of PBMCs targeted with RFT-labeled MHC-Multimers displaying the CMV pp65-derived HLA*0201-restricted peptide (NLVPMVATV) shows light exposure time-dependent biotinylation of unique CD8+ T cell populations. c) RFT-labeled MHC Multimers were used to selectively biotinylate CD8+ T cell populations from two different donors (Top, Donor 1,; Bottom, Donor 2) using multimers displaying the CMV pp65-derived HLA*0201-restricted peptide (NLVPMVATV). d) Paired alpha beta TCR clonotypes from single-cell sequencing of pp65-specific CD8+ T cells that were enriched with MHC-Multimer-PE (PE) or MHC-Multimer-RFT (RFT) methods from PBMCs of two separate donors. Percentages of each clonotype (with cell count ≥ 3) enriched with the two methods are shown in the respective bar graphs (PE, light blue; RFT, dark blue). VJ gene usage, CD3 amino acid sequences, antigen motifs and informativeness annotated with VDJdb are shown in the table. All paired clonotypes except one (unknown) display pp65-antigen recognition-motifs with high confidence.

### Targeted Labeling at the Immune Synapse Using PhoTag in a T cell/APC Co-culture System

To determine if targeted PhoTag could be used for tagging cell-cell interactions within the immune synapse, we utilized a T cell and APC co-culture system consisting of Jurkat PD-1 expressing T-cells (Jurkat-PD-1) and Raji PD-L1 expressing B cells (Raji-PD-L1) activated with the staphylococcal enterotoxin D (SED) superantigen that enables MHC class II/TCR engagement and signaling independent of antigen presentation^39^. Raji-PD-L1 cells were pre-labeled with α-PD-L1 using the Ab-PhoTag system followed by co-culture with Jurkat-PD-1 cells in the presence of SED and subsequent visible light irradiation. Tunable, light-dependent labeling of both PD-L1-targeted Raji cells in *cis* and the Jurkat-PD-1 cells in *trans* was observed (**Supporting Figures 18 and 19**). This result was supported by confocal imaging where clear evidence of transcellular biotinylation was observed using Ab-PhoTag, whereas excessive labeling on both cell types was observed with peroxidase-based proximity labeling (**Supporting Figure 18**).

While Ab-PhoTag directed at the immune synapse was successful at labeling cell-cell interfaces and achieving transcellular labeling, the abundant extrasynaptic, *cis*-biotinylation on the targeted cell limits the ability to covalently tag and isolate interacting cells. Given that size-based segregation of protein ectodomains occurs within plasma membrane regions close to the points of contact within an immune synapse, due to a restricted intermembrane distance (∼14 nm, **Figure 4a**)^40,41^, we reasoned that the Ab-PhoTag system consisting of a primary and secondary antibody may have limited access into the synapse. This compelled us to explore the use of smaller targeting species in conjunction with PhoTag for selective immune synapse tagging.

**Figure 4.**
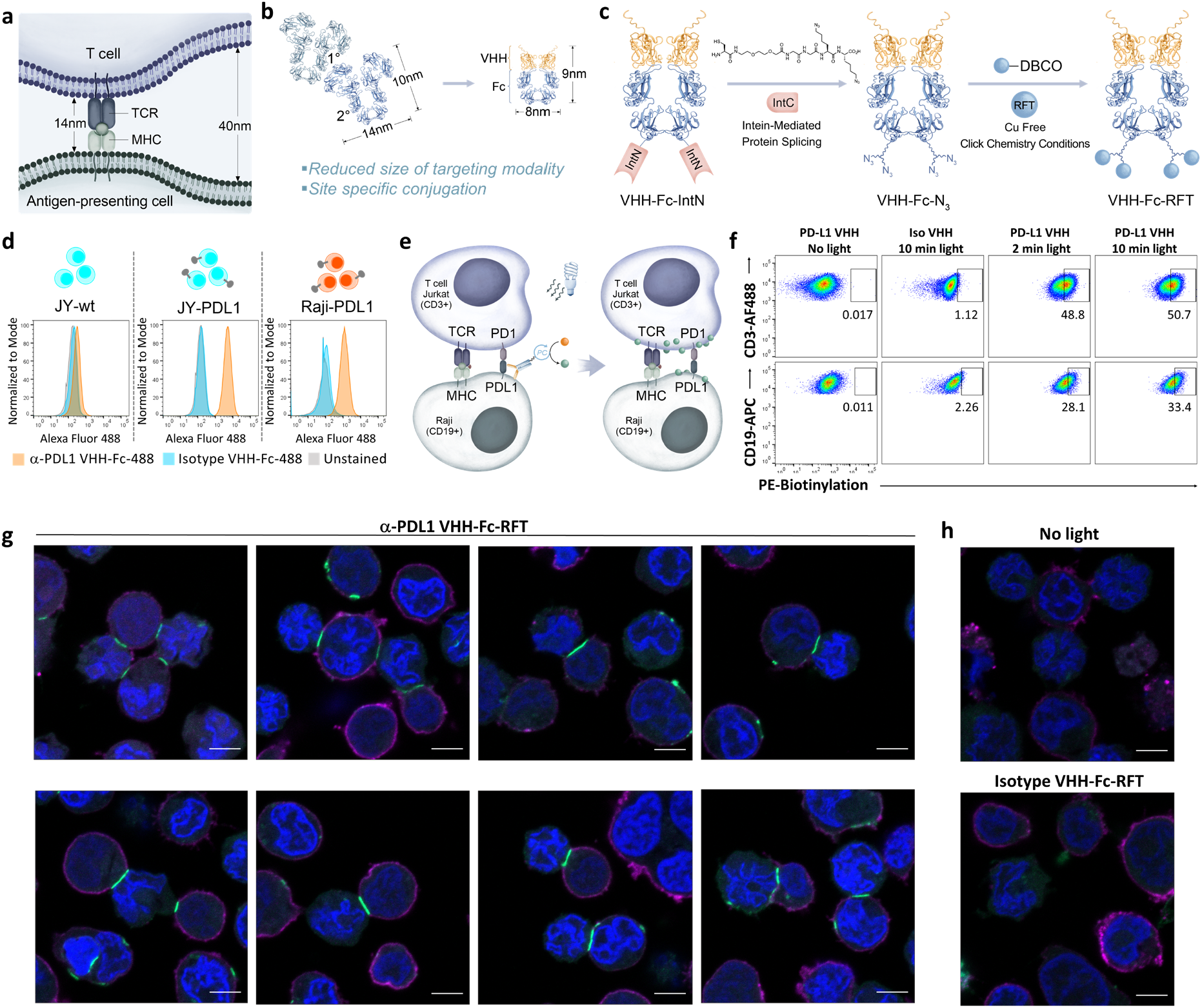
Selective biotinylation within cellular immune synapses. a) Schematic of the immune synapse highlighting the tight intermembrane distance between T cell and antigen-presenting cell. b) Size and dimensionality comparison between the primary/secondary antibody and VHH-Fc PhoTag systems. c) VHH is expressed as an Fc-Intein fusion for C-terminal attachment of the azido linker via intein-mediated ligation to form the VHH-Fc-N_3_ homodimer. RFT-DBCO is covalently attached to the VHH-Fc at the azido linkers to generate VHH-Fc-RFT. d) Flow cytometry analysis shows cell-surface binding of PD-L1 VHH-Fc (α-PD-L1 VHH-Fc labeled with 488 dye (α-PD-L1 VHH-Fc-488)) only on PD-L1-expressing cells. A non-binding control Isotype VHH (Isotype VHH-Fc-488) does not bind the cell surface. e) Schematic showing VHH-Fc-targeted labeling with the RFT photocatalyst (VHH-Fc-PhoTag) on PD-L1 within a Jurkat-Raji two-cell system. f) Flow cytometry analysis of the Jurkat-Raji co-culture system incubated with α-PD-L1 VHH-Fc-RFT (PD-L1 VHH) shows biotinylation on both Raji and Jurkat cells at 2 min and 10 min of light irradiation, but not when targeting with a non-binding Isotype VHH-Fc-RFT (Iso VHH) control construct or in the absence of visible light. g) Confocal imaging of the Jurkat-Raji two cell system incubated with α-PD-L1 VHH-Fc-RFT followed by light irradiation for 2 min results in highly selective biotinylation of the immune synapse whereas no biotinylation occurs in h) the absence of light or with isotype control targeting constructs. Cells were imaged for biotinylation (green), CD3 surface expression (magenta), and nuclei (Hoechst stain, blue). Scale bars indicate 5 μm. Eight replicate images for PD-L1 targeted labeling are shown.

We hypothesized that a single domain antibody (VHH) Fc domain (Fc) fusion construct that is directly labeled with the RFT photocatalyst (VHH-Fc-PhoTag) would overcome steric constraints imposed by the cell synapse. This VHH-Fc-PhoTag system would retain the principal design of the Ab-PhoTag system, but possess a 4-fold reduction in molecular weight and associated decrease in hydrodynamic radius (**Figure 4b**)^42^. To generate the VHH tool to profile the immune synapse, we took a design approach involving a non-blocking α-PD-L1 VHH that would provide minimal disruption of the PD-1/-PD-L1 binding interaction. Accordingly, an alpaca was immunized with human PD-L1 to develop a PD-L1 VHH phage library and the library was screened to find a PD-L1 binding VHH that does not disrupt the PD-1/PD-L1 interaction (**Supporting Information, VHH Generation**). This PD-L1 VHH was then genetically fused at the N-terminus to a human IgG Fc to generate a dimeric form of the VHH. A bis-azide linker was attached site-specifically to the C-terminus of the Fc via intein-mediated protein ligation and subsequently conjugated to the RFT photocatalyst via click chemistry to yield α-PD-L1 VHH-Fc-RFT (**Figure 4c**)^43,44^. Importantly, this VHH-Fc conjugate was confirmed to bind PD-L1 at the cell surface but not block the PD-1/PD-L1 interaction (**Figure 4d and Supporting Figure 20**).

To test this VHH-Fc-PhoTag modality, α-PD-L1 VHH-Fc-RFT was used to label the synapse between the Jurkat PD-1/Raji PD-L1 two cell system to achieve transcellular biotinylation (**Figures 4e and 4f**). In negative control experiments, targeted labeling on cells missing complementary receptors to the Raji cells (i.e. TCR^-^/PD-1^-^ Jurkat cells and A375 melanoma cells) resulted in minimal to no transcellular labeling (**Supporting Figure 21**). Similarly, transcellular tagging of Jurkat PD-1 cells was reduced in the presence of a PD-L1 blocking antibody that was added to Raji PD-L1 cells prior to performing VHH-Fc-PhoTag (**Supporting Figure 22**). Confocal imaging revealed highly selective and exquisite synaptic biotinylation between regions of cellular contact including the formation of multiple synapses from the same cell surface (**Figures 4g and 4h**). This selective synaptic biotinylation effect was also observed by confocal microscopy in two additional two cell systems (Jurkat PD-1/JY PD-L1 and Jurkat PD-1/CHO PD-L1) (**Supporting Figure 23**). In contrast, using a α-PD-L1 VHH-Fc-peroxidase conjugate results in non-selective surface labeling of both Jurkat PD-1 and Raji PD-L1 cell types (**Supporting Figure 24**). Increasing the size of the VHH-Fc labeling system by including a secondary antibody flavin conjugate increased the degree of extrasynaptic *cis*-labeling to similar levels as the primary/secondary Ab-PhoTag system confirming the importance of size reduction to achieve selective synaptic labeling (**Supporting Figure 24**). Collectively, these results highlight the remarkable precision by which the VHH-Fc-PhoTag system directs chemical tags into the immune synapse.

### Differential Analysis of Synapse-Forming Versus Bystander T cells

The selective tagging ability of the VHH-Fc-PhoTag system enables the downstream differential characterization of interacting cells. Accordingly, we investigated cell interactions between Raji PD-L1 and Jurkat PD-1 cells in the presence of SED superantigen using high versus (vs.) low transcellular biotinylation to uncover genetic differences unique to interacting vs. non-interacting, bystander cells (**Figure 5a**). After applying VHH-Fc-PhoTag on PD-L1 within the Jurkat-Raji co-culture system, Jurkat PD-1 cells were sorted into low or high biotinylation groups (**Figures 5b and 5c**). This was followed by RNA-seq analysis which clearly revealed unique genetic differences between these two isolated cell populations (**Supporting Figure 25**). Furthermore, for high vs. low biotin differentially expressed genes, expression profiling indicates that bystander (biotin low) Jurkat cell population resembled that of resting Jurkat cells, while the interacting (biotin high) Jurkat cells more closely matched the bulk gene expression pattern of the Jurkat + Raji co-culture system (**Figure 5d and Supplementary Figure 25**). Differential gene expression analysis resulted in the identification of 75 genes that were altered between biotin high and low cell populations and were primarily comprised of immune related annotations within the innate and adaptive immune response as well as in cell adhesion and cell recognition (**Figure 5e and Supplementary Figure 25**). The genetic differences identified between interacting and bystander cells ultimately highlight the role of MHC Class II-TCR engagement in driving the B and T cell interaction. Moreover, this experiment demonstrates the power of the VHH-Fc-PhoTag system to characterize differences between cellular interactors and bystanders in meaningful biological systems through transcellular tagging at the immune synapse.

**Figure 5.**
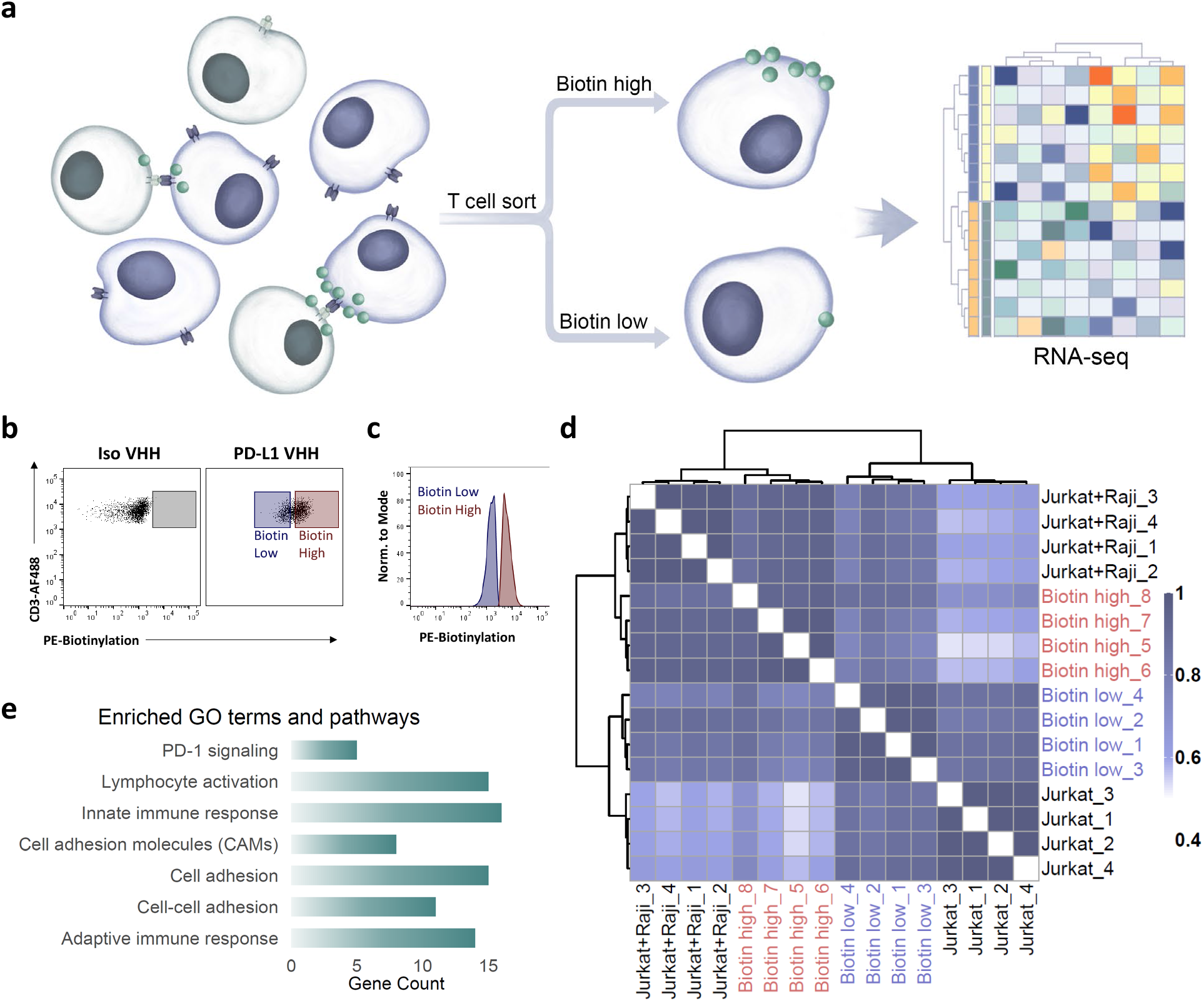
RNA-seq analysis of high vs. low biotinylated cells. a) Schematic of workflow to sort high vs. low biotinylated cells in the Jurkat-Raji co-culture system after using VHH-Fc-RFT-PhoTag for comparative RNA-seq analysis (Jurkat cells, blue; Raji cells, gray). b) Flow cytometry analysis of Jurkat cell transbiotinylation from Isotype VHH-Fc-RFT (Iso VHH) or α-PD-L1 VHH-Fc-RFT (PD-L1 VHH) targeted labeling. The isotype gating (gray box) was used to set biotin high gate (red box). Blue box indicates cells designated as biotin low. c) Gating strategy for cell sorting shows sample separation into distinct biotin high and biotin low populations. d) Correlation plot of biotin high and biotin low transcriptomic profiles for differentially expressed genes (absolute log2 FC >0.5, FDR <0.05) derived from a single experiment of 16 samples (n=4 in each group). Pearson correlation is denoted in the color key. Hierarchical clustering was performed with the dissimilarity set as 1-Pearson correlation and linkage criteria set as complete. e) Significantly enriched (FDR<0.05) GO and pathway terms of interest. Bars represent the gene counts within the significantly expressed gene list (FDR<0.05) present in each GO or pathway term.

## Discussion

Here we introduced PhoTag as a novel, non-enzymatic cell tagging technology. We demonstrated selective covalent labeling of antigen-specific T cells isolated from whole blood and captured uniquely interacting cell-cell interactions. The PhoTag method presented here leverages the formation of tyrosyl radicals to tag interacting cells through the use of visible light activation of a flavin cofactor-derived photocatalyst. Unlike other enzyme-based cell tagging methods^8-11^, the photoactivatable nature of this flavin system allows for exquisite control through easily tunable parameters such as light intensity and/or duration. Indeed, using the PhoTag method we observed time-dependent cell surface tagging within minutes of light exposure allowing for the ability to capture snapshots of biological processes that include the migration of PD-L1 into the synapse. PhoTag can be broadly utilized to profile cell surface interactions in a targeting modality-agnostic manner and without the need for additional cell manipulation through genetic engineering. We anticipate that this work will find wide applicability for exploring immunological cell interactions as well as profiling other biologically relevant intercellular junctions both to better understand their complex basic biology and to design rational therapeutic strategies that manipulate critical cell-cell interactions for positive clinical outcomes.

## Acknowledgements

We are very grateful to Nick Haining, Scott Lesley, Kalpit Vora, Alberto Visintin, and Tao Wang from Merck & Co., Inc. for helpful discussions.

## Author contributions

O.O.F., R.C.O., and T.R.R. conceived of the work. O.O.F., R.C.O., T.R.R., C.H.W., J.H.T., K.A.C., L.L., D.C., V.M.P. designed and executed experiments. D.H.P., S.D.O. interpreted results. S.F., M.V.-P. designed and engineered the VHH-Fc conjugates. E.P.B., D.V., and K.C. screened and identified the α-PD-L1 VHH. E.C.H., L.R.R., G.P., and D.J.H. provided insight and direction for experimental design. O.O.F., R.C.O., T.R.R., and C.H.W. wrote the manuscript with input from all authors.

