## Supporting Information for "Detection of Cell-Cell Interactions via Photocatalytic Cell Tagging"

### Supporting Figures and Tables

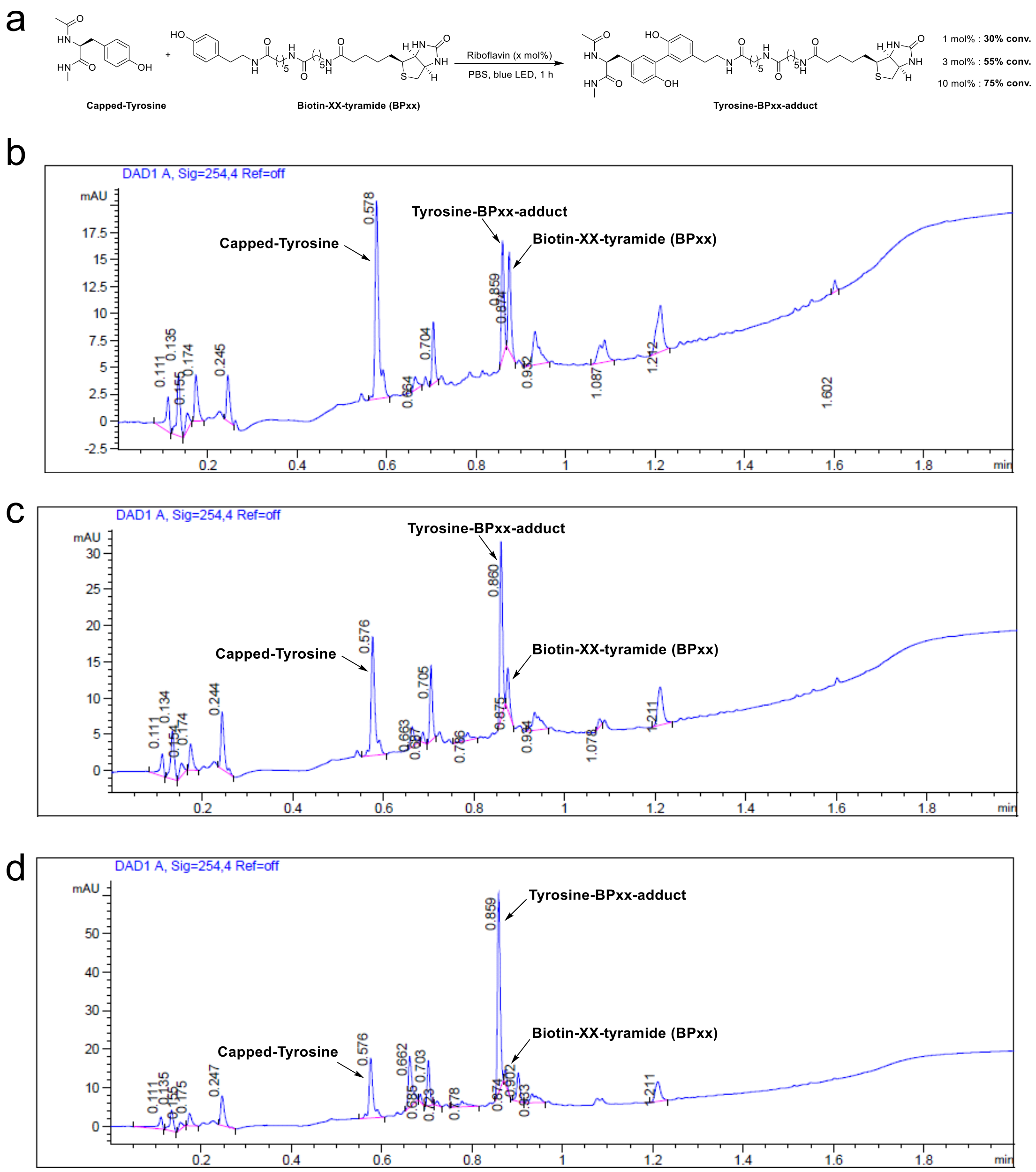

**Supporting Figure 1. Flavin-based photocatalytic mediated cross-coupling of biotin-xx-tyramide and capped tyrosine**. a) Schematic depicting model reaction of biotin phenol (biotin-xx-tyramide) and capped tyrosine in the presence of riboflavin photocatalyst under blue LED illumination (conversion calculated from UPLC areas analysis). b) LC-MS trace of cross-coupling of biotin-xx-tyramide and capped tyrosine using 1 mol% riboflavin photocatalyst. c) LC-MS trace of cross-coupling of biotin-xx-tyramide and capped tyrosine using 3 mol% riboflavin photocatalyst. d) LC-MS trace of cross-coupling of biotin-xx-tyramide and capped tyrosine using 10 mol% riboflavin photocatalyst.

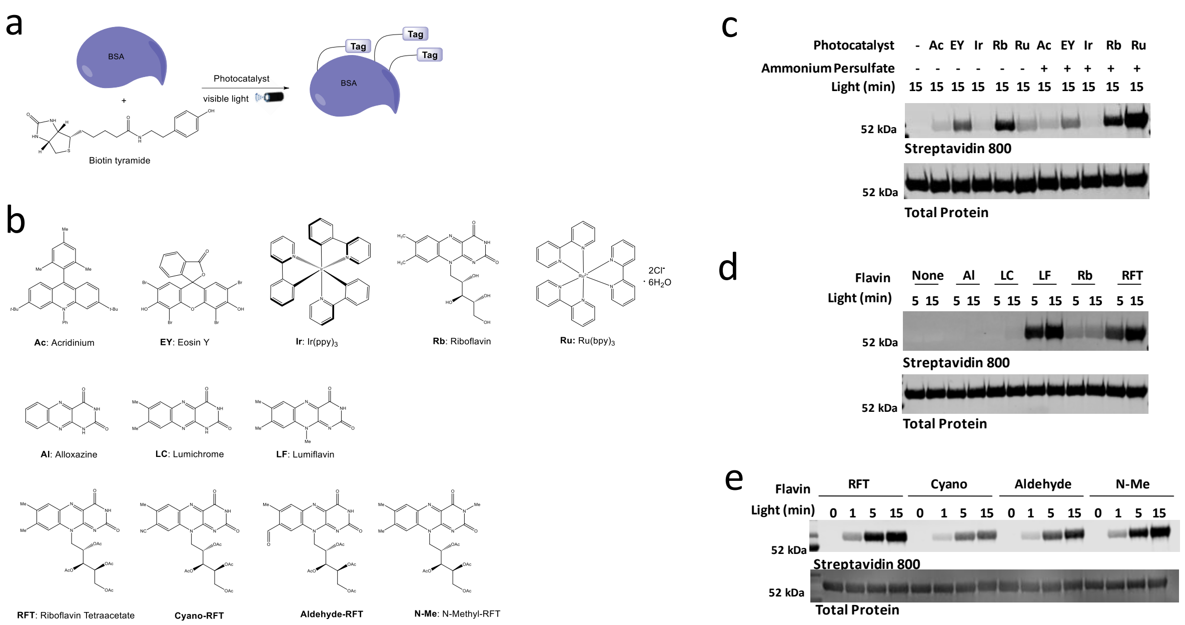

**Supporting Figure 2. Photocatalyst screen for BSA biotinylation**. a) Schematic depicting the protein labeling reaction with photocatalyst, biotin tyramide, and BSA. b) Photocatalysts tested in this work. c) Western blot analysis of BSA (1mg/ml) in the presence of biotin tyramide (250µM), different photocatalysts (10µM) and/or ammonium persulfate (500µM). Based on this screen, riboflavin (Rb) displayed the highest level of protein biotinylation compared to the other catalysts. The presence of ammonium persulfate increased biotinylation for riboflavin (Rb) and the ruthenium (Ru) catalyst. d) Different flavin photocatalysts were screened for BSA biotinylation using similar conditions described in panel c. Both Lumiflavin (LC) and riboflavin tetraacetate (RFT) yielded the highest biotinylation levels as measured by western blot. e) Cyano_RFT,[1] formyl-RFT, and *N*-methyl-RFT[2] (1µM) were screened for BSA biotinylation without increased labeling over RFT.

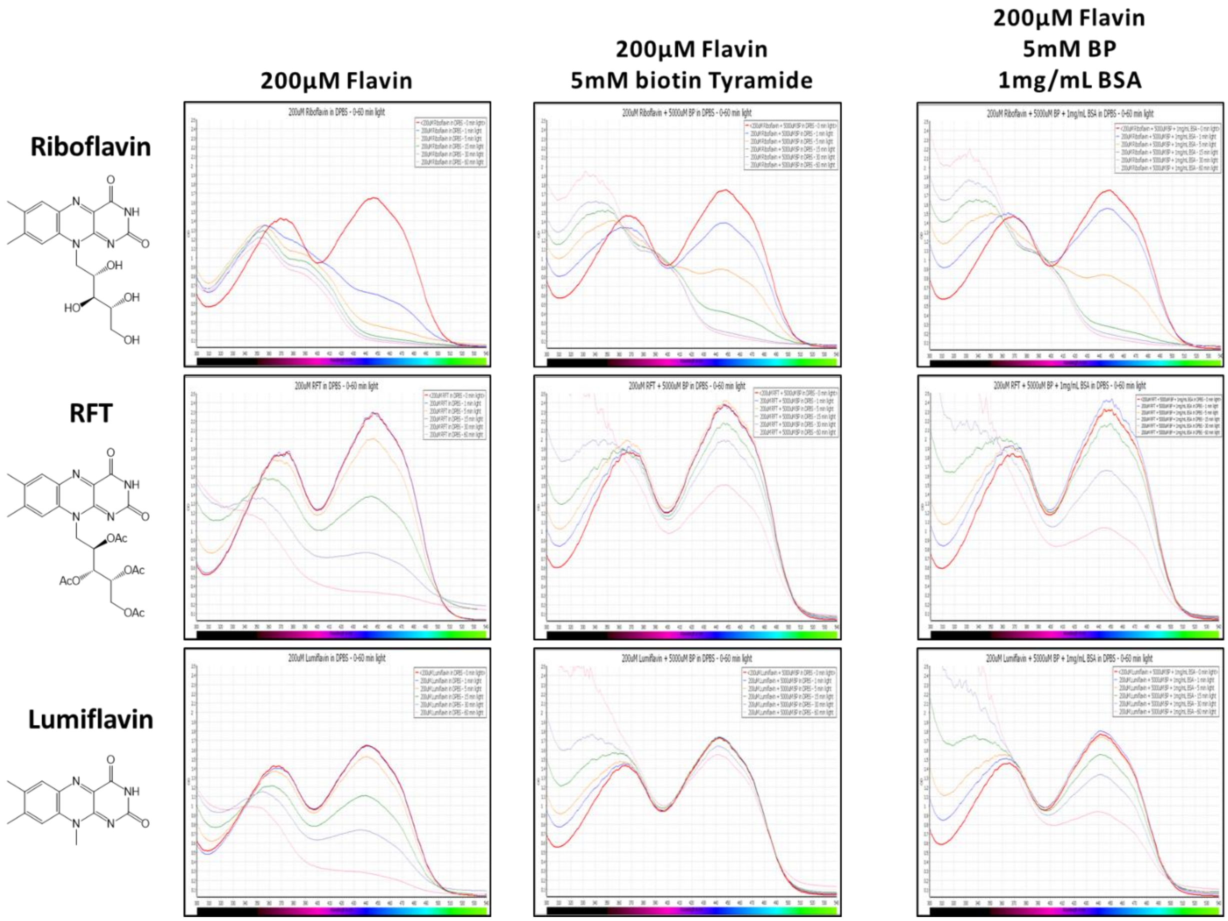

**Supporting Figure 3. Analysis of flavin catalyst photostability by UV-Vis spectroscopy.** UV-Vis absorption spectra (300-550 nm) for riboflavin, RFT, and lumiflavin in the presence or absence of 5mM biotin tyramide or 5mM biotin tyramide and 1mg/ml BSA were measured over a time course of visible light exposure in the bio-photoreactor (0 min (red line), 1 min (blue line), 5 min (yellow line), 15 min (green line), 30 min (purple line), or 60 min (pink line). The absorption band of both RFT and LF shows photostability over Rb. Further photostability of flavin catalysts in the presence of biotin tyramide and BSA confirms the catalytic turnover of the flavins.

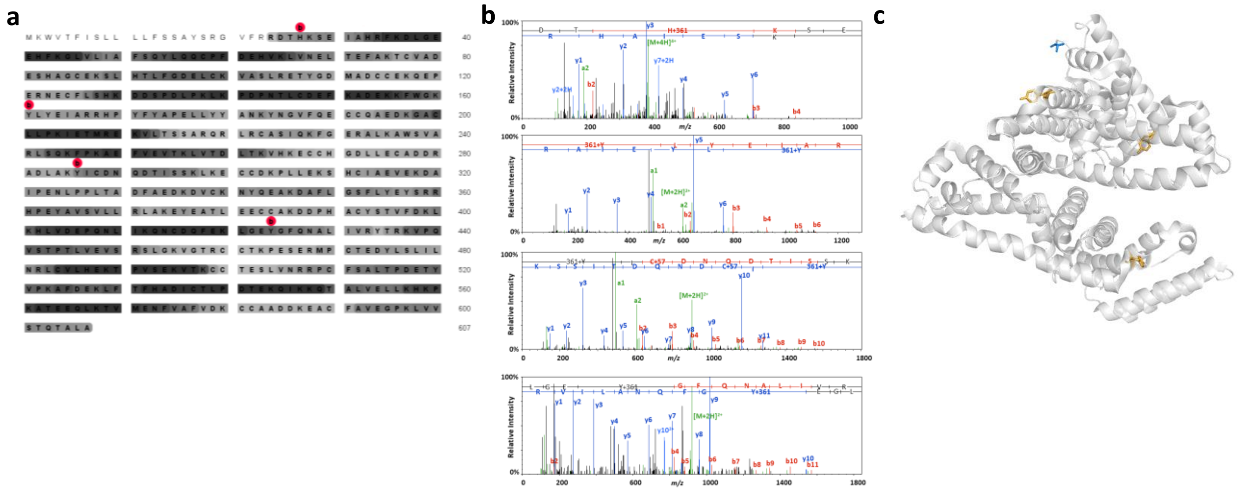

**Supporting Figure 4. LC-MS/MS analysis of BSA biotinylation.** a) BSA protein sequence coverage map with detected residues bearing the biotinylation modification indicated with a red dot. b) Peptide spectral matches for peptides bearing the biotinylation modification for each site identified on BSA. Assignments of theoretical b- and y-type fragments and parent ion species are indicated on the empirical MS/MS spectra. c) Crystal structure of BSA (PDB code: 3v03) with each detected site of biotinylation indicated on tyrosine (orange) or histidine (blue).

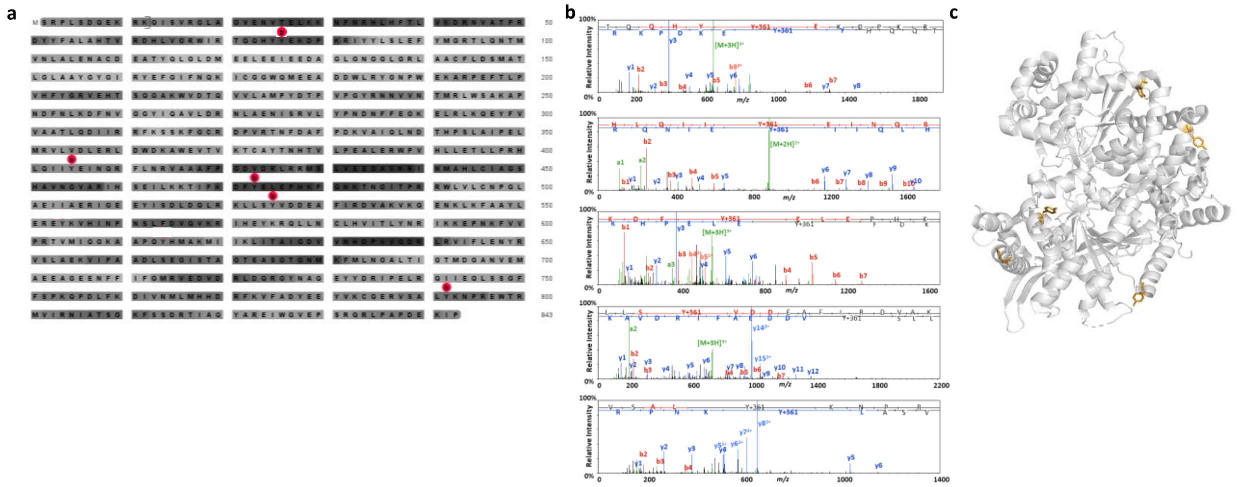

**Supporting Figure 5. LC-MS/MS analysis of Phosphorylase B biotinylation.** a) Phosphorylase B protein sequence coverage map with detected residues bearing the biotinylation modification indicated with a red dot. b) Peptide spectral matches for peptides bearing the biotinylation modification for each site identified on phosphorylase B. Assignments of theoretical b- and y-type fragments and parent ion species are indicated on the empirical MS/MS spectra. c) Crystal structure of Phosphorylase B (PDB code: 2qn2) with each detected site of biotinylation indicated on tyrosine (orange).

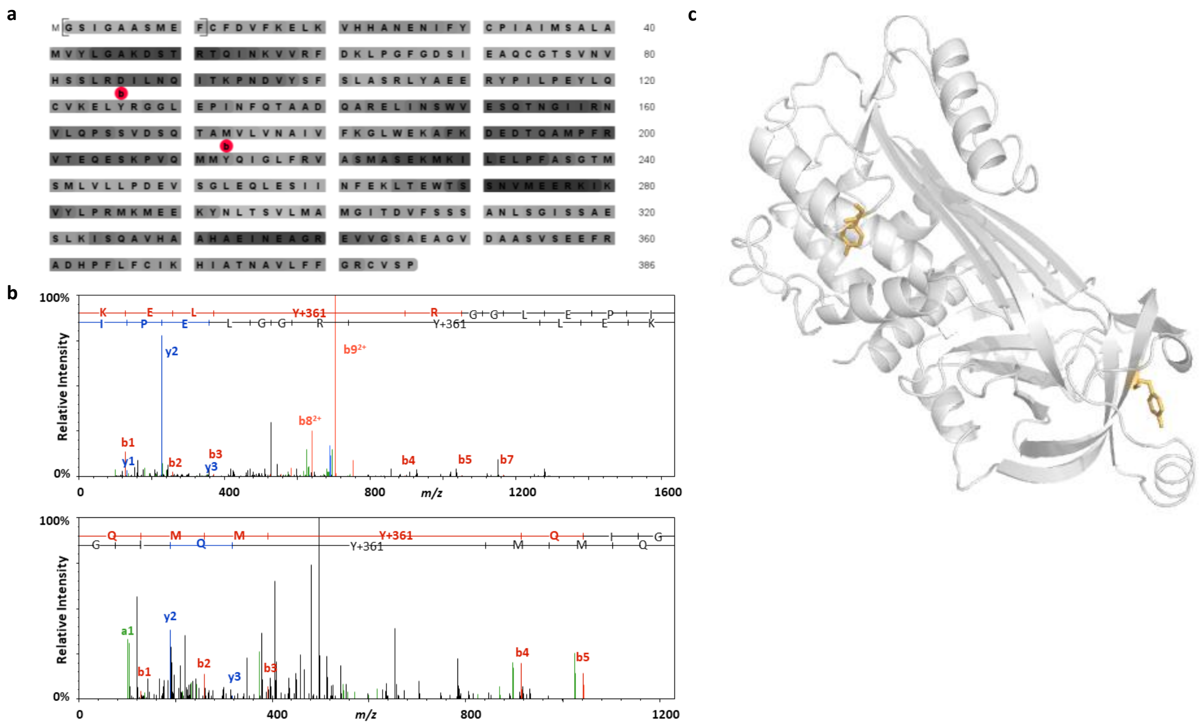

**Supporting Figure 6. LC-MS/MS analysis of Ovalbumin biotinylation.** a) Ovalbumin protein sequence coverage map with detected residues bearing the biotinylation modification indicated with a red dot. b) Peptide spectral matches for peptides bearing the biotinylation modification for each site identified on ovalbumin. Assignments of theoretical b- and y-type fragment ion species are indicated on the empirical MS/MS spectra. c) Crystal structure of ovalbumin (PDB code: 1ova) with each detected site of biotinylation indicated on tyrosine (orange).

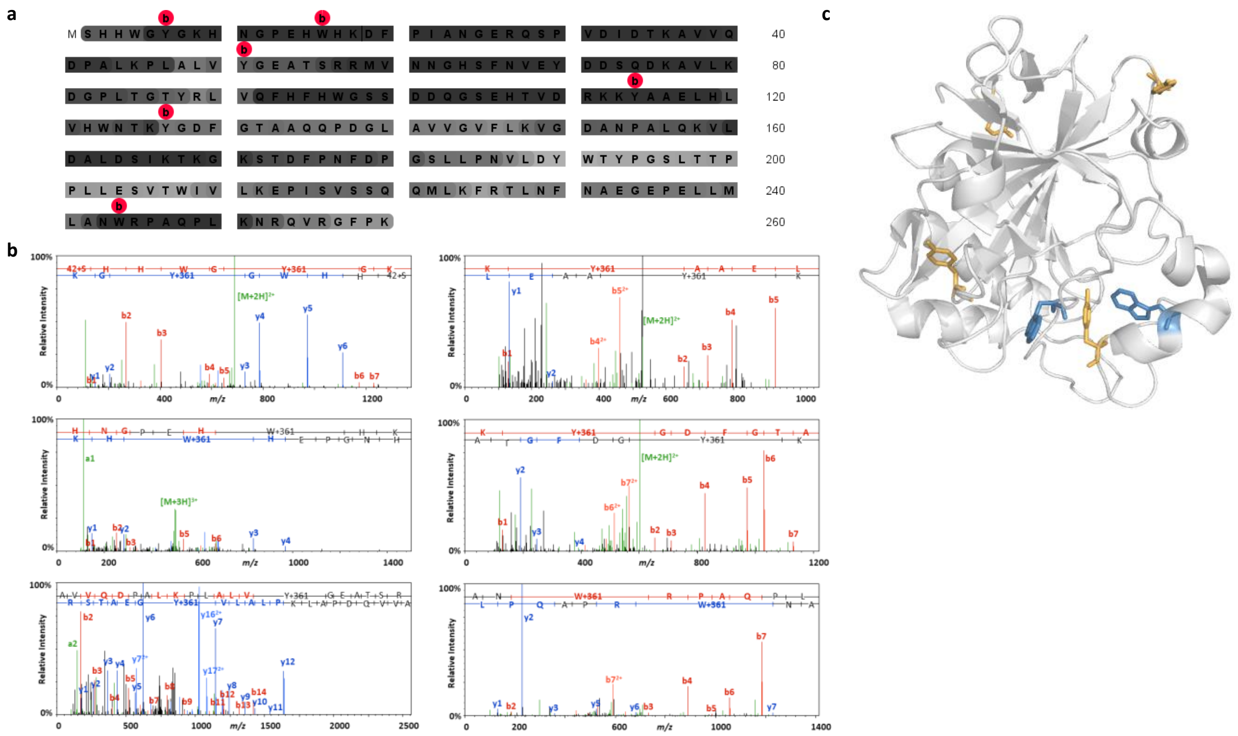

**Supporting Figure 7. LC-MS/MS Analysis of Carbonic Anhydrase biotinylation.** a) Carbonic anhydrase protein sequence coverage map with detected residues bearing the biotinylation modification indicated with a red dot. b) Peptide spectral matches for peptides bearing the biotinylation modification for each site identified on carbonic anhydrase. Assignments of theoretical b- and y-type fragments and parent ion species are indicated on the empirical MS/MS spectra. c) Crystal structure of carbonic anhydrase (PDB code: 1v9e) with each detected site of biotinylation indicated on tyrosine (orange) or tryptophan (blue).

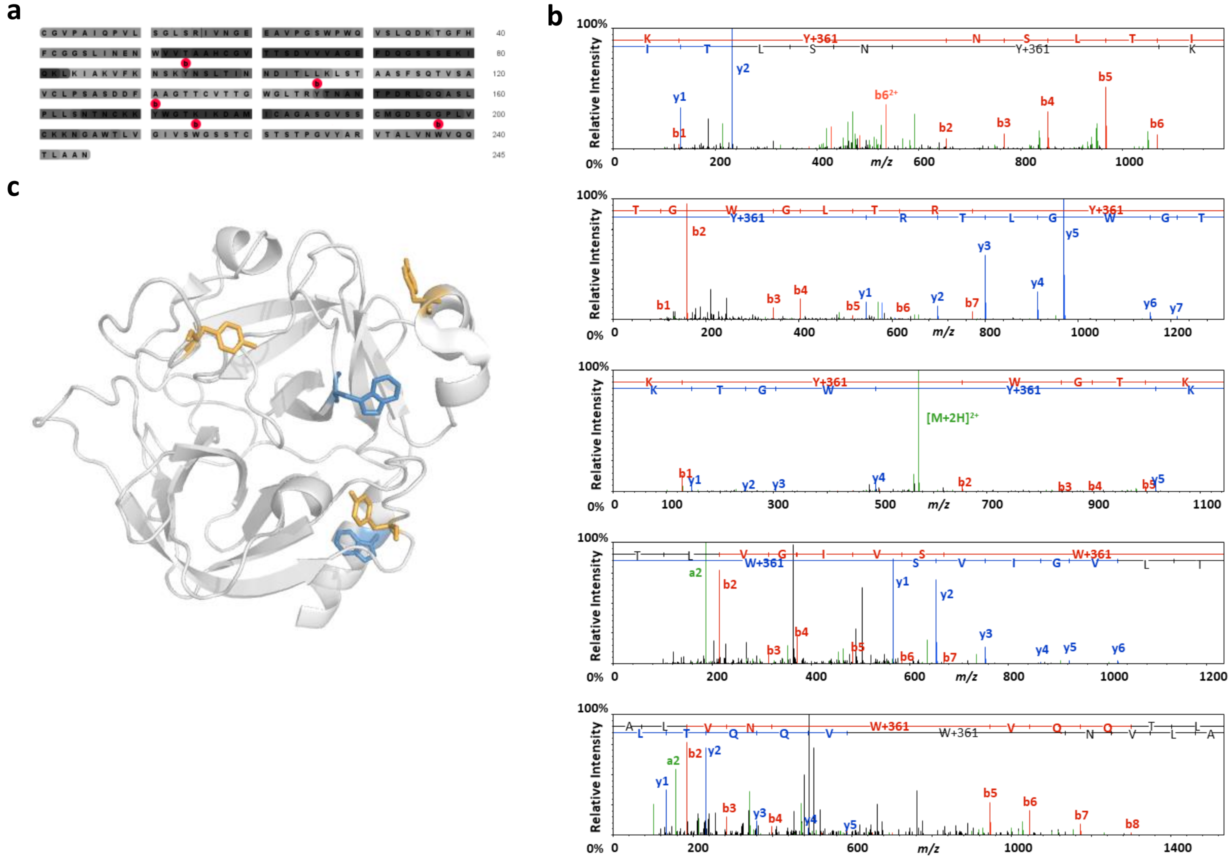

**Supporting Figure 8. LC-MS/MS analysis of Chymotrypsinogen A.** a) Chymotrypsinogen A protein sequence coverage map with detected residues bearing the biotinylation modification indicated with a red dot. b) Peptide spectral matches for peptides bearing the biotinylation modification for each site identified on Chymotrypsinogen A. Assignments of theoretical b- and y-type fragments and parent ion species are indicated on the empirical MS/MS spectra. c) Crystal structure of Chymotrypsinogen (PDB code: 2cga) with each detected site of biotinylation indicated on tyrosine (orange) or tryptophan (blue).

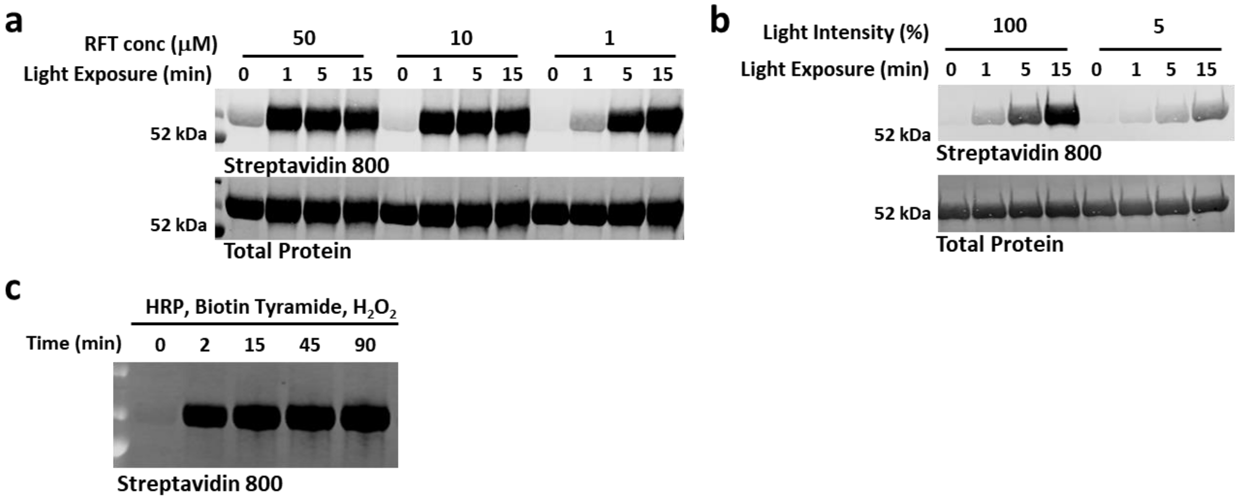

**Supporting Figure 9. Effect of light intensity and catalyst concentration on BSA biotinylation at various time points.** a) RFT-induced BSA biotinylation (1mg/ml BSA, 250µM biotin tyramide) is dependent on light exposure time and catalyst concentration as monitored by western blot. b) The extent of BSA biotinylation over time (1mg/ml BSA, 1µM RFT, 250µM biotin tyramide) can also be controlled by light intensity. c) Western Blot analysis of BSA biotinylation by horse radish peroxidase (HRP) (1mg/ml BSA, 500µM biotin tyramide, 1µM $H_2O_2$, 2.5 µg HRP) shows rapid biotinylation.

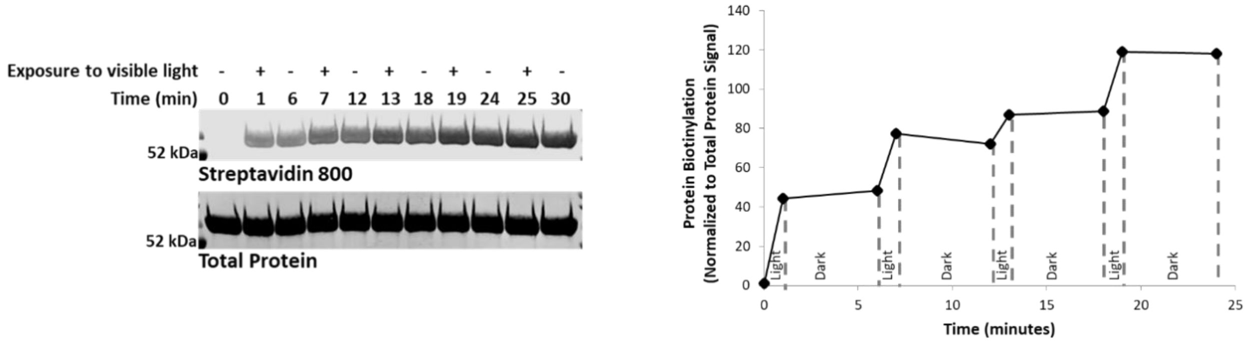

**Supporting Figure 10. Protein Light On/Off Labeling. (**Left Panel) BSA, biotin tyramide, and RFT were exposed to 1 min pulses of visible light irradiation followed by 5 min in the dark and analyzed by western blot at each indicated time point. (Right Panel) Densitometry measurement of the BSA biotinylation signal in panel c indicates that labeling increases with each subsequent pulse of visible light irradiation (light/dark pulse indicated with gray dashed line).

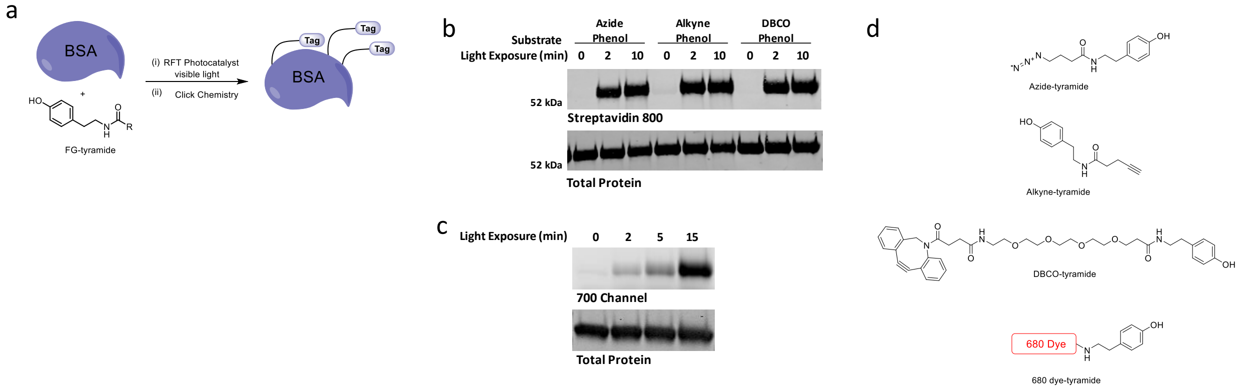

**Supporting Figure 11. Photo-induced protein labeling with functional groups relevant to other bioorthogonal reactions.** a) Reaction scheme depicting protein labeling with different phenol containing probes. b) BSA (1mg/ml) was labeled with azide-[3], alkyne-, or DBCO-tyramide (250µM each) in the presence of RFT (10µM) and visible light irradiation for the indicated time points, followed by attachment of biotin via click reaction. Western blot analysis shows time dependent labeling of each phenol probe. c) Time dependent labeling of α-chymotrypsinogen A (1mg/ml) with 680 dye tyramide fluorophore (250µM) occurred in the presence of RFT (10µM) and visible light irradiation as measured by gel fluorescence imaging. d) Chemical structures of the tyramide probes.

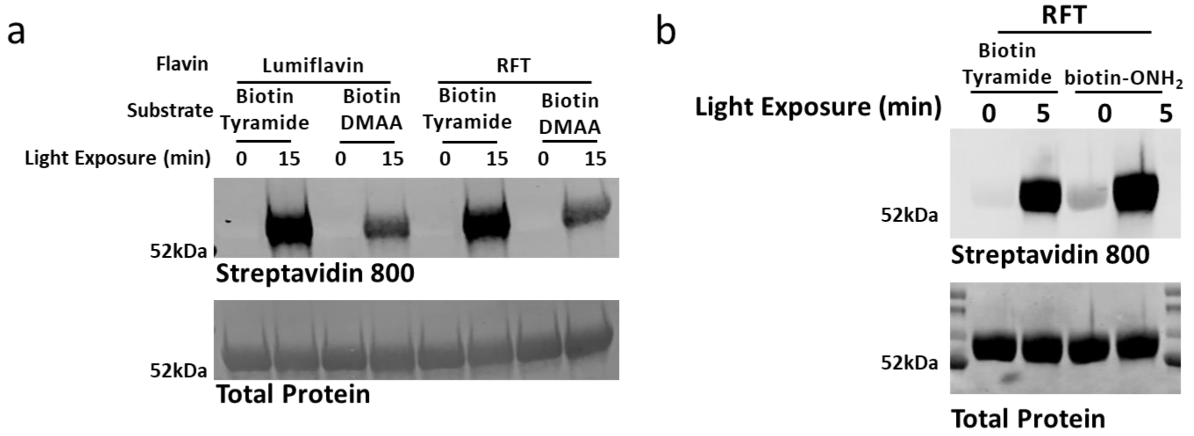

**Supporting Figure 12. Trapping of tyrosyl radicals or tyrosyl oxidative species with biotin probes.** Considering the possible direct flavin activation of protein tyrosine residues, we employed the use of biotin-dimethyl amine aniline (biotin-DMAA) as a radical cross-coupling partner to trap tyrosyl radicals[4]. Accordingly, BSA (1mg/ml), RFT or Lumiflavin (10µM) and 250µM of biotin tyramide or biotin-DMAA were combined and irradiated with visible light for 15 min, followed by western blot analysis. Tyrosine activation for subsequent trapping with biotin-DMAA resulted in decreased biotinylation levels on BSA (panel a), suggesting that labeling of protein tyrosines through flavin-mediated phenoxy radical generation of the added substrate is the more efficient covalent labeling approach. Furthermore, we found that light induced protein oxidation via flavin activation (BSA (1mg/ml), RFT (1µM) and 250µM of biotin tyramide or biotin-ONH$_2$ and irradiated with visible light for 5 min, followed by western blot analysis) can be trapped through use of a strong nucleophile (e.g. amino oxy biotin) albeit with higher background.

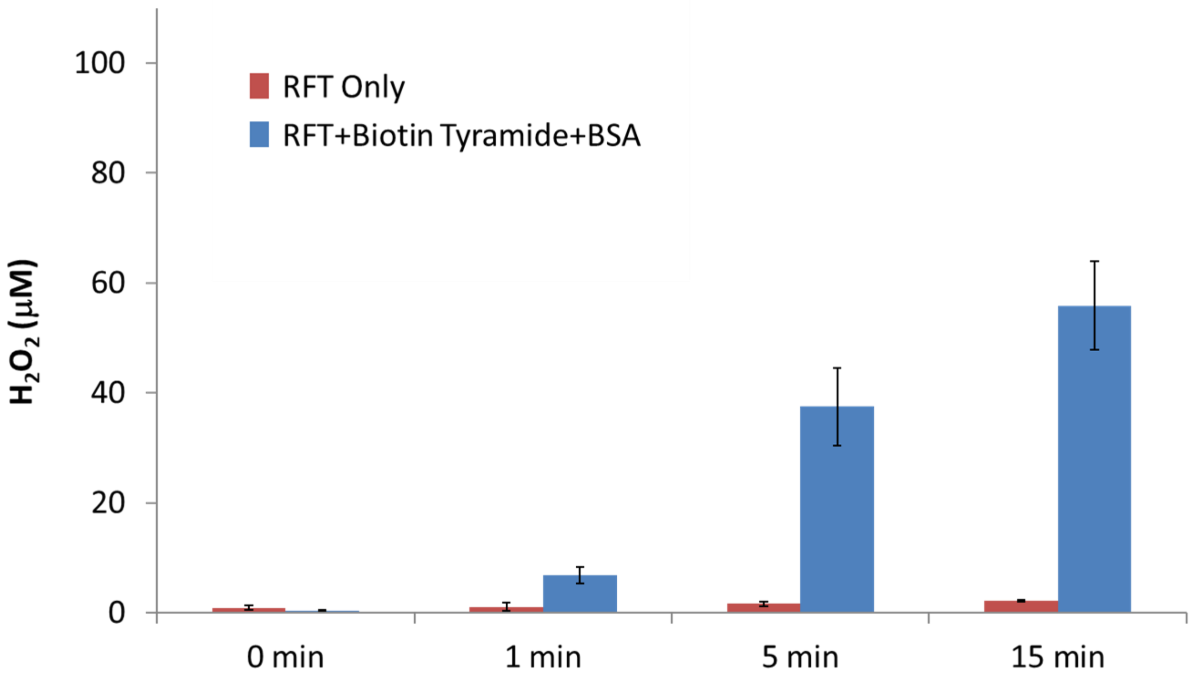

**Supporting Figure 13. Substrate scope and hydrogen peroxide generation during the RFT catalytic cycle.** BSA (1mg/ml), biotin tyramide (250μM), and 1μM RFT or RFT alone were irradiated with visible light for the indicated time points. Hydrogen peroxide generation was monitored by fluorometric analysis (Sigma-Aldrich: MAK165) and primarily detected only in the presence of biotin tyramide and BSA, indicating oxygen is required as the sacrificial electron acceptor to re-oxidize the reduced flavin (See Figure 2c Mechanism) to regenerate the flavin photocatalyst. Additionally, the high micromolar levels of generated $H_2O_2$ from 1μM RFT provide evidence for efficient catalytic turnover within this photolabeling flavin system (error bars represent the average ± s.d. of n = 3 experiments).

a

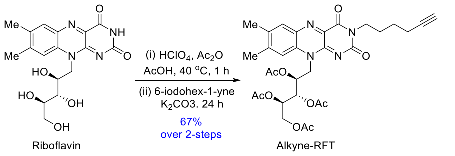

b

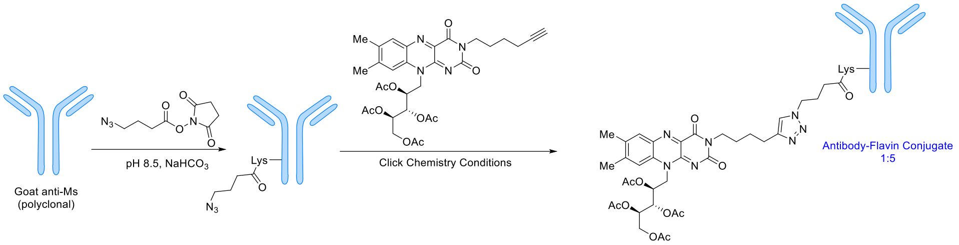

**Supporting Figure 14. Reaction scheme for antibody flavin conjugate preparation.** a) Synthetic route to alkyne-RFT from riboflavin. b) The secondary antibody-flavin conjugate (AFC) was prepared by non-selective labeling of lysine residues with azido NHS ester, followed by click reaction with an alkyne RFT to yield an approximate 1:5 ratio of antibody to flavin.

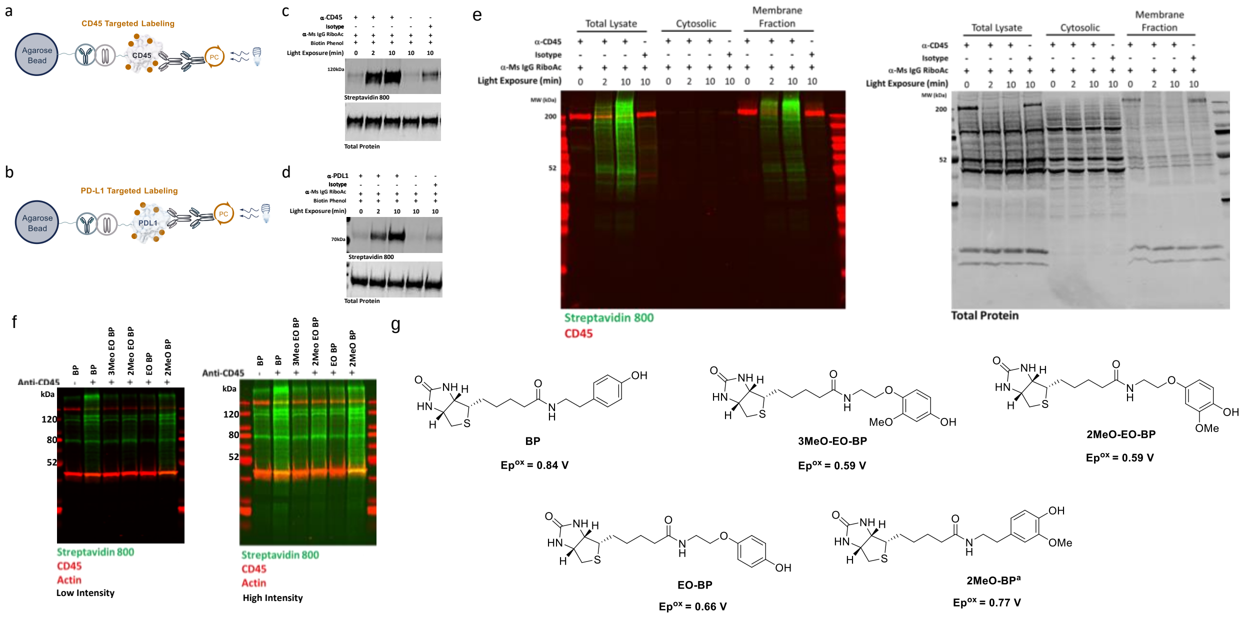

**Supporting Figure 15. Targeted labeling of CD45 or PD-L1.** Schematic depicting bead-based protein labeling of a) CD45-Fc or b) PD-L1-Fc fusion proteins bound to anti-Fc agarose beads followed by attachment of a primary antibody and secondary antibody flavin conjugate. CD45 or PD-L1 proteins are then labeled in the presence of biotin tyramide and visible light. Western blot analysis of light-dependent, targeted biotinylation of c) CD45 or d) PD-L1 bound to agarose beads for the indicated time points. Low levels of protein biotinylation are observed in both the secondary AFC (α-Ms IgG RFT) only and isotype only controls. e) Left panel, Jurkat cells were labeled with α-CD45 antibody or isotype control and secondary antibody flavin conjugate, followed by visible light irradiation for 0, 2, or 10 min. After photolabeling, total cell lysate, cytosolic lysate, or enriched membrane lysate was generated and analyzed by western blot for biotinylation and CD45 levels. The absence of biotinylation signal in the cytosolic fraction supports primary biotinylation at the membrane surface. Right panel, total protein stain of left panel image. f) Western blot analysis of CD45-targeted labeling on Jurkat cells using the primary/secondary antibody flavin conjugate. Left and right images display western blot and low and high intensity, respectively. g) The different probe analogs to achieve various degrees of labeling are shown in panel f, with their estimated oxidation potentials, based on previous reports[5-7]. The western blot signal intensities vary among each probe likely due to differences in probe oxidation potential and indicate biotin tyramide (BP) as an optimal probe for labeling. [a]Compound was prepared as described previously[8].

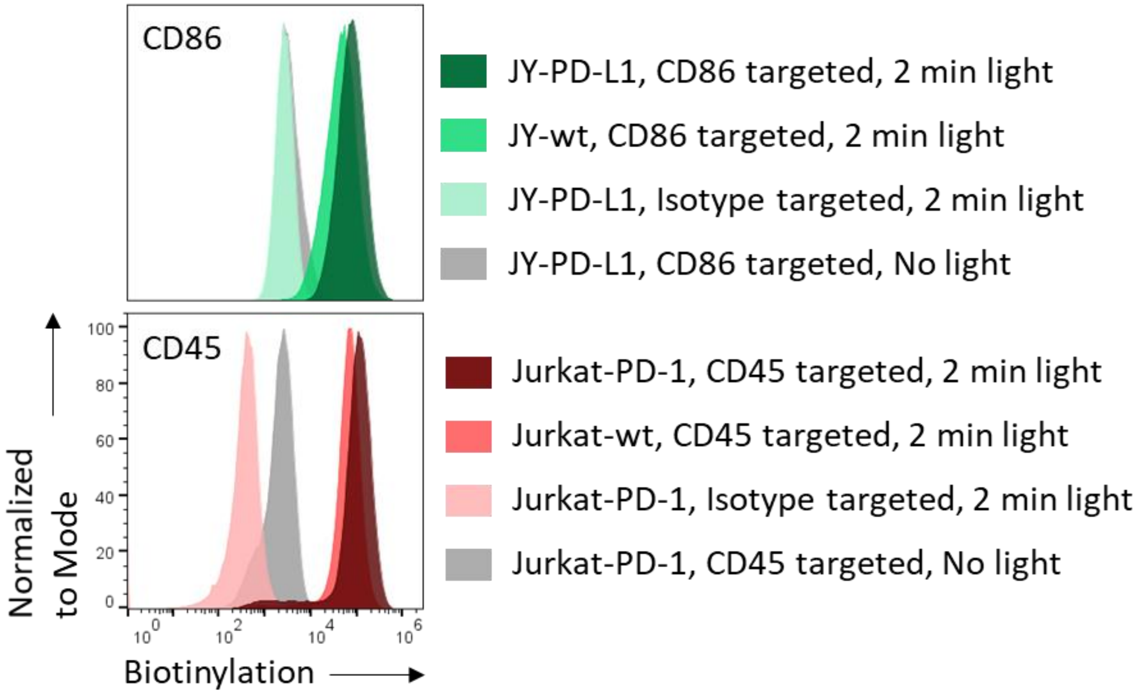

**Supporting Figure 16. Cell surface targeting of CD86 or CD45.** Flow cytometry analysis of targeted labeling of CD86 on JY wt or JY PD-L1 cells or CD45 on Jurkat wt or Jurkat PD-1 cells. Increased surface biotinylation occurs over isotype targeting and only in the presence of visible light.

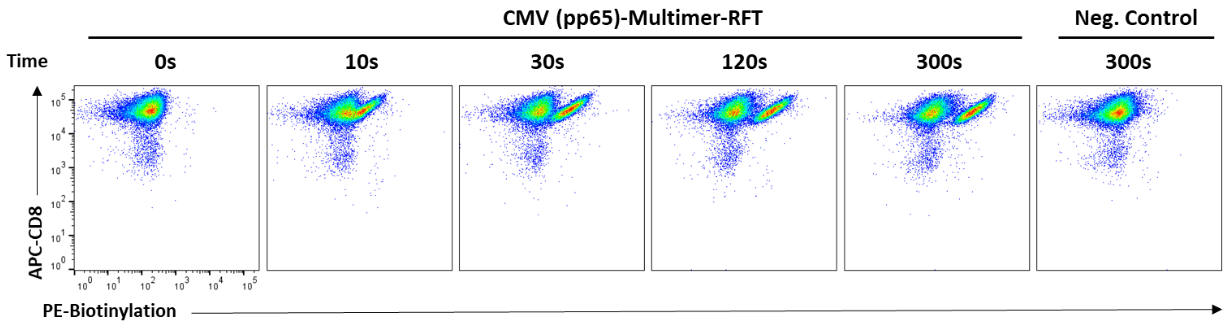

**Supporting Figure 17. Time course of antigen specific biotinylation of CD8+ T cells using PhoTag technology.** CMV-specific CD8+ T cells were incubated with MHC-Multimer-RFT displaying a CMV pp65 peptide or a non-binding negative control. The cells were irradiated with visible light in the presence of biotin tyramide for the indicated time points. A light exposure time-dependent increase in biotinylation of CMV-specific cells was observed by flow cytometry.

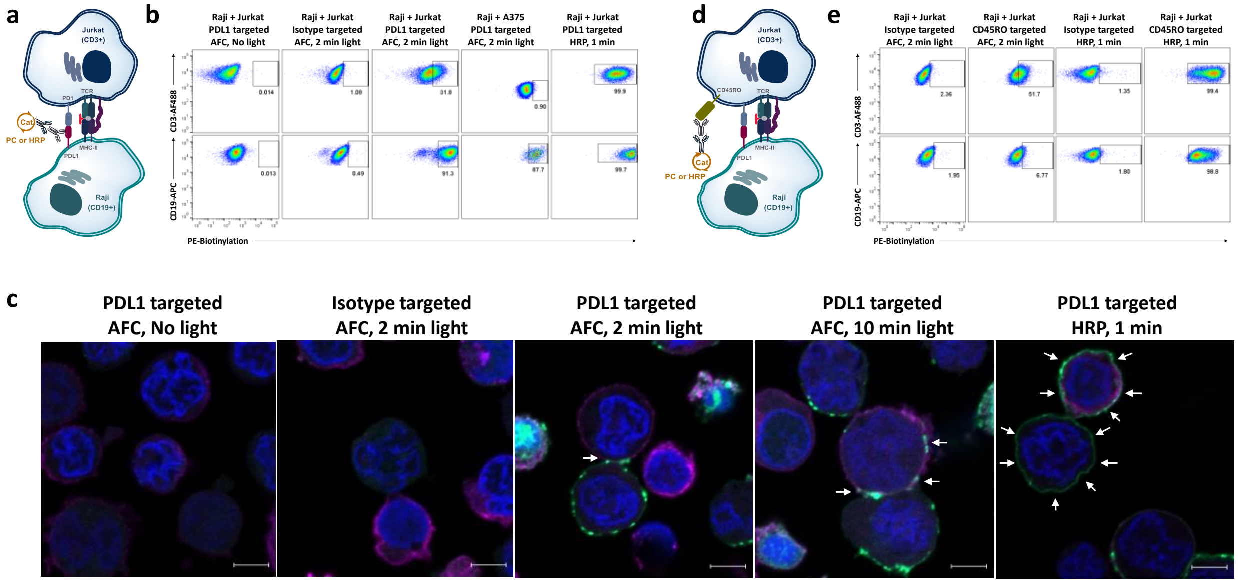

**Supporting Figure 18. Photocatalytic proximity labeling within a two-cell system.** a) Schematic depicting a two-cell system consisting of engineered Jurkat PD-1 and Raji PD-L1 cells. Antibody-targeted labeling with a photocatalyst (PC) or peroxidase (HRP) on PD-L1 is highlighted. b) Biotinylation is detected on both Raji and Jurkat cells with PD-L1 targeting using the antibody flavin conjugate (AFC), but not Isotype targeting or in the absence of visible light irradiation. Transcellular labeling was not observed between PD-L1-labeled Raji cells and suspended A375 cells that do not share complimentary receptors to the Raji PD-L1 cells. c) Confocal microscopy imaging of the Raji-Jurkat two cell system with PD-L1 targeting on Raji cells reveals labeling on both Raji cells and points of cellular contact on Jurkat cells using the AFC while excessive labeling on both cell types was observed with HRP (indicated with white arrows). Cells were imaged for biotinylation (green stain), CD3 surface expression (magenta stain), and nuclei (Hoechst stain). Scale bars indicate 5 µm. d) Schematic depicting a two-cell system consisting of engineered Jurkat and Raji cells. Antibody-targeted labeling with a photocatalyst (PC) or peroxidase (HRP) on CD45 is highlighted. e) Targeting labeling of CD45RO on Jurkat cells (known to be excluded from the synapse) resulted in low levels of Raji transcellular labeling using an AFC and nearly quantitative labeling when HRP was used.

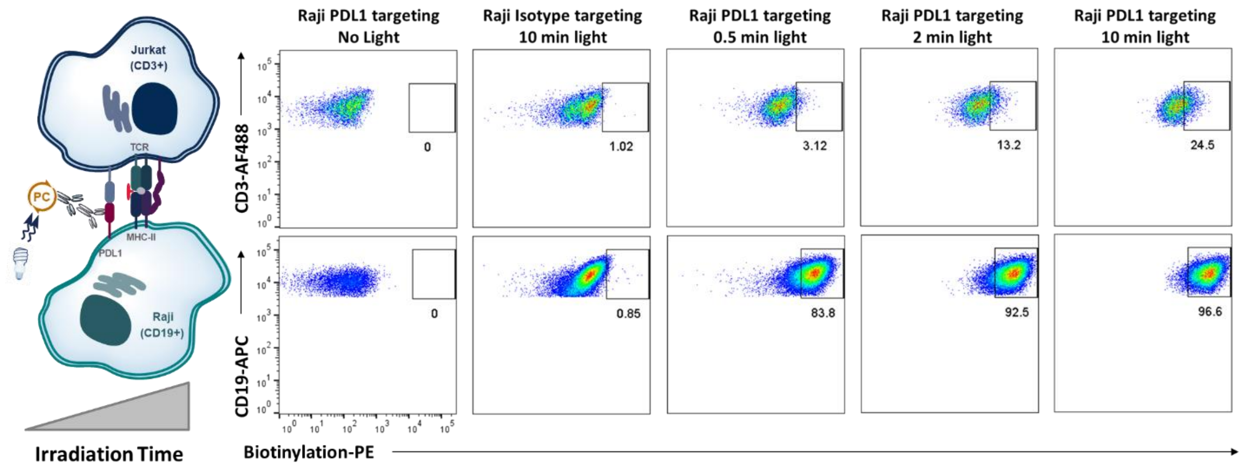

**Supporting Figure 19. Effect of light exposure time on two-cell system labeling with primary/secondary antibody system.** Raji PD-L1 cells (CD19+) were pre-labeled with α-PD-L1 or isotype control and secondary AFC. The cells were then mixed with Jurkat PD-1 cells (CD3+) in the presence of superantigen for 2.5 hours. Biotin tyramide was gently added to the cell mixture and the cells were then irradiated for the indicated time points. Both *cis*-biotinylation on the Raji PD-L1 cells and *trans*-biotinylation on Jurkat PD-1 cells are observed. Note: we observed that *cis*-biotinylation occurs over a shorter visible light irradiation time compared to the *trans*-biotinylation.

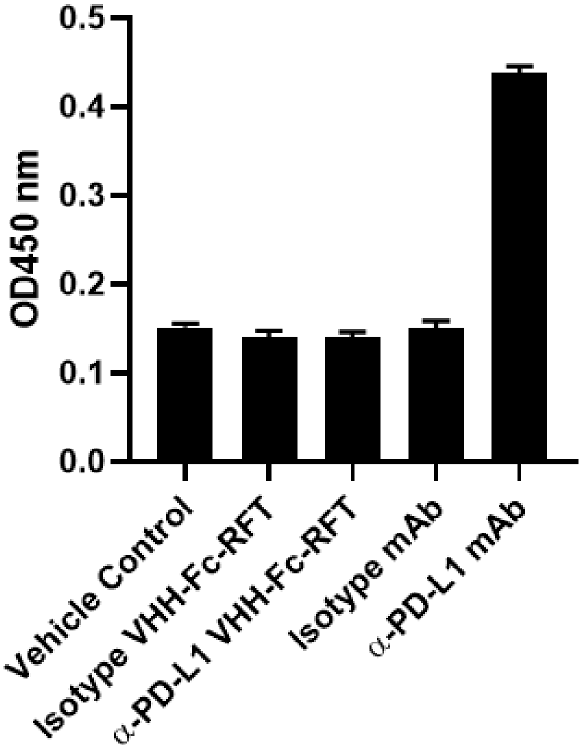

**Supporting Figure 20. α-PD-L1 VHH-Fc does not block the PD-1/PD-L1 interaction.** Raji PD-L1 cells were treated with staphylococcal enterotoxin D (SED) for 30 min and combined with Jurkat PD-1 cells pre-mixed with Isotype VHH-Fc-RFT, α-PD-L1 VHH-Fc-RFT, Isotype mAb, or α-PD-L1 mAb and incubated for 24 hours followed by analysis of IL-2 production. Bar plots of IL-2 production (as measured by an increase in $OD_{450}$) show that only the blocking α-PD-L1 mAb (clone MIH1) results in increased IL-2 production through disruption of the PD-1/PD-L1 interaction whereas the α-PD-L1 VHH does not have blocking effects. Error bars represent standard deviation of n = 3 experiments.

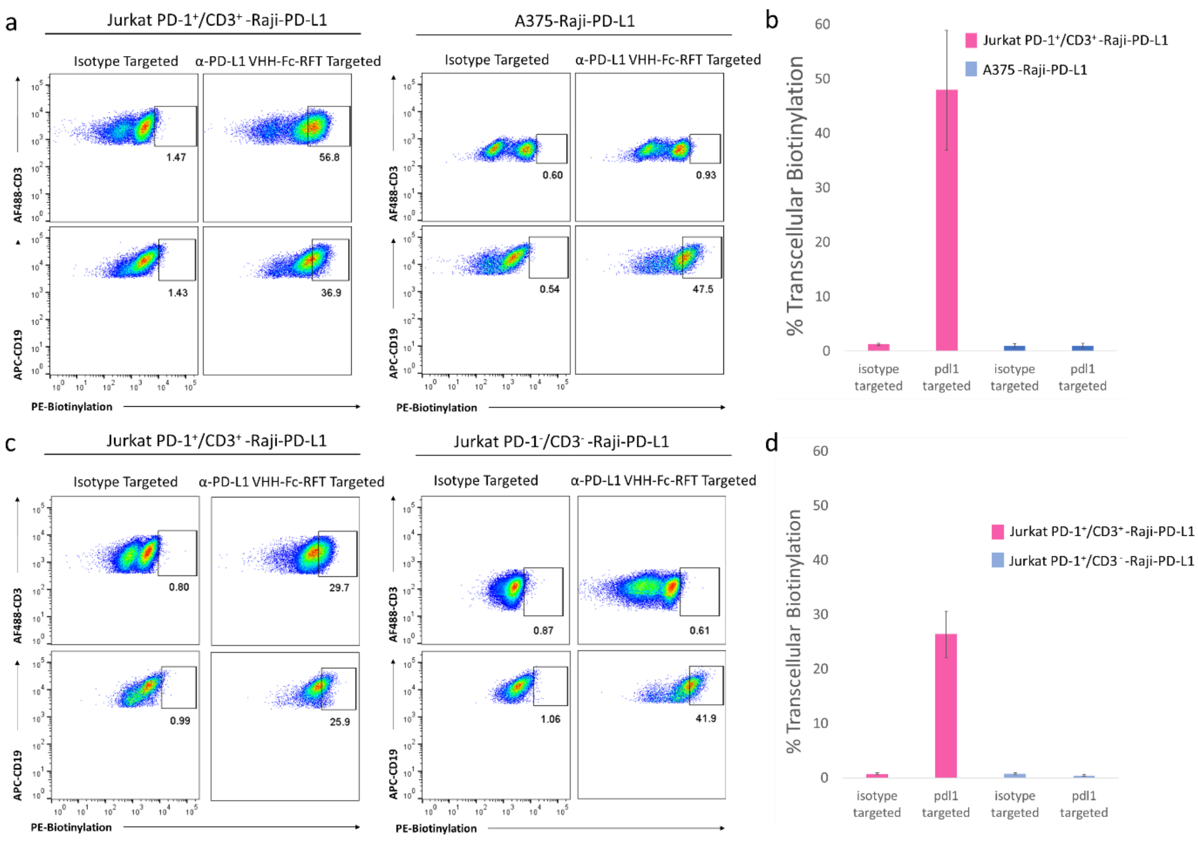

**Supporting Figure 21. Translabeling does not occur on A375 melanoma cells or Jurkat PD-1$^-$/CD3$^-$ cells.** a) Flow cytometry analysis of isotype (Isotype VHH-Fc-RFT) or α-PD-L1 VHH-Fc-RFT targeted labeling in a co-culture of A375 and Raji PD-L1 cells using 2 min light irradiation shows *cis*-labeling on Raji PD-L1 (CD19+) cells and no *trans*-cellular labeling on A375 cells. b) Bar plots of replicate analysis of cell biotinylation measured by flow cytometry in panel a for A375 cells. c) Flow cytometry analysis of isotype (Isotype VHH-Fc-RFT) or α-PD-L1 VHH-Fc-RFT targeted labeling in a co-culture of Jurkat PD-1$^-$/CD3$^-$ and Raji PD-L1 cells using 2 min light irradiation shows *cis*-labeling on Raji PD-L1 (CD19+) cells and no *trans*-cellular labeling on Jurkat PD-1$^-$/CD3$^-$ cells. d) Bar plots of replicate analysis of cell biotinylation measured by flow cytometry in panel a for Jurkat PD-1$^-$/CD3$^-$ cells. Error bars represent standard deviation of n = 3 experiments.

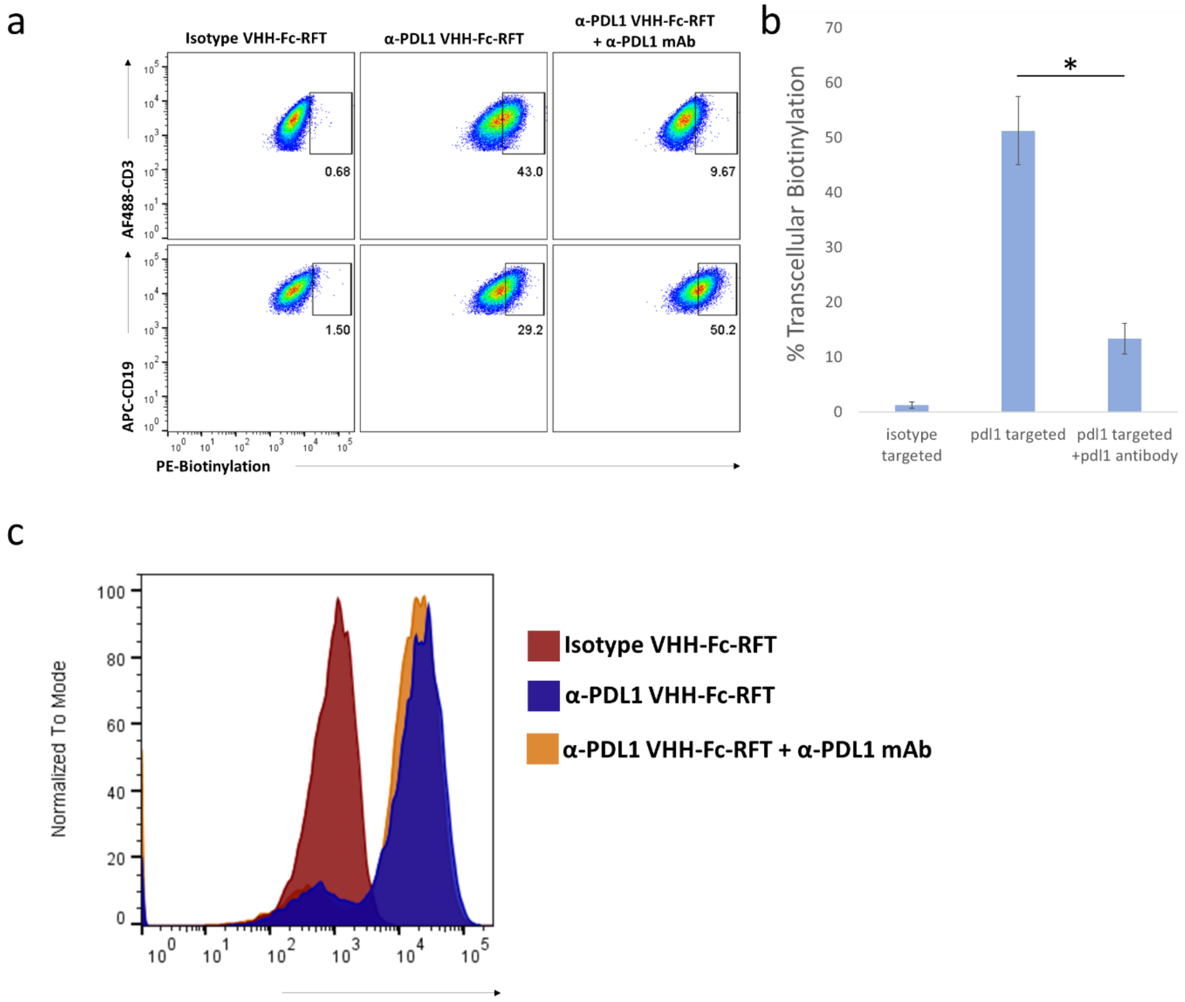

**Supporting Figure 22. Effect of anti-PD-L1 blocking antibody on transcellular labeling.** a) Flow cytometry analysis of isotype (Isotype VHH-Fc-RFT) or α-PD-L1 VHH-Fc-RFT targeted labeling in the Jurkat PD-1/Raji PD-L1 two cell system in the presence or absence of a PD-L1 blocking mAb (clone MIH1) using 2 min light irradiation shows *cis*-labeling on Raji PD-L1 (CD19+) cells and *trans*-cellular labeling on Jurkat-PD-1 (CD3+) cells. Addition of the PD-L1 mAb significantly reduces transcellular labeling on Jurkat cells. b) Bar plots of replicate analysis of cell biotinylation measured by flow cytometry in panel a for Jurkat PD-1 cells. Error bars represent standard deviation of n = 4 experiments; *$P < 0.01$. c) Flow cytometry analysis of cell surface biotinylation on Raji PD-L1 cells using α-PD-L1 VHH-Fc-RFT (blue) shows higher biotinylation compared to isotype targeted labeling (red) and that the presence of α-PD-L1 blocking antibody (α-PD-L1 mAb, clone MIH1) (orange) does not affect cell surface biotinylation by α-PD-L1 VHH-Fc-RFT.

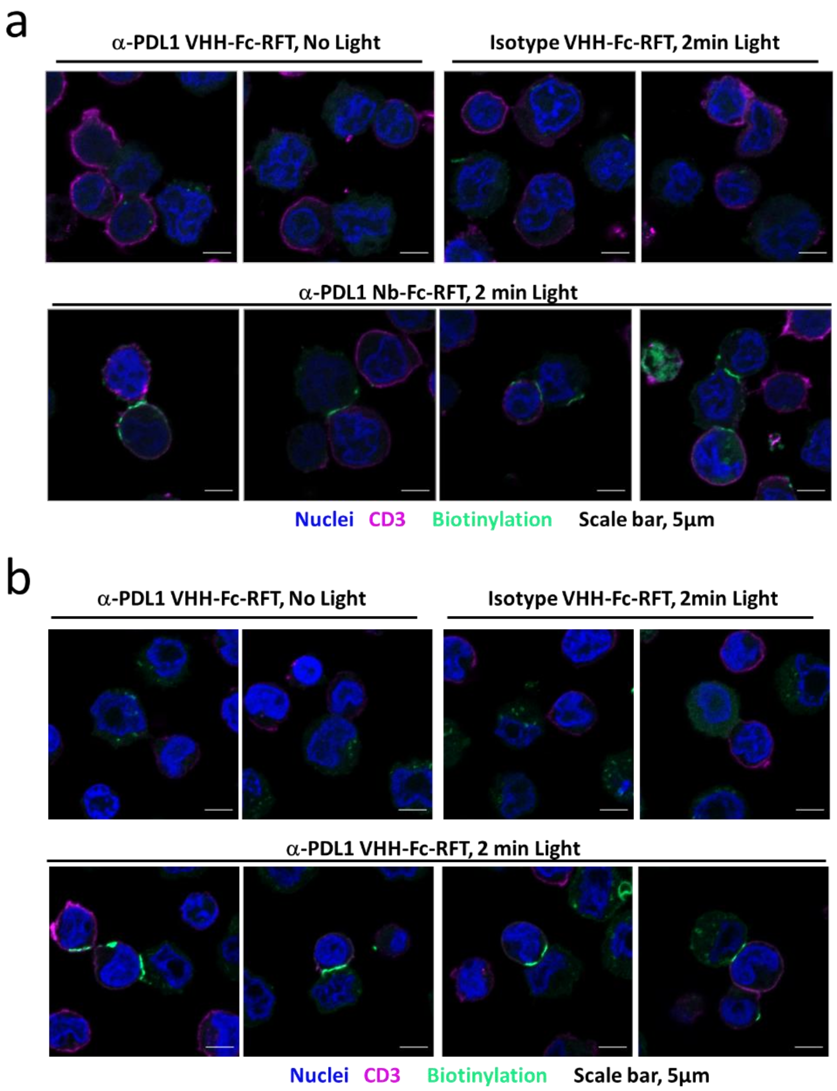

**Supporting Figure 23. α-PD-L1 VHH-Fc-RFT-mediated synaptic labeling of Jurkat PD-1/JY PD-L1 and Jurkat PD-1/CHO PD-L1 two cell systems.** Confocal microscopy imaging of a) Jurkat PD-1/JY PD-L1 or b) Jurkat PD-1/CHO PD-L1 two cell systems with α-PD-L1 VHH-Fc-RFT-targeted biotinylation on JY PD-L1 cells in the presence of 2 min visible light irradiation results in selective synaptic biotinylation whereas no biotinylation occurs in the absence of light or with isotype targeting. Cells were imaged for biotinylation (green), CD3 surface expression (magenta), and nuclei (Hoechst stain, blue). Scale bars indicate 5 μm.

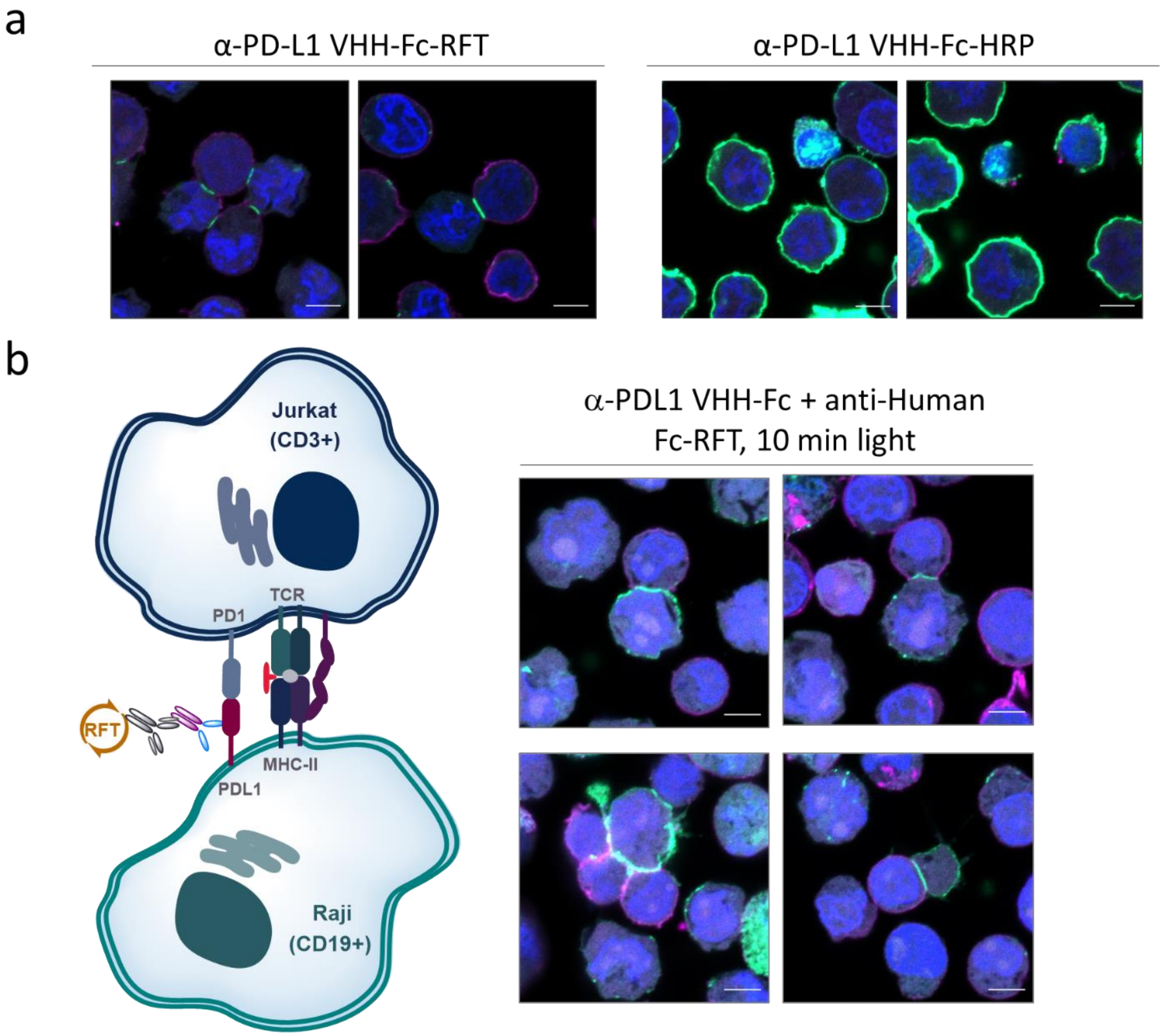

**Supporting Figure 24. Effect of peroxidase-based labeling and VHH/secondary-RFT based labeling on synaptic biotinylation.** Confocal microscopy imaging of a) α-PD-L1 VHH-Fc-RFT-targeted or b) α-PD-L1 VHH-Fc-Peroxidase-targeted biotinylation in the Jurkat PD-1/Raji PD-L1 two-cell system in the presence of biotin tyramide and 2 min visible light irradiation or 1 min in the presence of hydrogen peroxide and biotin tyramide, respectively. Complete labeling around both cell types is observed with the peroxidase-based system. b) Schematic of synaptic labeling with PD-L1 VHH-Fc and an anti-human Fc antibody RFT conjugate. c) Confocal microscopy imaging of PD-L1 VHH-Fc/anti-Human Fc-RFT-targeted biotinylation in the Jurkat PD-1/Raji PD-L1 two cell system in the presence of biotin tyramide and 10 min visible light irradiation. Unlike, the VHH system, complete extrasynaptic *cis*-labeling around the Raji PD-L1 cell is observed with the secondary antibody likely through the larger size of the labeling system that prevents facile access to the synapse. Cells were imaged for biotinylation (green stain), CD3 surface expression (magenta stain), and nuclei (Hoechst stain). Scale bars indicate 5 µm.

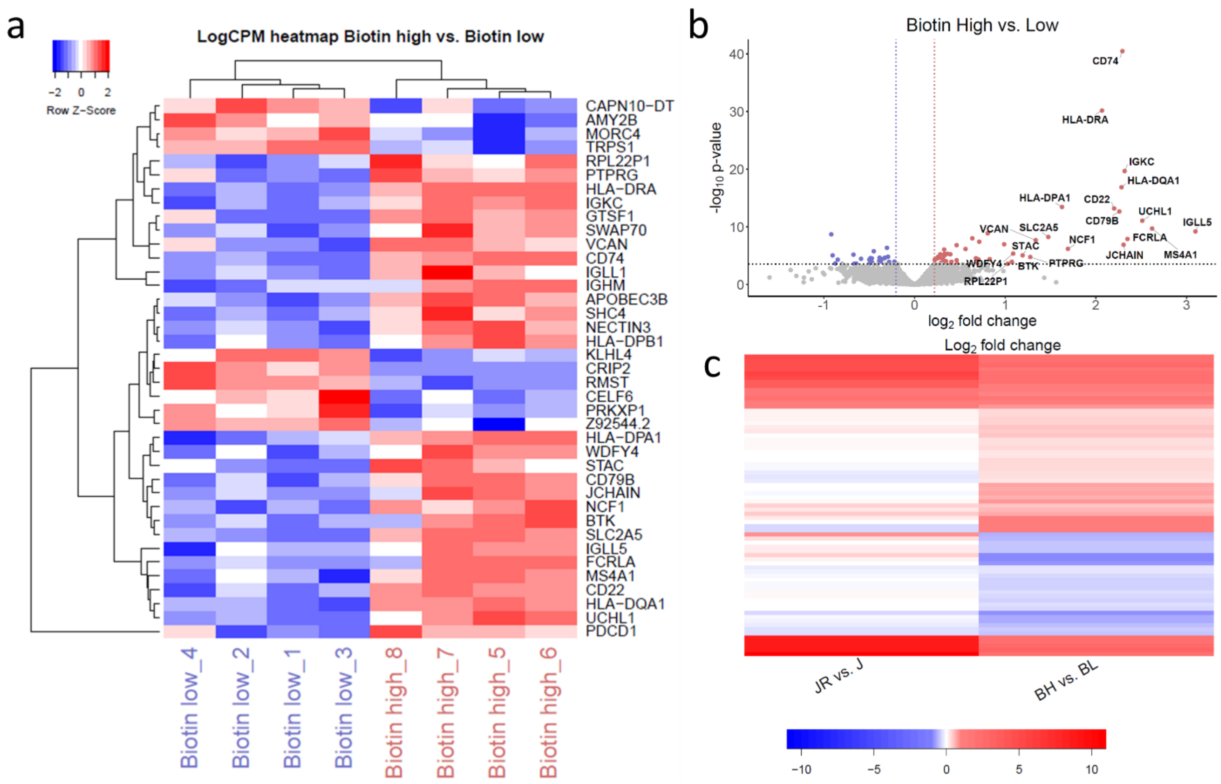

**Supporting Figure 25. Differential gene expression analysis of biotin high and biotin low cells.** a) Heat map of differential RNA expression levels (FDR<0.05) from biotin high vs. low Jurkat cell populations as described in figure 5). Gene data in the form of log2 count per million reads from 8 samples was clustered using hierarchical clustering with Euclidean distance and using the complete clustering algorithm. Clustering was performed on both genes and samples. Z-score was performed across rows with blue indicating negative and red positive z-scores. b) Volcano plot showing differential gene expression from biotinylated high vs. low Jurkat populations. The x-axis shows $\log_2$ fold change values and the y axis show the negative $\log_{10}$ p-value. Significantly enriched genes (FDR corrected p-value < 0.05, and $\log_2$FC>1) are labeled by gene name and colored according to GO terms which were enriched in upregulated genes. c) Differential gene expression of biotin high vs. biotin low (BH vs BL) cells and Jurkat+Raji vs Jurkat only cells (JR vs J) shows similar changes to the gene expression profile.

### General Materials

Ascorbic acid (BP321-500) and Ethanol (BP2818100) were purchased from Fisher Scientific (Pittsburgh, PA). Bovine Serum Albumin (BSA) (A7906-100G), Eppendorf Protein LoBind tubes (Z666505-100EA), A375 cells (88113005-1VL), HEK 293 cells (85120602-1VL) and Trolox (238813) were purchased from Sigma-Aldrich (St. Louis, MO). Sodium Azide (14314) was purchased from Alfa Aesar (Haverhill, MA). Biotin tyramide (LS-3500.1000) was purchased from Iris Biotech GMBH. RIPA Buffer (89900), Click-iT™ Protein Reaction Buffer Kit (Thermo Fisher Scientific: C10276), DPBS (14190144) and iBright Prestained Protein ladder (LC5615) were purchased from Thermo Scientific (Rockford, IL). TBST (IBB-581X) was purchased from Boston BioProducts (Ashland, MA). 5M Sodium Chloride (S24600-500.0) was purchased from Research Products International (Mt. Prospect, IL). Criterion XT 12% Tris TGX precast gels (5671044) and 4x Laemmli sample buffer (161-0747) were purchased from Bio-Rad (Hercules, CA). Goat α-rabbit secondary antibody (AP124) and Goat α-rabbit secondary antibody, Peroxidase conjugated (AP124P) was purchased from Millipore (Billerica, MA). 20% SDS solution (351-066-721) was purchased from Quality Biological (Gaithersburg, MD). JY wt and JY PD-L1 cells were a gift from Rene De Waal Malefyt and Sabine Le Saux (MRL, Merck & Co., Inc., Palo Alto, CA, USA). The Jurkat PD-1 and Raji PD-L1 co-culture system[9] was a gift from Aarron Willingham and Bhagyashree Bhagwat (MRL, Merck & Co., Inc., Palo Alto, CA, USA). PD-L1 aAPC/CHO-K1 cell system (J1252) was purchased from Promega (Madison, WI).

### General Method for Photolabeling of Proteins

Proteins were prepared in PBS to a final concentration of 1mg/ml protein followed by addition of biotin tyramide (25mM stock in DMSO) to give a final concentration of 250µM. To this mixture was added RFT to a final concentration ranging from 1-50µM depending on the experiment. The samples were irradiated with visible light in the bio-photoreactor (Efficiency Aggregators, BPR200) for the indicated time points. After visible light irradiation, 50µl of each sample was removed and mixed with 50µl of 4x loading buffer and then boiled at 95°C for 5 min. The samples were then analyzed by western blot according to the general Western blotting procedure described below. Note, for the α-chymotrypsinogen A samples, the labeling reaction was performed at pH 5.0 to prevent autolysis of the protein.

### General Cell Culture

Jurkat PD-1 cells were grown in RPMI 1640 1x with L-glutamine (Corning: 10-040-CV) containing 10% FBS (HyClone: SH30910.03), 1X MEM Non-Essential Amino Acid Solution (MEM NEAA, Sigma: M7145-100ML), 10mM HEPES Buffer (Fisher: BP299-100), 1mM Sodium Pyruvate (Corning: 25-000-CI), 2mM L-Glutamine (Lonza: 17-605E), 500µg/ml Geneticin (Thermo Fisher Scientific: 10131-035) and 20ng/ml Puromycin (Thermo Fisher Scientific: A11138-03).

Raji PD-L1 cells were grown in RPMI 1640 1x with L-glutamine (Corning: 10-040-CV) containing 10% FBS (HyClone: SH30910.03), 2mM L-Glutamine (Lonza: 17-605E), and 0.25µg/ml Puromycin (Thermo Fisher Scientific: A11138-03).

A375 cells were grown in 1x DMEM with GlutaMAX-I (Thermo Fisher Scientific: 10569-010) containing 10% FBS (Thermo Fisher Scientific: 10082-139), and 100 IU Penicillin/100µg/ml Streptomycin (Thermo Fisher Scientific: 15140-148).

JY wildtype (wt) and JY PD-L1 cells were grown in RPMI 1640 1x with L-glutamine (Corning: 10-040-CV) containing 10% FBS (HyClone: SH30910.03), 100 IU Penicillin/100µg/ml Streptomycin (Corning: 30-002-CI), 2mM L-Glutamine (Lonza: 17-605E), 1X MEM NEAA (Sigma: M7145-100ML), and 1mM Sodium Pyruvate (Cellgro: 25-000-CI).

Jurkat PD-1 Effector cells (Promega: J1252) were grown in RPMI 1640 (1x) with L-glutamine and HEPES (Corning: 10-041-CV) containing 10% FBS (Corning: 35-015-CV) for the first passage, after which the cells were kept in RPMI 1640 (1X) with L-glutamine and HEPES (Corning: 10-041-CV) containing 10% FBS (Corning: 35-015-CV), 200µg/ml Hygromycin B (Gibco: 10687010), 500µg/ml Antibiotic G-418 Sulfate Solution (Promega: V8091), 1 mM Sodium Pyruvate (Gibco: 11360), and 0.1 mM MEM NEAA (Gibco: 11140-050) per the manufacturer's instructions.

PD-L1 aAPC/CHO-K1 cells (Promega: J1252) were grown in Ham's F-12 medium (Gibco, Cat: 11765-054) containing 10% FBS (Corning: 35-015-CV), 200µg/ml Hygromycin B (Gibco: 10687010), and 250µg/ml Antibiotic G-418 Sulfate Solution (Promega, Cat: V8091). For passaging, cells were suspended from the flask using 0.25% Trypsin-EDTA (1x) (Gibco: 25200-056).

Jurkat TCR- (Jurkat ΔCD3/TCRαβ cells (JRT3-T3.5) (ATCC: TIB-153)) were grown in RPMI 1640 1x with L-glutamine (Corning: 10-040-CV) containing 10% FBS (HyClone: SH30910.03) and 100 IU Penicillin/100µg/ml Streptomycin (Corning: 30-002-CI).

HEK 293 cells were grown in 1x DMEM with GlutaMAX-I (Thermo Fisher Scientific: 10569-010) containing 10% FBS (Thermo Fisher Scientific: 10082-139), and 100 IU Penicillin/100µg/ml Streptomycin (Thermo Fisher Scientific: 15140-148).

All cells were grown at 37°C with 5% $CO_2$ in 25cm$^2$ (Corning: 430639), 75cm$^2$ (Corning: 430641U) or 150cm$^2$ (Corning: 430825) cell culture flasks (for Jurkat NF-κB, Jurkat PD-1, Raji PD-L1, JY wt and JY PD-L1 cells), or 10-cm (Falcon: 353003), 150-cm (Falcon: 168381) or 6-well plates (Falcon: 353046) for A375 cells as needed. For passaging, cells were suspended from the plate using 0.05% Trypsin-EDTA (1x) (Gibco: 25300054) as indicated.

All cell culture media was filter-sterilized using 0.2 µm Nalgene™ Rapid-Flow™ Sterile Disposable Filter Units with PES Membrane, 1,000ml capacity (Thermo Scientific: 567-0020) or 500ml capacity (Thermo Scientific: 569-0020).

### General Western Blotting Procedure

Samples were loaded onto 12% TGX Criterion gels (Bio-Rad: 5671044) and run at 180V to resolve the proteins. The proteins were then blotted onto PVDF or nitrocellulose membranes using the iBlot 2 gel transfer device (Thermo Fisher Scientific: IB21001). Membranes were blocked in TBST with 3% BSA followed by overnight incubation with IRDye 800CW Streptavidin at a 1:5,000 dilution (LI-COR: 926-32230), rabbit α-human CD45 monoclonal antibody (clone D9M8I, Cell Signaling Technology: 13917S) at a 1:5,000 dilution, or rabbit α-human PD-L1 monoclonal antibody (clone E1L3N, Cell Signaling Technology: 13684S) at a 1:5,000 dilution, all in TBST with 3% BSA. Membranes were washed 3xs with TBST and 1x with Milli-Q water and then imaged on the LI-COR Odyssey CLx to evaluate biotinylation. For α-CD45 or α-PD-L1 Western blots, membranes were further incubated with IRDye 680RD Goat anti-Rabbit 680 (LI-COR, Cat# 926-68071) for 1 hour in TBST, washed 3x with TBST and imaged on the LI-COR. Following imaging, membranes were stained with REVERT total protein stain (LI-COR: 926-11016) according to manufacturer's instructions and re-imaged on the LI-COR Odyssey CLx.

### Bio-photoreactor

The bio-photoreactor BPR200 designed by Merck & Co., Inc., Kenilworth, NJ, USA and Efficiency Aggregators (Richland, Tx, Fisher catalog number: NC1558343 BPR200) and manufactured by Efficiency Aggregators was used for all photolabeling experiments, using the 24-well microcentrifuge tube holder attachment. The low heat generation of this device prevents exposure of live cells to excessive heating that is observed with traditional blue LED systems. To operate the system, the bio-photoreactor chamber door was opened with the device light off, and the samples were transferred directly from ice onto the tube rack (see image below). The door was closed, and the light was turned on to 100% intensity using the device control app for the indicated time points. For each 0 min time point, the sample was left on ice and kept away from light for the duration of the photolabeling experiment.

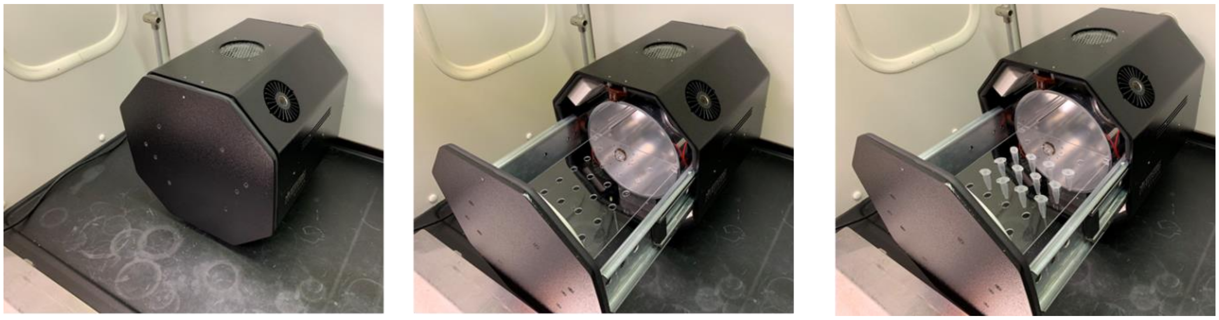

**Supporting Figure 26.** Left panel, bio-photoreactor (switched off), door closed. Middle panel, bio-photoreactor (switched off), door open. Right panel, bio-photoreactor (switched of), door open with Eppendorf tubes inserted.

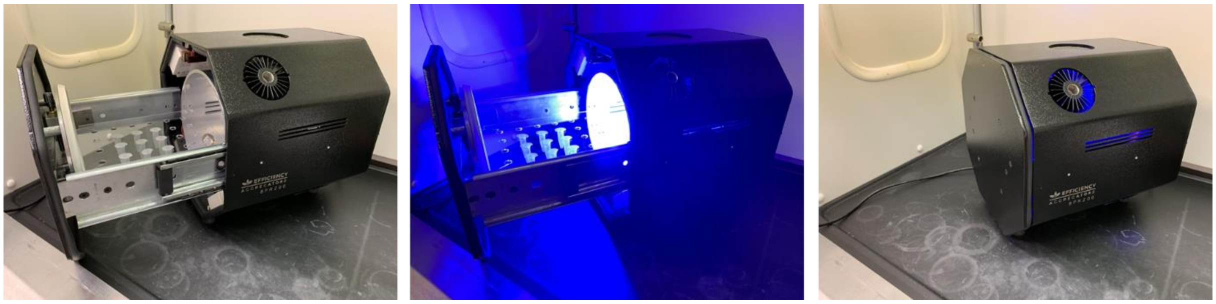

**Supporting Figure 27.** Left panel, bio-photoreactor (switched off) containing microcentrifuge tubes in sample rack. Middle panel, bio-photoreactor (switched on) with visible light irradiation at full intensity. Right panel, bio-photoreactor (switched on) in normal operating mode at full light intensity with chamber door closed.

### Photocatalyst Screen for BSA Biotinylation

BSA was prepared in PBS to a final concentration of 1mg/ml protein, followed by addition of biotin tyramide (using a 25mM stock in DMSO) to a final concentration of 250µM. Different flavin photocatalysts were added to this mixture, as outlined in Supporting Figure 2. For the indicated samples, ammonium persulfate (Sigma-Aldrich: A7460-100G) was added to a final concentration of 500µM. The samples were irradiated with visible light in the bio-photoreactor (Efficiency Aggregators, BPR200) for the indicated time points. After light irradiation, 50µl of each sample was removed and mixed with 50µl of 4x loading buffer and then heated to 95°C for 5 min. The samples were then analyzed by western blot according to the general western blotting procedure described above.

#### Flavin Photostability Analysis by UV-Vis Spectroscopy

200µM flavin (Rb, RFT, Lumiflavin) alone, or mixed with 5mM biotin tyramide, or 5mM biotin tyramide and 1mg/ml BSA was added to a 5ml Eppendorf tube (Eppendorf: 0030119401). The tube was placed into the bio-photoreactor and irradiated with visible light for the indicated time points (see Supporting Figure 3). At each time point, 300µl was removed and transferred to a well of a 96-well flat bottom plate and stored at room temperature in the dark until all time points were taken. A full absorbance spectrum for each sample was measured on a PHERAStar FSX plate reader.

#### Light ON/OFF Protein Labeling Experiment

BSA was prepared in 1x PBS to a final concentration of 1mg/ml protein followed by addition of biotin tyramide (using a 25mM stock in DMSO) to a final concentration of 250µM. RFT was then added to a final concentration of 1µM. After mixing, 50µl of sample was removed as the 0-min time point. The remaining samples were placed into the bio-photoreactor and irradiated with visible light in 1-min intervals. The light was turned off and the samples were left in the bio-photoreactor for 5 min between each 1-min light interval. At the desired time points, 50µl was removed and combined with 50µl of 4x loading buffer and then heated to 95°C for 5 min. The samples were then analyzed by western blot according to the general western blotting procedure described above.

#### Trapping of Tyrosyl Radicals with Biotin-DMAA Probe on BSA

BSA was prepared in PBS to a final concentration of 1mg/ml protein, followed by addition of biotin tyramide or biotin dimethyl amino aniline (biotin-DMAA) to a final concentration of 250µM. To this mixture RFT or Lumiflavin was added to a final concentration of 10µM. The samples were irradiated with visible light in the bio-photoreactor for the indicated time points. After visible light irradiation, 50µl of each sample was removed and mixed with 50µl of 4x Laemmli sample buffer and then heated at 95°C for 5 min. The samples were then analyzed by western blot according to the general western blotting procedure described above.

#### Trapping of Oxidative species with Biotin-$ONH_2$ Probe on BSA

BSA was prepared in PBS to a final concentration of 1mg/ml protein, followed by addition of biotin tyramide or Biotin-PEG3-oxyamine HCl salt (Broadpharm, BP-22179) to a final concentration of 250µM. To this mixture RFT was added to a final concentration of 1µM. The samples were irradiated with visible light in the bio-photoreactor for 5 minutes. After visible light irradiation, 50µl of each sample was removed and mixed with 50µl of 4x Laemmli sample buffer and then heated at 95°C for 5 min. The samples were then analyzed by Western blot according to the general western blotting procedure described above.

#### Measurement of Hydrogen Peroxide Generation upon Flavin Photoactivation

RFT (1µM final concentration) or RFT (1µM final concentration) mixed with BSA (1mg/ml) and biotin tyramide (250µM) was irradiated with visible light for the indicated time points. At each time point, 5µl was removed and Hydrogen Peroxide amounts were determined by fluorometric analysis (Sigma-Aldrich: MAK165) according to manufacturer's instructions.

#### Photolabeling of Proteins with Bioorthogonal Functional Groups

BSA was prepared in PBS to a final concentration of 1mg/ml protein, followed by addition of phenol alkyne, phenol azide, or phenol DBCO, to give a final concentration of 250µM. RFT was added to this mixture, to a final concentration 10µM. The samples were irradiated with visible light in the bio-photoreactor for the indicated time points. After irradiation, 300µl of sample was transferred to an

Eppendorf tube followed by addition of 1mM biotin azide (for conjugation to phenol alkyne- or phenol DBCO-labeled protein) or 1mM biotin alkyne (for conjugation to phenol azide-labeled protein). For phenol azide or alkyne labeled protein, click chemistry was performed using the Click-iT™ Protein Reaction Buffer Kit (Thermo Fisher Scientific: C10276), according to manufacturer's instructions. For click reactions involving the phenol DBCO-labeled proteins, the samples were incubated overnight at room temperature with no further additives. After the click reaction, 50µl of each sample was removed and mixed with 50µl of 4x loading buffer and then heated to 95°C for 5 min. The samples were then analyzed by Western blot according to the general Western blotting procedure described above. α-chymotrypsinogen A was labeled with 680R Tyramide fluorophore (Biotium: 92196) using the same conditions as above. After irradiation, 50µl of each sample was removed and mixed with 50µl of 4x loading buffer and then heated to 95°C for 5 min. The samples were then analyzed by gel fluorescence scanning at 700nm on the LI-COR Odyssey CLx.

#### Biotinylation of BSA with Horse Radish Peroxidase

BSA was prepared in 500 µl PBS to a final concentration of 1mg/ml protein, followed by addition of biotin tyramide 500µM, and 1 µM $H_2O_2$. 2.5 µg HRP-IgG (Millipore AP124) was then added to start the reaction. At the desired time points, 40µl of sample was removed and added to an Eppendorf containing 40µl of quench buffer (PBS containing 10mM Trolox, 10mM $NaN_3$, and 10mM Ascorbic Acid). 20µl of 4x loading buffer was added to each tube and the samples were boiled for 5min at 95°C. 15µl of each sample was then analyzed by western blot according to the general Western blotting procedure described above.

#### Flavin-based Biotinylation of 293 Cell Surface via SNAP-tag Fusion Protein

HEK 293 cells were seeded into a 6 well plate at 500,000 cell/well and grown in complete media (DMEM High glucose, +pyruvate, +10% FBS, +penicillin/streptomycin) for 24 hours. The cells were transfected with pSNAP-APELIN Receptor plasmid (Fisher: 50-246-959) using the Mirus TransIT-293 transfection reagent (Mirus, MIR2700) according to manufacturer's instructions. After transfection, the cells samples were incubated for 36 hours at 37°C. After 36 hours, the media was removed, and the cells were gently washed 2xs with complete media. To the cells was then added 2 mL of complete media containing 25µM $O^6$-benzylguanine RFT photocatalyst (BG-RFT) and incubated for 15 min 37°C. The cells were then gently washed 3xs with 2 mL complete media and 1x with 2 mL cold PBS. After washing, 2 mL cold PBS containing 250 µM biotin tyramide was then added to each well followed by light irradiation for 5 min in the Bio-photoreactor or 5 min in the dark as a no light control. The cells were then washed 2xs with 1mL cold DPBS and then resuspended in 0.5 mL 1x DPBS.

For microscopy analysis, round-shaped glass microscope coverslips (Fisherbrand: 12-545-81) were acid-etched by incubating in 1N HCl (Fisher: SA56-1) for 30 min at 50°C. The coverslips were then washed in distilled water 3x and placed in 100% ethanol (Fisher: BP2818-500). Acid-etched glass coverslips were placed into a 24-well plate (Thermo Fisher Scientific: 142485), one coverslip per well. Coverslips were washed with 1ml of 1x DPBS (Gibco: 14190-144) 2x to remove ethanol. 1ml of poly-L-lysine solution (Sigma: P4707-50ML) was added per well and the plate was incubated for 30 min at 37°C. Coverslips were washed 2x with 1ml of 1x DPBS. Labeled cells were loaded on the 24-well plate and centrifuged at 400xg for 3 min with deceleration set at 3.

After centrifugation, 6% PFA (PFA, Ted Pella: 18505-100) and 0.2% glutaraldehyde (Sigma-Aldrich: G5882-10X10ML) in 1x DPBS were added 1:1 drop wise for a final concentration of 3% PFA and 0.1% glutaraldehyde per well. Samples were incubated in fixative for 10 min at 4°C. The fixative was removed, and the coverslips were washed 3x in Stain Buffer (BD Biosciences: 554656) and incubated overnight in 1 ml of Stain Buffer at 4°C. The following day, the samples were stained with Alexa Fluor 488 streptavidin (BioLegend: 405235) at a 1:200 dilution in 500µl of Stain Buffer. The plate was sealed with parafilm, covered in foil, and incubated overnight at 4°C. The next day, the coverslips were washed 1x in 500µl of Stain Buffer. Hoechst DNA dye (Cayman Chemical Company: 600332) was added at a 1:10,000 dilution in 500µl of Stain Buffer and incubated for 10 min at room temperature, protected from light. The coverslips were washed 2x in Stain Buffer, followed by a final fixing step with 500µl of 3% PFA and 0.1% Glutaraldehyde solution in 1x DPBS for 5 min at room temperature. The coverslips were washed 2x in Stain Buffer. One drop of ProLong Gold Anti-fade mountant (Invitrogen: P36934) was added to each microscope slide (J. Melvin Freed Brand: 301MF, frosted, 3 x 1”) using High Precision Straight Tapered Ultra Fine Point tweezers (Fisherbrand: 12-000-122). Each coverslip was placed on top of the mountant on their respective slides and allowed to dry overnight at room temperature, protected from light. The slides were imaged using a Zeiss LSM800 inverted, confocal microscope using a 63X oil immersion objective and Airyscan settings.

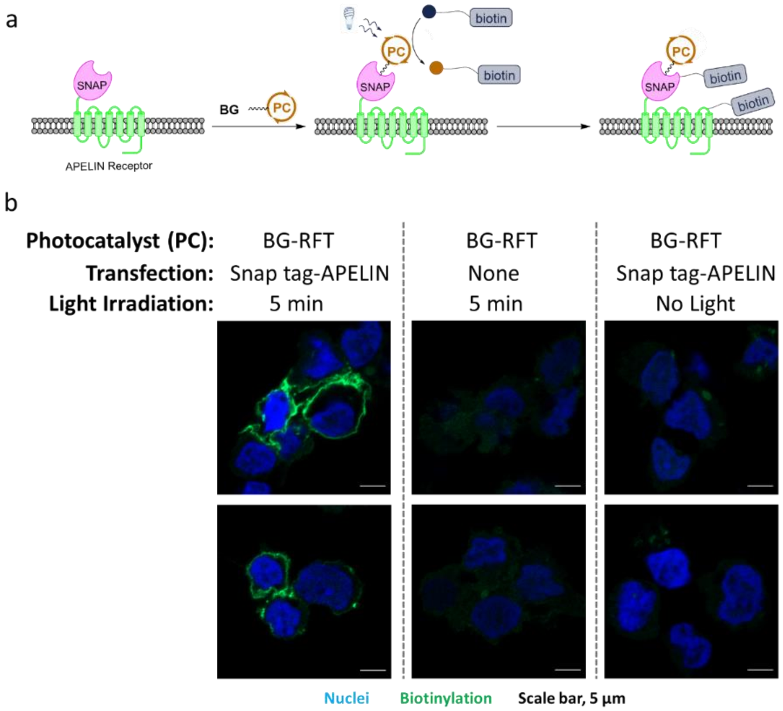

**Supporting Figure 28. Flavin-mediated cell surface tagging using SNAP-tag self-labeling enzyme**. a) Schematic depicting cell surface labeling using cell surface SNAP-tag fusion protein. Briefly, the SNAP tag-APELIN receptor fusion protein was expressed on HEK 293 cells and labeled with $O^6$-benzylguanine photocatalyst (PC) (BG-PC). After labeling, the cells were then biotinylated via irradiation with visible light in the presence of biotin tyramide. b) Cell surface labeling showing three difference conditions. Left condition, HEK 293 were transfected with SNAP-tag APELIN, labeled with BG-RFT photocatalyst and irradiated with visible light for 5 min. Middle condition, non-transfected HEK 293 cells were labeled with BG-RFT photocatalyst and irradiated with visible light for 5 min. Right condition, HEK 293 were transfected with SNAP-tag APELIN, labeled with BG-RFT photocatalyst incubated in the dark for 5 min. Biotinylation was only detected in the presence of SNAP-tag APELIN expression, BG-RFT, and visible light irradiation. Cells were imaged for biotinylation (green), and nuclei (Hoechst stain, blue). Scale bars indicate 5 µm.

### LC-MS/MS Analysis of Purified Proteins Biotinylated in vitro

BSA (Sigma-Aldrich: A7906-100G), Rabbit Phosphorylase (Sigma-Aldrich: P6635-25MG), Chicken Ovalbumin (Sigma-Aldrich: A2512-1G), Carbonic Anhydrase II (Sigma-Aldrich: C2624-100MG), or α-chymotrypsinogen A (Sigma-Aldrich: C4879-1G) were biotinylated following the general protein photolabeling method above and submitted for proteomic sample preparation and LC-MS/MS analysis at MSBioworks (Ann Arbor, MI). Briefly, 10µg of total protein was reduced with 10mM dithiothreitol at 60°C for 30 min, followed by alkylation with 15mM iodoacetamide at room temperature for 45 min. Samples were digested with 0.5µg of either trypsin (Promega: V5111), chymotrypsin (Promega: V1061) or elastase (Promega: V1891) overnight at 37°C. The samples were then acidified with formic acid and desalted using 3M Empore SD solid phase extraction plates. The samples were then lyophilized and reconstituted in 0.1% TFA for analysis by nano-UPLC-MS/MS using a Waters NanoAcquity UPLC system interfaced with a Thermo Fisher Scientific Q Exactive mass spectrometer. Peptide material (1µg) was loaded on a trapping column and eluted over a 75µm analytical column at 350nl/min; both columns were packed with Phenomenex Luna C18 resin. The mass spectrometer was operated in data-dependent mode, with the Orbitrap operating at resolution settings of 70,000 FWHM and 17,500 FWHM for MS and MS/MS, respectively. The fifteen most abundant ions were selected for MS/MS. Peptide assignments from the raw data were achieved with the Byonic search engine (Protein Metrics) using a custom database consisting of BSA (P02769), Carbonic Anhydrase II (P00921), Chymotrypsinogen A (P00766), Rabbit Phosphorylase (P00489) and Chicken Ovalbumin (P01012) protein sequences using the following parameters: enzymatic cleavage was set to either trypsin with up to 3 missed cleavages (for tryptic samples) or non-specific (for chymotrypsin and elastase samples), variable modifications included Carbamidomethyl (C), Deamidated (N), Oxidation (C,H,M,P,W,Y), and Biotin tyramide, $C_{18}H_{23}N_3O_3S$ (Y,C,H,F,W,S,T,D,E), and peptide and fragment mass tolerance was set to 10 ppm and 0.02 Da, respectively. Sequence coverage maps and annotated peptide spectral matches were generated with Scaffold PTM V3.2 (Proteome Software). Data were filtered for a minimum localization probability of 50% and putative modified sites were validated manually. Modifications were considered correctly assigned if ladders of fragment ions spanning both the modified residue and its flanking *N*- and *C*-terminal amino acids were identified.

### Labeling of Antibody with Riboflavin Tetraacetate (Antibody Flavin Conjugate Preparation)

300µl of polyclonal Goat α-Mouse IgG (AP124) was added to a Protein LoBind tube, followed by addition of 30µl of 1M Sodium Bicarbonate buffer (pH 8.5) added to the reaction. A stock solution of azide-NHS linker (C3 linker) was prepared in DMSO (100mM final concentration). 3µl of this stock solution was added to the antibody reaction mixture and incubated for 1.5 hours in the dark. After 1.5 hours, an additional 3µl of the stock solution was added, and the sample was incubated for another 1.5 hours. Following incubation, the sample was then placed onto a Zeba Spin desalting column (Thermo Fisher Scientific: 87769, 2ml column, 40,000 MWCO) and buffer exchanged into 50mM Tris pH 8.0, according to manufacturer's instructions. 200µl of the labeled antibody was then added to a Protein LoBind tube for a Click chemistry reaction using the Click-iT™ Protein Reaction Buffer Kit (Thermo Fisher Scientific: C10276) according to manufacturer's instructions. Briefly, 300µM RFT (final concentration from a 5mM stock in DMSO) was added to the azide labeled antibody. 12.5µl of copper sulfate was added to the reaction followed by 12.5µl of additive 1. The sample was briefly mixed and incubated at room temperature for 3 min followed by addition of 15µl of additive 2. The sample was then incubated in the dark for 30 min at room temperature. The sample was then placed onto a Zeba Spin desalting column (87769, 2ml column, 40,000 MWCO) and buffer exchanged into 1x PBS according to manufacturer's instructions. The final protein concentration of the antibody was measured using the BCA Protein Assay kit according to manufacturer's instructions (Thermo Fisher Scientific: 23227). RFT concentration was determined by measuring absorbance at 450nm (A450) and comparing to a standard curve of free RFT of known concentrations. The micromolar concentration of RFT was divided by the micromolar concentration of antibody to determine the antibody:RFT ratio. Under the above conditions, we routinely obtained an antibody:RFT ratio of 1:5.

#### Targeted Protein Labeling on Agarose Beads

50µl of anti-human IgG Fc specific agarose beads (Sigma-Aldrich: A3316-2ml) were transferred to a Protein LoBind tube and washed 2x with TBST (1 min at 1,000xg, 4°C). 30µg of PD-L1-human IgG (R&D Systems: 156-B7) and 30µg of CD45-human IgG (Sinobiologic: 10086-H02H) were added separately to 50µl beads in 1ml TBST and incubated for 30 min at 4°C. After incubation, the cells were pelleted by centrifugation (1 min at 1,000xg, 4°C) to remove the supernatant. The beads were then washed 2x with 1ml cold PBS, then were pelleted by centrifugation (4 min at 1,000xg, 4°C). The beads were then incubated with either 0.5µg α-PD-L1 antibody (Thermo Fisher Scientific: 14598382) or α-CD45 antibody (BD Biosciences, clone HI30: 555480) in 1ml TBST for 30 min at 4°C, followed by two washes with TBST as described above. After the final wash, the beads were pelleted by centrifugation (4 min at 1,000xg, 4°C) and resuspended in 1ml of TBST containing 0.5µg of α-Mouse IgG RFT conjugate and incubated for 30 min. The beads were then washed 2x with TBST as described above, followed by centrifugation for 1 min at 1,000xg, 4°C. beads were then resuspended in 400µl of 250µM biotin tyramide and incubated in the bio-photoreactor. At each of the indicated time points, 100µl of sample was removed from the light source and stored in the dark on ice to stop the reaction. The beads were pelleted by centrifugation (1 min at 1,000xg, 4°C). The supernatant was removed, and the beads were incubated with 25µl of 4x Laemmli sample buffer for 5 min at 95°C. 15µl of each sample was then analyzed by Western blot according to the general method described above.

#### Photolabeling of Membrane Proteins on Live Cells (Flow Cytometry Analysis)

5 million Jurkat PD-1, JY PD-L1, Jurkat wt, or JY wt per sample were centrifuged for 4 min at 800xg, 4°C and resuspended in 1 mL of cold 1x DPBS (Gibco) in Protein LoBind microcentrifuge tubes. 5 µg of each of the following primary antibodies were used to target surface proteins: Isotype control (BD Pharmingen, Purified Mouse IgG1 κ, clone MOPC-21: 556648), Mouse α-Human CD274 (PD-L1), clone MIH1 (Invitrogen: 14-5983-82), Mouse α-Human CD279 (PD-1), clone J116 (Invitrogen: 14-9989-82), Mouse α-Human CD45, clone HI30 (BD: 555480), Mouse α-Human CD86, clone 2331 (FUN-1) (BD: 555655). Samples were incubated for 30 min in a rotisserie at 4°C, followed by 2 washes in 1 mL of cold 1x DPBS with pelleting cells for 4 min at 800xg, 4°C in between washes. 5 µg Goat α-Mouse IgG RFT conjugate were added to all samples and incubated for 30 min in a rotisserie at 4°C, followed by 2 washes in 1 mL of cold 1x DPBS with pelleting cells for 4 min at 800xg, 4°C in between washes. Cell pellets were resuspended in 500 µL of 1x DPBS and biotin tyramide was added to all samples at a final concentration of 250 µM. Samples were incubated in the biophotoreactor (BPR200) for 0 or 2 min under visible light at 100% intensity. Cells were then centrifuged for 4 min at 800xg, 4°C, washed 1x in 1 mL 1x DPBS, and centrifuged again. Each pellet was resuspended in 100 µL of Stain Buffer (BD Biosciences: 554656) per well on V-bottom plates (Falcon: 3894) containing a 1:100 dilution of Fc block (BD Biosciences: 564220). The cells were incubated for 20 min on ice then centrifuged for 4 min at 500xg, 4°C, washed once with 200 µL Stain Buffer and centrifuged again as above. Cells were resuspended in 100 µL of Stain Buffer containing Streptavidin PE at a 1:200 dilution (BD Biosciences: 554061). Compensation controls were performed using BD FACSDiva software, UltraComp beads (BD Biosciences: 01-2222-42), BV421 Mouse α-Human CD45RO, clone UCHL1 (BD Biosciences: 562641), and PE Mouse α-Human CD3, clone SK7 (BioLegend: 344806). The plate was

incubated for 30 min on ice, the cells were washed once with 200 µL of cold 1x DPBS then resuspended in a 1:500 dilution of Zombie Violet Viability kit (BioLegend: 423113) and incubated for 15 min at room temperature, protected from light. The cells were centrifuged, washed 1x in 200 µL of cold 1x DPBS and centrifuged again. Pellets were resuspended in 200 µL of Stain Buffer and transferred to 5 mL FACS tubes (Fisherbrand: 14-956-3D) to acquire on a BD FACSCelesta (Model No: 660344) with FACSDiva software (v8.0.1.1). Data was analyzed using FlowJo v10 (FlowJo, LLC).

To evaluate α-Human PD-L1 antibody-mediated blockade of VHH-Fc-RFT PhoTag, 5 million Raji PD-L1 cells per sample were incubated with 2.5 µg of Isotype or α-PD-L1 VHH-Fc-RFT in 250 µL of cold Raji PD-L1 media for 1 hour in a rotisserie at 4°C in Protein LoBind microcentrifuge tubes (Eppendorf). Cells were washed 2x with 1 mL of cold Raji PD-L1 media with centrifugation for 4 min at 800xg and 4°C to pellet cells in between washes. Cells were resuspended in 250 µL of cold Raji PD-L1 media containing Mouse α-Human PD-L1 antibody, clone MIH1 (Invitrogen: 14-5983-82) at a final concentration of 10 µg/mL for 30 min on a rotisserie at 4°C. Cells were washed 2x in cold 1x DPBS and pelleted in between washes. Cell pellets were resuspended in 500 µL of 1x DPBS and labeling and staining to evaluate biotinylation levels by flow cytometry was performed as indicated above.

### Photolabeling of CD45 on Live Cells (Western blot Analysis)

Jurkat cells were washed 2x in cold 1x DPBS, resuspended in cold 1x DPBS at 5 million cells/ml and transferred to Protein LoBind tubes in 1ml aliquots. The cells were pelleted by centrifugation (4 min at 1,000xg, 4$^{o}$C) and then resuspended in 1ml of cold 1x DPBS containing 5µg Isotype control (BD Biosciences, Purified Mouse IgG1 κ, clone MOPC-21: 556648) or α-CD45 antibody (BD Biosciences, clone HI30: 555480). The cells were incubated on a rotisserie for 30 min at 4$^{o}$C. After incubation, the cells were pelleted by centrifugation (4 min at 1,000xg, 4$^{o}$C) to remove the supernatant. The cells were then washed 2x with 1ml cold 1x DPBS. After the final wash, the cells were pelleted by centrifugation (4 min at 1,000xg, 4$^{o}$C) and resuspended in 1ml of cold 1x DPBS containing 5µg of Goat α-Mouse IgG-RFT conjugate and incubated on a rotisserie for 30 min at 4$^{o}$C. The cells were then pelleted by centrifugation (4 min at 1,000xg, 4$^{o}$C) to remove the supernatant and washed 2x with 1ml cold 1x DPBS. After the final wash, the cells were pelleted by centrifugation (4 min at 1,000xg, 4$^{o}$C) and resuspended in 1ml of cold PBS containing 250µM biotin tyramide. The samples were left on ice (0-min treatment) or placed in the bio-photoreactor for 2 min or 10 min and irradiated at full light intensity. After irradiation, the cells were then pelleted by centrifugation (4 min at 1,000xg, 4$^{o}$C) to remove the supernatant and lysed in 150µl 4x Laemmli sample buffer (Bio-Rad: 161-0747) that was pre-diluted 1:1 in RIPA buffer (2x final concentration). The samples were sonicated (5s at power level 6) and then heated for 5 min at 95$^{o}$C. 15µl of each sample was then analyzed following the General Western Blot Procedure described above.

### Cellular Labeling with Biotin Phenol Analogs

Jurkat cells were washed 2x in cold 1x DPBS, resuspended in cold 1x DPBS at 5 million cells/ml and transferred to microcentrifuge tubes in 1ml aliquots. The cells were pelleted by centrifugation (4 min at 1,000xg, 4°C) and then resuspended in 1ml of cold 1x DPBS containing no antibody, 5µg isotype control (BD Biosciences Purified Mouse IgG1 κ, clone MOPC-21: 556648) or α-CD45 antibody (BD Biosciences, clone HI30: 555480) for Jurkat cells. The cells were incubated on a rotisserie for 30 min at 4°C. After incubation, the cells were pelleted by centrifugation (4 min at 1,000xg, 4°C) to remove the supernatant. The cells were then washed 2x with 1ml cold PBS. After the final wash, the cells were pelleted by centrifugation (4 min at 1,000xg, 4°C) and resuspended in 1ml of cold PBS containing 5µg of Goat α-Mouse IgG-RFT conjugate and incubated on a rotisserie for 30 min at 4°C. After incubation, the cells were pelleted by centrifugation (4 min at 1,000xg, 4°C) to remove the supernatant. The cells were then washed 2x with

1ml cold PBS. After the final wash, the cells were pelleted by centrifugation (4 min at 1,000xg, 4°C) and resuspended in 1ml of cold 1x DPBS containing 250µM of biotin tyramide or structurally-related analogs; biotin tyramide analogs are described in Supporting Figure 15 (all small molecules were prepared in 5mM DMSO stock). The samples were left on ice (0-min treatment) or placed in the bio-photoreactor for 5 min and irradiated at full light intensity. After irradiation, the cells were then pelleted by centrifugation (4 min at 1,000xg, 4°C) to remove the supernatant. The cell pellets were then lysed in 150µl loading buffer (Bio-Rad: 161-0747) that was pre-diluted 1:1 in RIPA buffer (2x final concentration). The samples were then sonicated (5s at power 6) and heated for 5 min at 95°C. 15µl of each sample was then analyzed following the General Western Blot Procedure described above.

### Labeling of MHC Multimer with RFT-Biotin

250 µL of unlabeled MHC Klickmer (MHC Klickmer A*0201/NLVPMVATV/NONE, Immudex: KL-WB2132-NONE, Lot: 20190520-EA1 (pp65 specific)) or (MHC Klickmer A*0201/Neg. Control/NONE, Immudex: KL-WB2666-NONE, Lot: 20190607-PL2) was aliquoted into an Eppendorf tube followed by addition of 550 nM RFT biotin from a 90 µM DMSO stock solution (prepared as described below). The sample was briefly mixed and then diluted 1:1 with assembly buffer and incubated at 4°C for 1hr in the dark. The sample was then stored at 4°C in the dark till use.

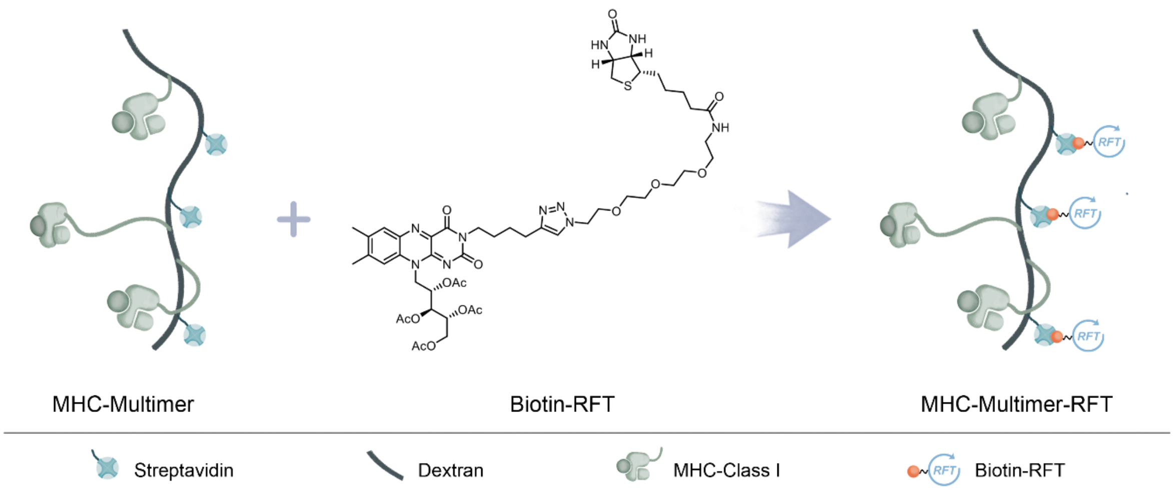

**Supporting Figure 29.** Schematic depicting labeling of MHC-Multimer with Biotin-RFT to form the MHC-Multimer-RFT for the MHC-PhoTag system.

### Selective Biotinylation of CMV specific CD8+ Cells using MHC Multimers

Cyropreserved human CMV-specific CD8+ T cells (Astarte: 1049, Lot:4422AU19) were rapidly thawed in a 37°C water batch and diluted 1:5 in Stain Buffer (BD Pharmingen 554656). The cells were pelleted (400xg, 4min) and the supernatant was discarded. The cells were resuspended at 500,000 cells/mL in Stain Buffer aliquoted in 1 mL aliquots to Eppendorf tubes. The cells were spun at 400xg for 4min and resuspended in 50 µL of Stain Buffer followed by addition of 15 µL of MHC-Multimer (MHC Klickmer HLA-A*0201/NLVPMVATV/NONE, Immudex: KL-WB2132-NONE, Lot: 20190520-EA1 (pp65 specific)) or (MHC Klickmer HLA-A*0201/Neg. Control/NONE, Immudex: KL-WB2666-NONE, Lot: 20190607-PL2) that was pre-labeled with RFT-biotin. The samples were incubated for 30 min on ice and washed 1x with 0.5 mL of cold Stain Buffer and 1x with cold PBS. The cells were resuspended in 200 µL of PBS containing 2 µM biotin tyramide and irradiated in the bio-photoreactor for the indicated time points. The samples were removed and washed 2x with 0.5 mL of cold Stain Buffer and then resuspended in 100 µL of cold Stain Buffer followed by addition of 5 µL of anti-CD8 APC (BioLegend: 344722), and 1 µL strep PE (BD Biosciences: 554061). The cells were incubated for 20 minutes on ice and then washed 1x with cold Stain Buffer and 1x with cold PBS. The cells were resuspended in 200 µL of PBS and transferred to 5 mL FACS tubes (Fisherbrand: 14-956-3D) and acquired samples on a BD FACSCelesta (Model No: 660344) with BD FACSDiva software (v8.0.1.1). Data was analyzed using FlowJo v10 (FlowJo, LLC).

### MHC Multimer Flavin-Based Biotinylation and Sorting of CMV Specific CD8+ Cells from Human Donor PBMCs

Cryopreserved human PBMCs from (Astarte: 1001, Lot# 4444SE19 or 4487OC19) were rapidly thawed in a 37°C water batch and diluted 1:5 in Stain Buffer (BD Pharmingen 554656). The cells were pelleted (400xg, 4min) and the supernatant was discarded. The cells were resuspended at 20-30 million cells/mL in Stain Buffer and aliquoted in 1 mL aliquots to Eppendorf tubes. The cells were spun at 400xg for 4min and resuspended in 50 µL of Stain Buffer followed by addition of 15 µL of MHC Multimer (MHC Klickmer HLA-A*0201/NLVPMVATV/NONE, Immudex: KL-WB2132-NONE, Lot: 20190520-EA1 (pp65 specific)) or (MHC Klickmer HLA-A*02:01/Neg. Control/NONE, Immudex: KL-WB2666-NONE, Lot: 20190607-PL2) that was pre-labeled with RFT-biotin. The samples were incubated for 30 min on ice and washed 1x with 1 mL of cold Stain Buffer and 1x with cold PBS. The cells were resuspended in 200 µL of PBS containing 2 µM biotin tyramide and irradiated in the bio-photoreactor for the indicated time points. The samples were removed and washed 2x with 1 mL of cold Stain Buffer and then resuspended in 100 µL of cold Stain Buffer followed by addition of 5 µL of anti-CD8 APC (BioLegend: 344722), and 1 µL strep PE (BD Biosciences: 554061). The cells were incubated for 20 minutes on ice and then washed 1x with 1mL cold Stain Buffer and 1x with 1mL cold PBS. The cells were then resuspended in 100 µL of PBS containing Zombie Violet viability stain (BioLegend: 423113) (diluted 1:500 in PBS) and incubated for 15 minutes at room temperature. The cells were washed 1x with 1mL PBS and then resuspended in either PBS for flow cytometry analysis or in FACS sorting buffer (PBS with 2% BSA) and transferred to 5 mL FACS tubes (Fisherbrand: 14-956-3D) and acquired samples on a BD FACSCelesta with BD FACSDiva software. Data was analyzed using FlowJo v10 (FlowJo, LLC). Cell sorting was performed using a BD FACSMelody with BD FACSChorus software. Additional data analysis was done using FlowJo v10. After cell sorting, samples were pelleted by centrifugation (400xg, 4min) and resuspended in FACS sorting buffer at 1,000 cells/µL and submitted for TCR sequencing analysis. Prior to sample submission, 10 µL of each sorted sample was analyzed by flow cytometry to determine final cell population purity.

### MHC Multimer PE Staining and Sorting of CMV Specific CD8+ Cells from Human Donor PBMCs

Cryopreserved human PBMCs from (Astarte: 1001, Lot: 4444SE19 or 4487OC19) were rapidly thawed in a 37°C water bath and diluted 1:5 in Stain Buffer (BD Pharmingen: 554656). The cells were pelleted (400xg, 4min) and the supernatant was discarded. The cells were resuspended at 20-30 million cells/mL in Stain Buffer and aliquoted in 1 mL aliquots to Eppendorf tubes. The cells were spun at 400xg for 4min and resuspended in 50 µL of Stain Buffer followed by addition of 15 µL of Phycoerythrin (PE)-Labeled MHC Multimer (MHC Dextramer HLA-A*02:01/NLVPMVATV/PE, Immudex: WB2132-PE, Lot: 20190513-EA8 (pp65 specific)) or (MHC Klickmer A*02:01/Neg. Control/PE, Immudex: WB2666-PE, Lot: 20190520-PL3). The samples were incubated for 10 min on ice followed by addition of 5 µL of anti-CD8 APC (BioLegend: 344722) to each sample. The samples were incubated for another 20 min on ice. After incubation, the samples were pelleted by centrifugation (400xg, 4min) and washed 1x with 1 mL of cold Stain Buffer and

1x with 1 mL cold PBS. The samples were then resuspended in 100 µL of PBS containing Zombie Violet viability stain (BioLegend: 423113) (diluted 1:500 in PBS) and incubated for 15 minutes at room temperature. The cells washed 1x with PBS and then resuspended in either PBS for flow cytometry analysis or in FACS sorting buffer (PBS with 2% BSA) and transferred to 5 mL FACS tubes (Fisherbrand: 14-956-3D) and acquired samples on a BD FACSCelesta with BD FACSDiva software. Data was analyzed using FlowJo v10 (FlowJo, LLC). Cell sorting was performed using a BD FACSMelody with BD FACSChorus software. Additional data analysis was done using FlowJo v10. After cell sorting, samples were pelleted by centrifugation (400xg, 4min) and resuspended in FACS sorting buffer at 1,000 cells/µL and submitted for TCR sequencing analysis. Prior to sample submission, 10 µL of each sorted sample was analyzed by flow cytometry to determine final cell population purity.

#### Single Cell TCR Sequencing and Data Analysis

31.7 µl of the cell suspension (targeting about 5,000 cells) was mixed with 68.3 µl of RT reaction mix and loaded into a Chromium microfluidics chip (10x Genomics). mRNA of each cell was captured by the barcoded bead within each oil droplet, and barcoded cDNA was generated with reverse transcription. The alpha beta TCR repertoire was enriched with the 10x Genomics protocol for Single Cell V(D)J TotalSeqC (feature barcoding). The final PCR products after two rounds of amplifications were purified via SPRI beads, and 50ng of purified samples was carried forward for library construction. The constructed libraries were sequenced in a 150bp-by-150bp paired-end rapid run on an Illumina HiSeq 2500.

Raw bcl files were demultiplexed with bcl2fastq (Illumina) and were then processed with CellRanger vdj (v3.0.2, 10x Genomics) to assemble TCR sequences. Full-length, productive and paired ab TCR sequences of each single cell were retained for downstream analysis. We filtered out singletons and examined the distributions of paired clonotypes (number of cells >3) across donors and treatments in R v3.6.0. We looked up the antigen specificity of each alpha and beta chain TCR clonotype based on their amino acid sequences of the CDR3 region in VDJdb (https://vdjdb.cdr3.net) and reported the most probable epitope of each TCR and their informativeness scores.

#### Photocatalytic Cell Tagging (PhoTag) in Two-Cell System

For transcellular labeling using the primary/secondary antibody flavin conjugate system, 10 million Jurkat PD-1 cells and 10 million Raji PD-L1 cells per sample were centrifuged for 4 min at 800xg, 4°C in Protein LoBind tubes, and each cell pellet was resuspended in its respective complete media. 5µg of Mouse α-Human CD45RO (BD Pharmingen, clone UCHL1: 562641) was added to Jurkat-PD-1 cells and 5µg of Mouse α-Human CD274 (PD-L1), clone MIH1 (Invitrogen: 14-5983-82) was added to Raji PD-L1 cells. 5µg of isotype control (BD Pharmingen, Purified Mouse IgG1 κ, clone MOPC-21: 556648) was added to the respective Jurkat PD-1 and Raji PD-L1 cells. Samples were incubated on a rotisserie at 4°C for 30 min followed by 1 wash with 1ml of cold complete media (cells centrifuged at 800xg for 4 min at 4°C to pellet cells). Cell pellets were then resuspended in 1ml of cold complete media, and 5µg of Goat α-Mouse IgG Riboflavin tetraacetate was added to each sample. All cells were incubated at 4°C for 30 min on the rotisserie and then washed twice with 1ml of cold 1x DPBS (cells centrifuged at 800xg for 4 min at 4°C to pellet cells).

For transcellular labeling using peroxidase, 10 million cells per sample were resuspended in 500µl of cold complete media for each cell line (see General Materials section for cell line growth media) in Protein LoBind tubes containing 5µg of Purified Mouse α-Human CD45RO (BD Pharmingen, clone UCHL1: 562641), Purified Mouse α-Human PD-L1 (Thermo Fisher Scientific, clone MIH1: 14598382), or Isotype control (BD Biosciences, Purified Mouse IgG1 κ, clone MOPC-21: 556648) and incubated for 30 min in a rotisserie at

4°C. The cells were pelleted at 800xg for 4 min at 4°C, washed once in cold complete media and resuspended in 500µl of complete media containing 5µg of Goat α-Mouse IgG Horseradish Peroxidase (Millipore, AP124P) and incubated for 30 min in a rotisserie at 4°C. The cells were pelleted at 800xg for 4 min at 4°C, washed twice with cold 1x DPBS and the cell pellet resuspended in 500µl of cold 1x DPBS.

For transcellular labeling using α-PD-L1 VHH-Fc-RFT conjugate, 10 million Raji PD-L1, JY PD-L1 or PD-L1 aAPC/CHO-K1 cells per sample were centrifuged for 4 min at 800xg, 4°C in Protein LoBind microcentrifuge tubes, and resuspended in 500µl of cold complete media. 5µg of Isotype VHH-Fc-RFT or α-PD-L1 VHH-Fc-RFT conjugate was added to the cells and incubated on a rotisserie at 4°C for 1 hour, followed by 2 washes with 1ml of cold 1x DPBS with pelleting at 800xg for 4 min at 4°C in between washes.

120 ng/mL of Staphylococcal Enterotoxin D (SED) (Toxin Technology: DT303) was added to all Raji or JY cells and incubated for 30 min on ice. No SED was added to PD-L1 aAPC/CHO-K1 cells. After incubation, 1 million Raji or JY cells were combined with 1 million Jurkat PD-1 cells, or 1 million Raji cells were combined with 1 million non-binding cells (suspended A375 cells or Jurkat ΔCD3/TCRαβ cells (JRT3-T3.5) with or without co-pelleting at 800xg for 4 min at 4°C 1 million PD-L1 aAPC/CHO-K1 were combined with 1 million Jurkat PD-1 Effector cells (Promega: J1252). Combined cells were resuspended in a final volume of 45µl of 1x DPBS and incubated for 2.5 hours at 37°C + 5% $CO_2$ or for shorter time points as highlighted in Supporting Figure 31.

For labeling of co-cultured cells using antibody/VHH-targeted RFT, a 25mM biotin tyramide stock in DMSO was diluted 1:10 in 1x DPBS and added at a final concentration of 250µM with gentle mixing, followed by irradiation for 2 min at 100% light intensity in the bio-photoreactor. "No light" control samples were kept in the dark for 2 min. After irradiation, cells were centrifuged for 4 min at 800xg, 4°C and washed once with 1ml of cold 1x DPBS.

For HRP labeled samples, reaction buffer (250 µM biotin tyramide and 1 mM $H_2O_2$ in cold 1x DPBS) was added to each sample and incubated for 1 min at 4°C. After 1 min, 50 µL of quenching buffer (cold 1x DPBS containing 5 mM Trolox, 10 mM Sodium Ascorbic Acid, 10 mM $NaN_3$) was added to the cells. The cells were then pelleted by centrifugation (4 min at 1,000xg, 4°C), washed again with 50 µL of quenching buffer and pelleted. Cells were washed once in 1 mL of cold 1x DPBS, pelleted and resuspended in 500 µL of 1x DPBS until ready to be processed for flow cytometry or microscopy analysis.

#### Flow Cytometry Analysis of Cell Tagging in Two-Cell System

After labeling, cells were pelleted by centrifugation (4 min at 1,000xg, 4°C) and resuspended in 100 µL of Stain Buffer (BD Biosciences: 554656) and transferred to a 96-well V-bottom plate (Falcon: 3894) containing a 1:100 dilution of Fc block (BD Biosciences: 564220). The cells were incubated for 20 min on ice then centrifuged for 4 min at 500xg, 4°C, washed once with 200 µL Stain Buffer and centrifuged again as above. Cells were resuspended in 100 µL of Stain Buffer containing the following antibodies at the indicated dilutions: Alexa Fluor 488 Mouse α-Human CD3, clone SP34-2 at a 1:40 dilution (BD Biosciences: 557705), APC Mouse α-Human CD19, clone HIB19 at a 1:10 dilution (BD Biosciences: 555415), and Streptavidin PE at a 1:200 dilution (BD Biosciences: 554061). Compensation controls were generated using UltraComp beads (BD Biosciences: 01-2222-42), BV421 Mouse α-Human CD45RO, clone UCHL1 (BD Biosciences: 562641), PE Mouse α-Human CD3, clone SK7 (BioLegend: 344806), Alexa Fluor 488 Mouse α-Human CD3, and APC Mouse α-Human CD19, clone HIB19. The plate was incubated for 30 min on ice, the cells were washed once with 200 µL of cold 1x DPBS then resuspended in a 1:500 dilution of Zombie Violet

Viability kit (BioLegend: 423113) and incubated for 15 min at room temperature, protected from light. The cells were centrifuged as above and resuspended in 200 µL of cold 1x DPBS and centrifuged again. The final pellets were resuspended in 200 µL of Stain Buffer and transferred to 5mL FACS tubes (Fisherbrand: 14-956-3D) and acquired samples on a BD FACSCelesta (Model No: 660344) with BD FACSDiva software (v8.0.1.1). Data was analyzed using FlowJo v10 (FlowJo, LLC).

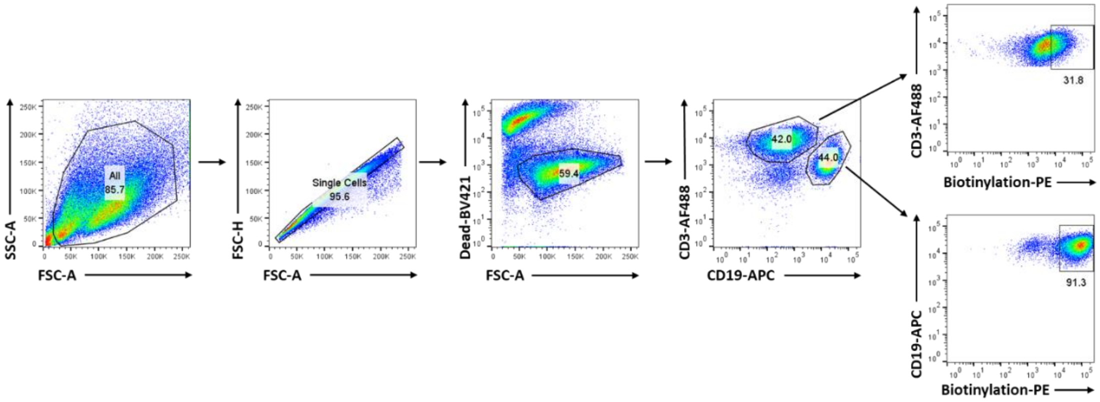

**Supporting Figure 30. Gating strategy for flow cytometry analysis of Jurkat-Raji two cell system.** Briefly, cells were gated for SSC-A v. FSC-A, followed by single cell gating using FSC-H v. FSC-A. Cells were then selected for viability using Dead-BV421 v. FSC-A, followed by separating T and B cell population by gating on CD3-Alexa Fluor 488 (AF488) v. CD19-APC. Biotinylation of T and B cells was evaluated by gating on CD3-AF488 or CD19-APC v. Biotinylation (Streptavidin)-PE.

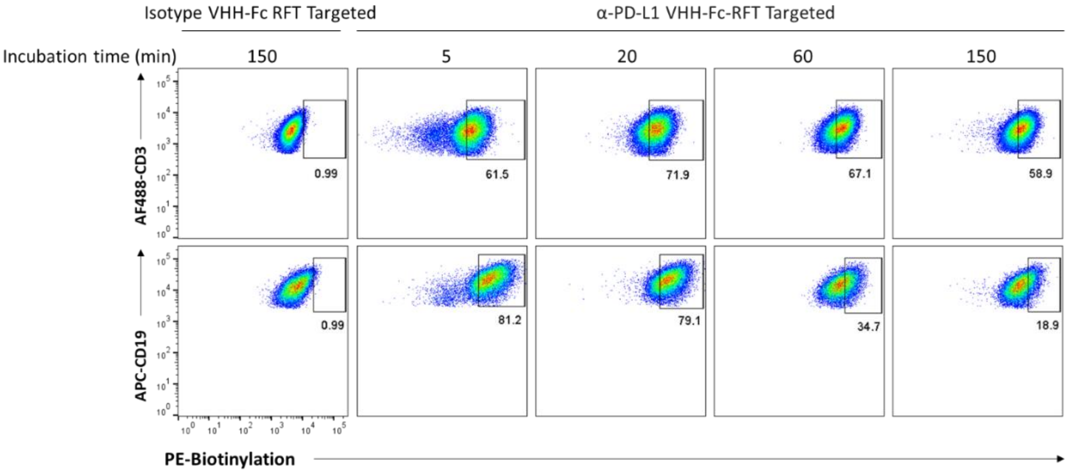

**Supporting Figure 31. Transcellular biotinylation at different cell-cell incubation time points.** a) Raji PD-L1 cells (CD19+) were pre-labeled with α-PD-L1 VHH-Fc-RFT or isotype control (Isotype VHH-Fc-RFT) and co-cultured with Jurkat PD-1 cells (CD3+) in the presence of staphylococcal enterotoxin D (SED) superantigen for 0, 5, 20, 60, or 150 minutes followed by visible light irradiation in the presence of biotin tyramide for 2 minutes. Flow cytometry analysis shows similar levels of both *cis*- and *trans*-biotinylation over the different cell-cell incubation times. Data is representative of two independent experiments.

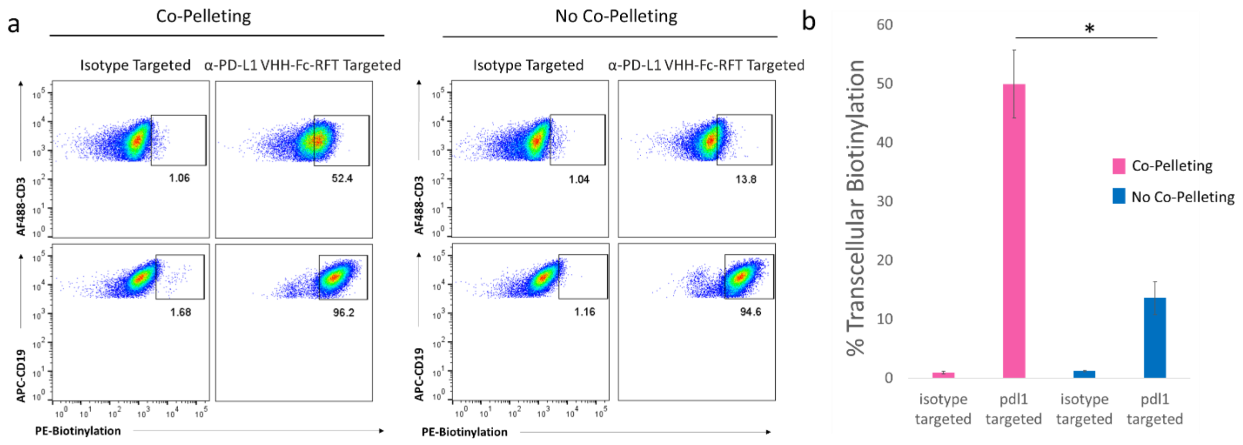

**Supporting Figure 32. Effect of co-pelleting on translabeling.** a) Raji PD-L1 cells (CD19+) were pre-labeled with α-PD-L1 VHH-Fc-RFT or isotype control (Isotype VHH-Fc-RFT) and co-cultured with Jurkat PD-1 cells (CD3+) in the presence of staphylococcal enterotoxin D (SED) superantigen and immediately irradiated with visible light in the presence of biotin tyramide for 2 minutes (no co-pelleting) or pelleted by centrifugation to induce cell-cell contact, gently resuspended and irradiated with visible light irradiation in the presence of biotin tyramide for 2 minutes (co-pelleting). Flow cytometry analysis shows significantly increased levels of transcellular biotinylation in co-pelleted cells. b) Bar plots of replicate analysis of cell biotinylation measured by flow cytometry in panel a for Jurkat PD-1 cells. Error bars represent standard deviation of n = 3 experiments; *$P < 0.01$.

### Confocal Microscopy Imaging of Cell Tagging in Two-Cell System

Microscope coverslips (Fisherbrand, 12-545-81) were acid-etched in 1N HCl (Fisher, SA56-1) for 30 min at 50°C, followed by 3x washes in distilled water then stored in 100% ethanol (Fisher, BP2818-500) at room temperature. The coverslips were placed into a 24-well plate (Thermo Fisher Scientific, 142485), and washed 2x with 1 mL of 1x DPBS (Gibco, 14190-144). 0.5 mL of poly-L-lysine solution (Sigma: P4707-50ML) was added per well and incubated for 30 min at 37°C. The coverslips were washed 2x with 1 mL of 1x DPBS. Approximately 2 million labeled cells using the photocatalytic cell tagging or peroxidase method were loaded in 500 µL of 1x DPBS per well on the 24-well plate. The cells were centrifuged at 400xg for 3 min with deceleration set at 3 using a Sorvall Legend XTR table-top centrifuge (Thermo Scientific).

Samples were incubated in a 3% PFA (Ted Pella: 18505-100) and 0.1% glutaraldehyde (Sigma-Aldrich: G5882-10X10ML) fixative mixture in 1x DPBS for 10 min at 4°C to adhere the cells to the coverslips. The coverslips were then washed 3x in Stain Buffer (BD Biosciences: 554656) and incubated overnight in 1 mL of Stain Buffer at 4°C to block. The following day, the monoculture samples were stained with 500 µL of streptavidin, Alexa Fluor 647 conjugate antibody (Invitrogen: S32357) at a 1:200 dilution in Stain Buffer. For transcellular labeling, the samples were stained with Alexa Fluor 488 streptavidin (BioLegend: 405235) at a 1:200 dilution in 500µl of Stain Buffer. The plate was sealed and stored overnight at 4°C, protected from light. The coverslips were washed 1x in 500µl of Stain Buffer and Hoechst DNA dye (Cayman Chemical Company: 600332) was added at a 1:10,000 dilution in 500µl of Stain Buffer, followed byincubation for 10 min at room temperature, protected from light. The coverslips were washed 2x in

Stain Buffer and fixedwith 500µl of 3% PFA and 0.1% Glutaraldehyde solution in 1x DPBS for 5 min at room temperature. The coverslips were washed 2x in Stain Buffer. ProLong Gold Anti-fade mountant (Invitrogen: P36934) was added to each microscope slide (J. Melvin Freed Brand: 301MF, frosted, 3 x 1") and each coverslip was placed on top of mountant on their respective slides and dried overnight at room temperature, protected from light. Imaging was performed using a Zeiss LSM800 inverted, confocal microscope with a 63X oil immersion objective and Airyscan settings.

### IL-2 Measurement in Two-Cell System

45 µL of a 8e6 cells/mL stock of Jurkat PD-1 cells in assay media (RPMI 1640, Corning: 10-040-CV + 10% dialyzed FBS, HyClone: SH30079.03) were mixed with 45 µL of vehicle control, or Isotype VHH-Fc-RFT (1:3 VHH-Fc to RFT), α-PD-L1 VHH-Fc-RFT (1:3 VHH-Fc to RFT), mouse IgG1 κ isotype control antibody, clone MOPC-21 (BD Biosciences: 556648), or α-human PD-L1 antibody, clone MIH1 (Invitrogen: 14-5983-82) at a final concentration of 10 µg/mL in assay medium. Cells wereincubated for 30 min at room temperature in a 96-well U-bottom plate. A final concentration of 120 ng/mL of Staphylococcal Enterotoxin D (SED) (Toxin Technology: DT303) in assay medium was added to a 8e5 cells/mL stock of Raji PD-L1 cells in assay medium. The cells wereincubated for 30 min at 37°C + 5% $CO_2$.

125 µL of SED-loaded Raji PD-L1 cells were aliquoted per well in a separate 96-well U-bottom plate and mixed with 25 µL of the antibody-loaded Jurkat PD-1 cells (for 100,000 cells per cell line) andincubated for 24 hours at 37°C + 5% $CO_2$. The cells were then pelleted for 1 min at 1,100 RPM and room temperature. 50 µL of culture supernatant per well were used per Human IL-2 ELISA kit manufacturer instructions (Thermo Fisher: EH2IL25). 550 nm absorbance values were subtracted from 450 nm absorbance values to obtain IL-2 readout. Data analysis and graph generation using MS Excel and GraphPad Prism software (8.1.1.330).

### Sorting of Biotinylated Jurkat Cells from Two-Cell System Synaptic Labeling

After cell labeling (see Photocatalytic Cell Tagging (PhoTag) in Two-Cell System), cells were stained for flow cytometry (see Flow Cytometry Analysis of Cell Tagging (PhoTag) in Two-Cell System) and acquired on a BD FACSMelody with BD FACSChorus software by gating on total CD3+ cells, CD3+ biotin high or CD3+ biotin low cells relative to isotype-targeted controls. 250,000 cells were sorted per sample. Additional data analysis was done using FlowJo v10. Sorted cells were pelleted by centrifugation (400xg, 4min), flash frozen in dry ice and stored at -80°C before submitting samples for bulk RNA sequencing analysis.

### RNA-Seq Analysis of from Two-Cell System Synaptic Labeling

**Sequencing**

Samples were provided to Q2 solutions for sequencing on the HiSeq2500. Library preparation was performed using the TruSeq stranded mRNA kit to generate 50bp paired end reads with a library depth of approximately 40 million reads per sample. Fastq files were provided by Q2 and were mapped to the human genome version hg38. Mapped reads were quantified using the Ensembl version 93 gene annotation file to create a gene count matrix.

After mapping, quality control metrics such as boxplots, IQR and Median measurements, PCA plots, and library size measurements indicated that all samples were well within an expected range of < 3 standard deviations from mean values, so no sample removal was performed.

**Differential Expression Analysis**

Genes were filtered to remove any gene which did not reach at least 1 read count per million in at least 4 samples and Ensembl IDs known to belong to long non-coding RNAs and microRNAs. Differential expression was then performed using the program EdgeR[10,11]. This program utilizes an overdispersed Poisson model which accounts for both biological and technical variability coupled with an empirical Bayes method to moderate the degree of overdispersion. Library size normalization factors were generated using the TMM method (trimmed mean of M-values) from the calcNormFactors function in the edgeR library to be used as covariates in modeling. Data was then fit to a generalized linear model (GLM) and a GLM likelihood ratio test was performed to determine if the coefficient representing the contrast between the conditions of interest was equal to 0 indicating no differential expression. Significance for differential expression was set at an FDR corrected p-value < 0.05.

Differentially expressed genes were then subjected to gene set enrichment analysis using toppgene (https://toppgene.cchmc.org/) to identify genes belonging to immune related categories. Specifically, adaptive and innate immune response, and cell adhesion. Other genes not belonging to those categories,but played a role in the immune system were noted as "known immune function" in graphical output.

**Generation of Graphics**

Volcano plots were generated in R with the ggplot library[12]. Log2FC and p-value estimates from EdgeR were subset to those reaching a log2FC of 1 and an FDR corrected p-value of 0.05. Proteins were colored based on the category described in the figure legend.

Heatmaps were generated using the heatmap.2 function in the gplots library[13]. Dendrograms were created using the Euclidean distance measure with the "complete" clustering algorithm.

Correlation heatmaps were generated using the ComplexHeatmap[14] library in R. Correlation was performed on the list of differentially expressed genes between the biotin high and biotin low conditions after subsetting this list to only genes with an absolute magnitude log2 fold change >0.5 and an FDR corrected p-value <0.05. The distance metric used in clustering was Pearson correlation and the clustering algorithm was "complete".

### VHH generation

An alpaca was immunized with a DNA expression vector encoding human PD-L1. Blood samples were collected, and peripheral blood mononuclear cells prepared using Ficoll-Hypaque according to the manufacturer's instructions (Amersham Biosciences). Total RNA was extracted from the cells and used as starting material for RT-PCR to amplify the VHH/VHH-encoding DNA segments which were then cloned into a phage display vector to create VHH antibody libraries, essentially as described in WO 05/044858. Phage were prepared according to standard protocols and stored after filter sterilization at 4°C for further use. Biotinylated human PD-L1 was used for two rounds of in-solution selection of the VHH library, selected phage plated, and individual phage transferred to a 96 well plate.

Phage-encoded VHH antibodies were expressed in *E. coli* and periplasmic extracts prepared. Primary screening focused on flow cytometry-based binding to CHO cells expressing human PD-L1 (CHO-K1.hPD-L1) or human PD-L2. Secondary screening indicated the presence of PD-L1 binding, but PD-1-non-blocking/CD80-non-blocking VHHs with acceptable off-rates that could be used as a proximity labeling tool. Selected non-blocking PD-L1 VHHs were recloned into the *E. coli* expression vector with a C-terminal $FLAG_3$-$HIS_6$ tag. Electrocompetent cells were transformed, positive clones were sequenced, shake flask cultures started, and VHH purified via IMAC purification.

**Binding to human PD-L1**

CHO-K1.hPD-L1 cells were resuspended in Stain Buffer (PBS, 2% FBS, 0.05% sodium azide) and 1E5 cells/well were transferred to a 96-well V-bottom plate and centrifuged. Cells were suspended in 100µL/well serial diluted VHH or antibody in Stain Buffer and incubated for 30 minutes at 4°C. VHH binding was detected by resuspending the samples subsequently in 100µL/well mouse a-FLAG antibody (Sigma: F1804) and detected with PE-labeled goat anti-mouse IgG (Jackson ImmunoResearch: 115-116-071) as detection antibody and 5nM TOPRO3 (Molecular Probes: T3605) as dead dye. Between each step, the cells were collected via centrifugation for 5 minutes at 200xg and washed three times with 100µL/well Stain Buffer.

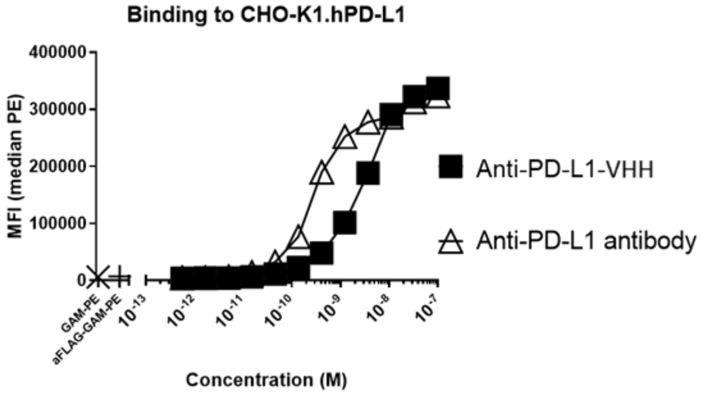

**Supporting Figure 33.** Binding of α-PD-L1 VHH to CHO-K1.hPD-L1 expression cells.

**Blockade of human PD-1 binding to human PD-L1**

CHO-K1.hPD-L1 cells were resuspended in Stain Buffer (PBS, 2% FBS, 0.05% sodium azide) and 1E5 cells/well were transferred to 96-well V-bottom plates and centrifuged. Cells were suspended in 100µL mixture of serially diluted VHH (or antibody) and 9.3 nM human PD-1-Fc in Stain Buffer and incubated for 1.5 hours at 4°C, 300rpm. Residual binding of human PD-1-Fc was detected with 100µL PE-labeled goat anti-human IgG antibody (Southern Biotech: 2043 09). Between each step, the cells were collected via centrifugation for 5 minutes at 200xg and washed three times with 100µL/well Stain Buffer. Prior to analysis, the samples were resuspended in 5 nM TOPRO3 (Molecular Probes: T3605) to exclude dead cells.

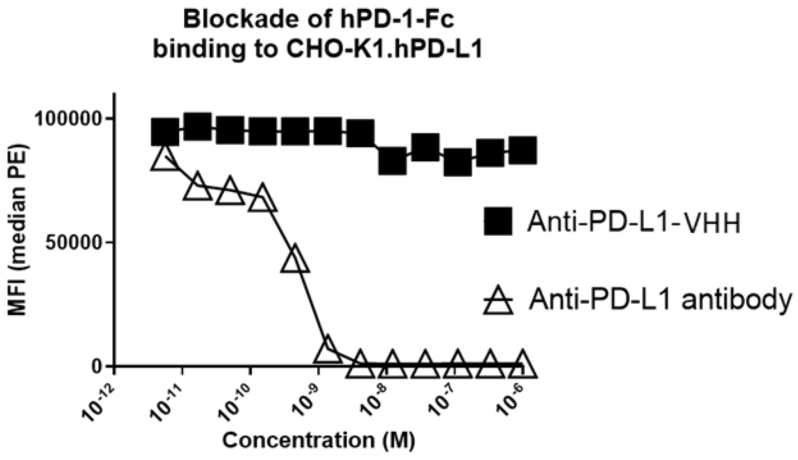

**Supporting Figure 34.** PD-1-Fc mediated blockade of α-PD-L1 VHH binding to CHO-K1.hPD-L1 expression cells.

**Blockade of human CD80 binding to human PD-L1**

An ELISA was performed in a final volume of 25 µL in 384HB Spectraplates (PerkinElmer: 6007500). 2 µg/mL human CD80-Fc (Sino Biological: 10698-H03H) was coated overnight at 4°C. Assay plates were blocked with a 1% casein solution in PBS for at least 1 hour at room temperature. A serial dilution of anti-PD-L1 VHH, anti-PD-L1 antibody, or non-binding VHH control was prepared in assay buffer (PBS, 0.05% Tween-20, 0.1% casein) and mixed with an equal volume of biotinylated hPD-L1-Fc diluted in assay buffer to obtain a final concentration of 50 nM. The samples were added to the appropriate wells and after 1 hour incubation at room temperature, residual binding of biotinylated hPD-L1-Fc was detected with extravidin-HRP (Sigma, Cat E2886). Absorbance at 450 nm was measured with the Tecan Infinite (Tecan) after addition of esTMB substrate (SDT GmbH, Cat esTMB) and HCL (1M).

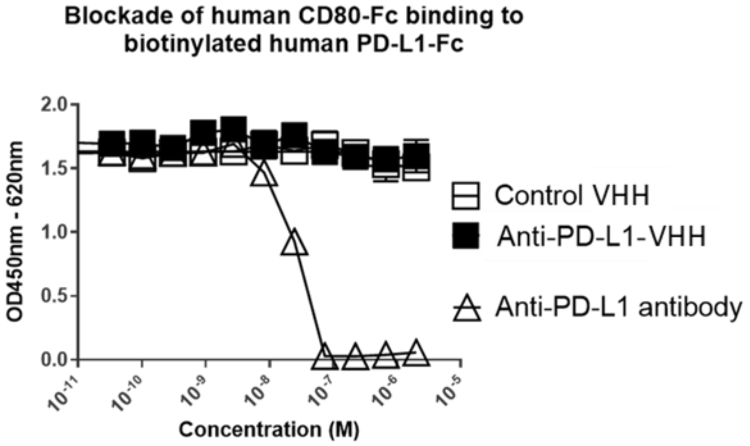

**Supporting Figure 35.** Blockade of human CD80-Fc binding to biotinylated human PD-L1-Fc.

### Preparation of VHH-Fc-$N_3$ Conjugates

**Cloning of Isotype and PD-L1 VHHs**

The DNA encoding the isotype VHH and $\alpha$-PD-L1 VHH, including an N-terminal signal peptide, were purchased from Genewiz (Germany) and fused to the N-terminus of the CH2 domain of the previously described pFUSEN-hG1Fc-IntN[15] vector via overlap extension PCR[16] to obtain the construction Isotype-VHH-Fc-AvaN and PD-L1-VHH-FC-AvaN.

**Expression and purification of Isotype and PD-L1 VHHs:**

Expi293 cells were transiently transfected with either Isotype-VHH-Fc-AvaN or PD-L1-VHH-FC-AvaN constructs using Expifectamine (Life Technology), according to the manufacturer's instructions.

In all cases, after 6 days incubation at 37°C with 8% $CO_2$, cell supernatants were harvested and spun down at 2000 rcf for 20 min at 4°C. After addition of complete protease inhibitors cell supernatants were dialyzed against PBS and directly purified over Protein A column. For Western blot analysis, samples were loaded onto 8 % acrylamide Bis-Tris gels and run in MES-SDS running buffer. Pure fractions were pooled and stored at -80°C.

**Generation of $Npu^C$-Cys-OMe (IntC) protein and Cys-2N3-linker.**

$Npu^C$-Cys-OMe was obtained as previously reported via thiolysis from a Mxe GyrA intein fusion[17].

**Synthesis of Cys-2N3-linker:**

The linker was custom-made at the Pompeu Fabra University Peptide Synthesis Facility (Barcelona, Spain) by standard Fmoc SPPS (solid phase peptide synthesis) on 2-Chlorotrityl resin (0.25 mmol/g, Iris Biotech, Marktredwitz Deustchland). Chain assembly was carried out with HBTU activation (4.8 eq) using a 5-fold excess of amino acid over the resin in DMF (dimethylformamide) with DIEA (N,N- diisopropylethylamine). The Fmoc protecting group was removed with 20% piperidine in DMF (1 x 2 minutes, followed by 2 x 10 minutes). Peptidyl-resin was washed between coupling cycles with DMF for 3 minutes by alternating batch or flow washes. Peptide was cleaved from the resin using 94% TFA, 1% triisopropylsilane (TIS), 2.5% ethanedithiol, and 2.5% $H_2O$ (cleavage cocktail). Crude peptide products were precipitated and washed with cold $Et_2O$, dissolved in solvent A (0.1 % TFA in water) with a minimal amount of solvent B (0.1% TFA 90% acetonitrile in water) and then purified by RP-HPLC.

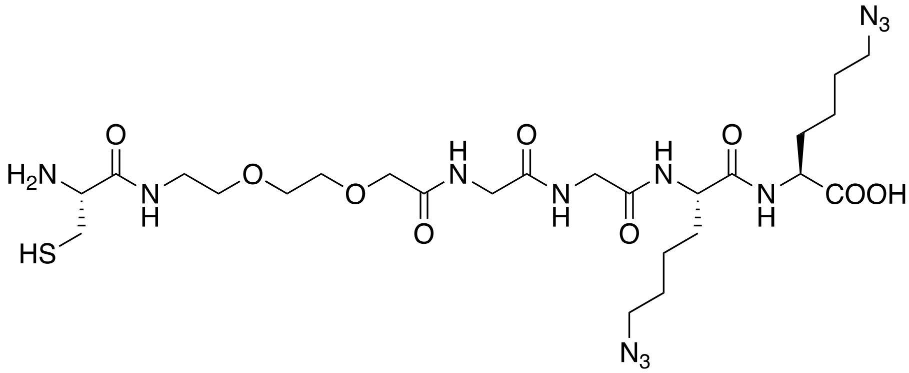

**Supporting Figure 36.** Chemical structure of Cys-2N3 linker compound (Cys-2N3).

#### SEPL reaction of Cys-2N3-linker:

Purified Isotype-VHH-Fc-AvaN or PD-L1-VHH-FC-AvaN constructs were concentrated down to 1 mg/ml. The ligation reaction was initiated by addition of 0.5 mM Cys-2N3 linker, 2 eq. of IntC, 100 mM of MESNa and 0.25 mM TCEP and adjusting pH to 7.5-8.0. The reaction was incubated in the dark at r.t. for 24 h and monitored by SDS-PAGE and Coomassie staining. Once the reactions were completed, they were dialyzed into PBS pH 7.4. The resulting VHH products (Isotype-VHH-Fc-2N3, PD-L1-VHH-FC-2N3) were purified by size-exclusion on an S200 column using the AKTA Purifier system. Elution from the column was monitored by UV-Vis absorbance at 280 nm. The elution volumes were in good agreement with the estimated MW. Pure fractions were pooled and concentrated down and finally analyzed by SDS-PAGE under reduced and non-reduce conditions (see figure below). The total overall yield was 40%, including thiolysis, ligation and purification.

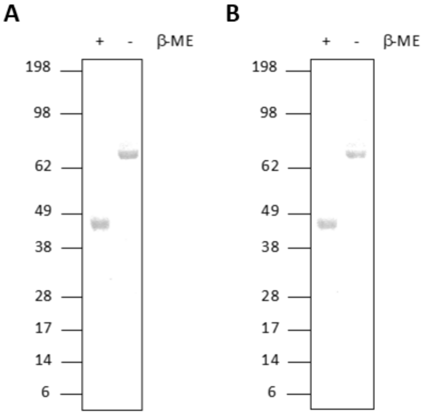

**Supporting Figure 37.** Coomassie stained SDS-PAGE analysis of purified Isotype-VHH-Fc-2N3 (A) and PD-L1-VHH-FC-2N3 (B), under reducing (+ β-ME) and non-reducing (- β-ME) conditions.

#### LC/MS intact mass analysis of conjugates:

Prior to LC/MS analysis, Isotype-VHH-Fc-2N3 and PD-L1-VHH-FC-2N3 were deglycosylated with PNGase F (NEB) under non-denaturing conditions at 37°C overnight. For analysis under reduced conditions, sample was denatured by exchanging buffer to 6 M Gn·HCl in PBS pH 7.4 and treated with 10 mM DTT at 37°C for 1 hr. Deglycosylated and reduced samples were analyzed by RP-HPLC on a Zorbax 300SB C8 column using a 15-70 % linear gradient of solvent C (0.25 % Formic acid and 0.02 % TFA in water) in solvent D (90 % isopropanol in water with 0.25 % Formic acid and 0.02 TFA) over 30 min at 1 mL/min flow rate and 70°C, preceded by a 5 minutes isocratic phase at 15% A. Peaks were collected and analyzed by electrospray ionization mass spectrometric (ESI-MS). ESI-MS analyses were performed on an LCT Premier TOF (Waters) (see Figure below). For analysis under non-reduced conditions, deglycosylated samples were analyzed directly by ESI-MS (see figure below).

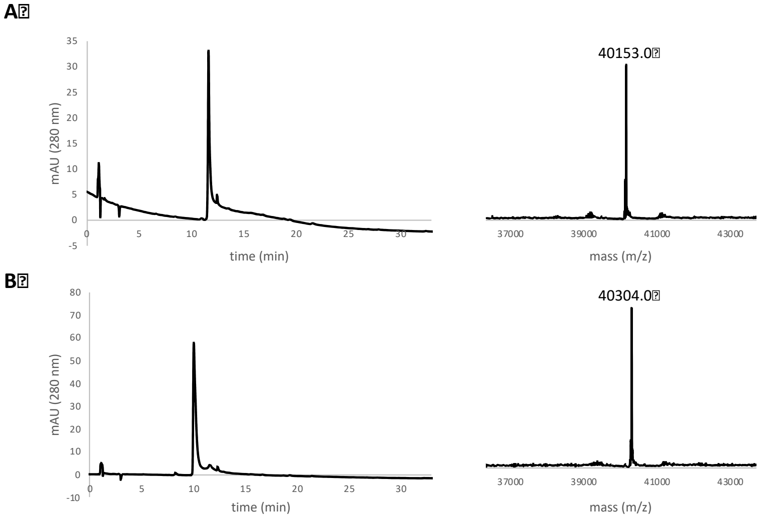

**Supporting Figure 38.** RP-HPLC and ESI-MS analysis under reducing conditions of bis-azido conjugates. Samples were deglycosylated using PNGase F and fully reduced with DTT under denaturing conditions. RP-HPLC analysis was performed over a 15-70%B gradient on a Zorbax 300SB C8 column. A) RP-HPLC (left panel) and ESI-MS (right panel) for Isotype-VHH-Fc-2N3 ($MW_{Obs}$: 40153.0, $MW_{Calc}$: 40155.1), B) RP-HPLC (left panel) and ESI-MS (right panel) for PD-L1-VHH-Fc-2N3 ($MW_{Obs}$: 40304.0, $MW_{Calc}$: 40306.1)

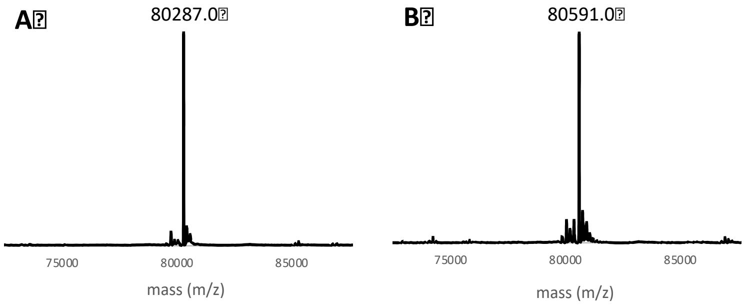

**Supporting Figure 39.** ESI-MS analysis under non-reducing conditions. Samples were deglycosylated using PNGase F and analyzed by LC-ESAI-MS. A) Isotype-VHH-Fc-2N3 ($MW_{Obs}$: 80287.0, $MW_{Calc}$: 80294.2) B) PD-L1-VHH-Fc-2N3 ($MW_{Obs}$: 80591.0, $MW_{Calc}$: 80596.2).

#### Cell Surface Binding with VHH-Fc-488 Conjugates

200 µL of VHH-Fc (1mg/ml in PBS) was combined with 600 µM MB™ 488 DBCO (clickchemistrytools, 1190-5) and incubated for overnight at 4°C. The labeled VHH-Fc was then added to a Zeba Spin desalting column (Thermo Fisher Scientific: 87769, 2 mL column, 40,000 MWCO) and buffer exchanged into PBS, according to manufacturer's instructions. The final protein concentration of the VHH-Fc was measured using the BCA Protein Assay kit according to manufacturer's instructions (Thermo Fisher Scientific: 23227). 488 DBCO concentration was determined by measuring absorbance and comparing to a standard curve of free 488 DBCO of known concentrations. The micromolar concentration of 488 DBCO was divided by the micromolar concentration of VHH-Fc to determine the VHH-Fc:488 fluorophore ratio. The protein concentration was measured by the BCA assay and the photocatalyst concentration was obtained by measuring the absorption compared to free photocatalyst.

PD-L1 VHH-Fc:488 DBCO ratio: 1:3

Isotype VHH-Fc:488 DBCO ratio: 1:3

For cell surface binding, 1 million JY-wt, JY-PD-L1, or Raji-PD-L1 cells were washed with 2x 1ml cold PBS and combined with 2.5 µg Isotype VHH-Fc-488 (Isotype) or α-PD-L1 VHH-Fc-488 conjugate and incubated for 1hr at 4°C on a rotisserie. The cells were washed 2xs with PBS. After washing, the cell pellet was resuspended in 200 µl PBS and transferred to 5ml FACS tubes (Fisherbrand: 14-956-3D) and analyzed on a BD FACSCelesta with BD FACSDiva software. Data was analyzed using FlowJo v10 (FlowJo, LLC).

#### Preparation of Isotype and α-PD-L1 VHH-Fc-RFT Conjugates

200 µl of VHH-Fc (1mg/ml in PBS) was combined with 600uM RFT DBCO and incubated for overnight at 4°C. The labeled VHH-Fc was then added to a Zeba Spin desalting column (Thermo Fisher Scientific: 87769, 2ml column, 40,000 MWCO) and buffer exchanged into PBS, according to manufacturer's instructions. The final protein concentration of the VHH-Fc was measured using the BCA Protein Assay kit according to manufacturer's instructions (Thermo Fisher Scientific: 23227). RFT concentration was determined by measuring absorbance at 450nm (A450) and comparing to a standard curve of free RFT of known concentrations. The micromolar concentration of RFT was divided by the micromolar concentration of VHH-Fc to determine the VHH-Fc:RFT ratio. The protein concentration was measured by the BCA assay and the photocatalyst concentration was obtained by measuring the absorption compared to free photocatalyst.

PD-L1 VHH-Fc:RFT ratio: 1:3

Isotype VHH-Fc:RFT ratio: 1:3

#### Preparation of Isotype and α-PD-L1 VHH-Fc-Peroxidase Conjugates

VHH-Fc proteins were conjugated to horse radish peroxidase (HRP) using an HRP Conjugation Kit (Abcam ab102890) according to manufacturer's instructions. Briefly, 100 µl of VHH-Fc (1mg/ml in PBS) was mixed with 10 µl of modifier reagent and then combined with 100 µg of HRP protein and incubated at rt overnight in the dark. 10 µl of quencher solution was added to each sample and incubated for 30 min. Peroxidase labeled VHH-Fc proteins were stored at 4°C until used in labeling experiments described above.

### General Synthetic Information

Unless specified, all commercially reagents were used as received. Acetonitrile was purchased from EMD Chemicals Inc. (DriSolv, 25 ppm BHT) and used as received. Methanol, dichloromethane, ethyl acetate and heptane were purchased from Fisher Scientific and used as received. The flavin photocatalysts (alloxazine and lumichrome were purchased from Aldrich (A28651 and 103217), lumiflavin was purchased from Santa Cruz Biotechnology and riboflavin was purchased from Fisher) and used as received. Other photocatalysts (Tris(2,2'-bipyridyl)dichlororuthenium(II) hexahydrate, Tris[2-phenylpyridinato-$C^2$,*N*]iridium(III) and Eosin Y) were purchased from Aldrich and used as received.

All NMR spectra were collected on either a Bruker 400 Avance III with a 5 mm BBFO probe (400 MHz for $^1$H; 101 MHz for $^{13}$C) or a Bruker 500 Avance III HD with a 5 mm BBO Nitrogen cryoprobe (500 MHz for $^1$H; 126 MHz for $^{13}$C). The proton signal for non-deuterated solvent (d 7.27 for $CHCl_3$, $\delta$ 2.50 for DMSO) was used as an internal reference for 1H NMR spectra. For $^{13}$C NMR spectra, chemical shifts are reported relative to the $\delta$ 77.00 resonance of $CDCl_3$ or $\delta$ 39.52 resonance of DMSO-*d6*. Deuterated solvents ($CDCl_3$ and DMSO-*d6*) were purchased from Cambridge Isotope Laboratories Inc. and used as received.

HPLC analyses were performed on an Agilent 1260 Infinity II LC system using a 100 mm Agilent Zorbax 300SB-C18 3.5 µm analytical column. Peptide purification performed using semi-preparative HPLC on an Agilent 1260 Infinity II LC system using a Vydac C-18 218TP510 semi-preparative column. Solvent removal was performed using a Benchtop Pro evaporator.

Low-Resolution Mass Spectrometry analyses were conducted on an Agilent 1290 Infinity II LC system (1290 Infinity II multisampler, 1290 Infinity II Binary Pump) with Agilent 6130 Single Quadruple MS and Supelco Ascentis Express C18 column (2.1 mm x 50 mm, 2.7 µm); Column Temperature 50 °C; 0.1% formic acid in water (v/v) as the mobile phase A; 0.1% formic acid in acetonitrile (v/v) as the mobile phase B; 1 mL/min as the flow; ESI+/-, 100-1000 m/z scan, 0.34 sec scan time as the MS method.

High-Resolution Mass Spectrometry analyses were conducted on an Agilent 6545 QTOF mass spectrometer (Agilent Technologies, Santa Clara, CA) in positive or negative electrospray mode. The system was calibrated to greater than 1 ppm accuracy across the mass range prior to analyses according to manufacturer's specifications. The samples were separated using UHPLC on an Agilent 1290 (Agilent Technologies, Santa Clara, CA) system prior to mass spectrometric analysis. The resulting spectra were automatically lockmass corrected and the target mass ions and any confirming adducts (Na+, NH4+) were extracted and combined as a chromatogram. The mass accuracy was calculated for all observed isotopes against the theoretical mass ions derived from the chemical formula using MassHunter software (Agilent Technologies, Santa Clara, CA).

Analytical thin layer chromatography (TLC) was performed on 60 $F_{254}$ glass plates precoated with a 0.25-mm thickness of silica gel purchased from EMD chemical Inc. and TLC plates were visualized with UV light. Column chromatography was performed on TELEDYNE ISCO devices; *CombiFlash® Rf*+ version: 2.0.4.

#### Synthesis of (2*R*,3*S*,4*S*)-5-(3-(hex-5-yn-1-yl)-7,8-dimethyl-2,4-dioxo-3,4-dihydrobenzo[g]pteridin-10(2*H*)-yl)pentane-1,2,3,4-tetrayl tetraacetate (9, Alkyne-RFT)

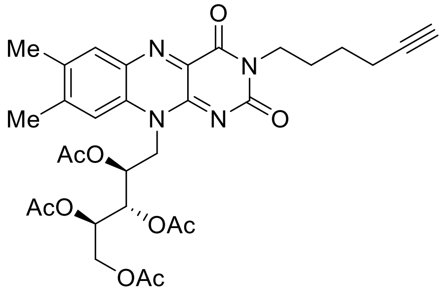

**9, Alkyne-RFT**

A suspension of 2',3',4',5'-tetraacetylriboflavin (1200 mg, 2.20 mmol), potassium carbonate (457 mg, 3.31 mmol), and catalytic amounts of potassium iodide (36.6 mg, 0.22 mmol) in 8 ml of dry *N*,*N*-dimethylformamide was stirred at room temperature for 45 min. Then a solution of 6-iodohex-1-yne (1.16 mL, 8.82 mmol) in 1 ml of dry *N*,*N*-dimethylformamide was added slowly, and the stirring was continued until the 2',3',4',5'-tetraacetylriboflavin was completely alkylated (24 h). The reaction mixture was diluted with 100 ml of dichloromethane, and the organic phase was washed with saturated sodium $NaHCO_3$ solution, water, and brine. Organic solvent was dried over anhydrous $Na_2SO_4$, evaporated, and the remaining residue was purified by column chromatography (EtOAc:DCM; 1:2) to get desired product as an oil (650 mg; 47%).

1H NMR (400 MHz, Chloroform-d) δ 7.98 (s, 1H), 7.51 (s, 1H), 5.64 (d, J = 8.6 Hz, 1H), 5.48 – 5.34 (m, 2H), 4.88 (s, 1H), 4.41 (dd, J = 12.3, 2.5 Hz, 1H), 4.22 (dd, J = 12.3, 5.7 Hz, 1H), 4.07 (t, *J* = 7.3 Hz, 2H), 2.93 (s, 1H), 2.85 (s, 1H), 2.53 (s, 3H), 2.41 (s, 3H), 2.27 (s, 3H), 2.23 (dt, *J* = 7.3, 3.7 Hz, 6H), 2.20 (s, 4H), 2.05 (s, 3H), 1.90 (t, *J* = 2.4 Hz, 1H), 1.81 (p, *J* = 7.6 Hz, 2H), 1.71 (s, 3H), 1.59 (p, *J* = 7.2 Hz, 2H).

$^{13}C$ NMR (101 MHz, $CDCl_3$) δ 170.70, 170.38, 169.94, 169.73, 162.61, 159.74, 155.01, 149.22, 147.55, 136.61, 135.78, 134.74, 132.98, 131.26, 115.45, 84.21, 70.46, 69.08, 68.65, 61.96, 44.53, 41.48, 36.57, 31.51, 27.05, 25.94, 21.50, 21.15, 20.90, 20.79, 20.42, 19.52, 18.29.

HRMS (ESI), m/z: calculated for $C_{31}H_{36}N_4O_{10}$ $[M+H]^+$: 625.2501, found: 625.251.

### Synthesis of DBCO-PEG-Tyramide, 12

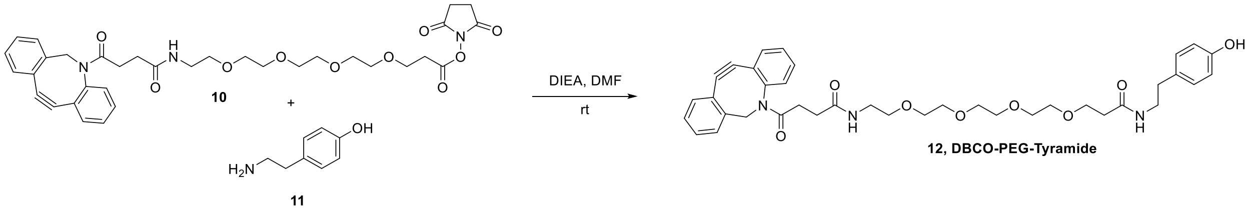

To a solution of tyramine (22 mg, 0.16 mmol) in DMF (1 ml) was added DBCO-PEG-4-NHS ester (100 mg, 0.16 mmol) and DIEA (0.055 ml, 0.31 mmol). The mixture was stirred at room temperature overnight. LC-MS of crude showed product. The mixture was dried with a stream of nitrogen overnight purified on ISCO using DCM: MeOH (100% to 90% to 50%) gave clear gum of the desired product (70 mg, 67%).

$^1$H NMR (400 MHz, Methanol-$d_4$) δ 7.67 (d, *J* = 7.1 Hz, 1H), 7.48 (ddt, *J* = 6.2, 4.0, 2.2 Hz, 1H), 7.45 (d, *J* = 3.2 Hz, 2H), 7.36 (dtd, *J* = 16.1, 7.4, 1.4 Hz, 2H), 7.27 (dd, *J* = 7.4, 1.5 Hz, 1H), 7.04 (d, *J* = 8.4 Hz, 2H), 6.76 – 6.69 (m, 2H), 5.14 (d, *J* = 14.0 Hz, 1H), 3.73 – 3.65 (m, 3H), 3.63 – 3.51 (m, 12H), 3.49 (dd, *J* = 5.5, 2.4 Hz, 2H), 3.36 (t, *J* = 7.3 Hz, 2H), 3.33 (dt, *J* = 3.2, 1.6 Hz, 1H), 3.31 – 3.24 (m, 1H), 3.15 (dt, *J* = 13.7, 7.0 Hz, 1H), 2.70 (t, *J* = 7.3 Hz, 2H), 2.51 (dt, *J* = 16.0, 6.5 Hz, 1H), 2.40 (t, *J* = 6.1 Hz, 2H), 2.28 (t, *J* = 6.2 Hz, 2H), 2.04 (dt, *J* = 15.9, 7.1 Hz, 1H).

$^{13}$C NMR (101 MHz, DMSO) δ 172.25, 169.53, 153.16, 150.22, 149.97, 132.61, 129.41, 128.86, 127.33, 125.43, 118.52, 116.88, 116.03, 115.33, 66.18, 65.84, 54.90, 37.48, 25.97, 22.99.

HRMS (ESI), m/z: calculated for $C_{38}H_{45}N_3O_8$ $[M+H]^+$: 672.3279, found: 672.3287.

### Synthesis of *N*-(2-(4-hydroxy-2-methoxyphenoxy)ethyl)-5-((3aS,4S,6aR)-2-oxohexahydro-1H-thieno[3,4-d]imidazol-4-yl)pentanamide (15, 3MeO-EO-BP):

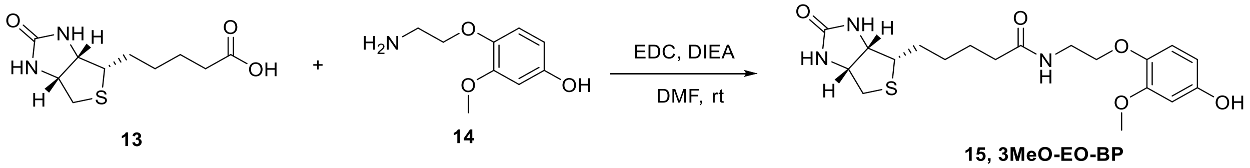

To a solution of 4-(2-aminoethoxy)-3-methoxyphenol (410 mg, 2.23 mmol) in DMF (5 ml) was added EDC (515 mg, 2.69 mmol), biotin (547 mg, 2.23 mmol) and DIEA (1.2 ml, 6.71 mmol). The mixture was stirred at room temperature overnight. The mixture was dried with a stream of nitrogen and purified on ISCO using DCM: MeOH (100% to 90% to 50%) gave yellow solid of the desired product (285 mg, 31%).

$^1$H NMR (400 MHz, DMSO-$d_6$) δ 9.01 (s, 1H), 7.97 (t, *J* = 5.5 Hz, 1H), 6.76 (d, *J* = 8.6 Hz, 1H), 6.42 (s, 1H), 6.40 (d, *J* = 2.7 Hz, 1H), 6.22 (dd, *J* = 8.6, 2.7 Hz, 1H), 4.29 (dd, *J* = 7.6, 4.7 Hz, 1H), 4.10 (dd, *J* = 7.7, 4.4 Hz, 1H), 3.82 (t, *J* = 5.9 Hz, 2H), 3.70 (s, 3H), 3.32 (q, *J* = 5.8 Hz, 2H), 3.11 – 3.02 (m, 1H), 2.81 (dd, *J* = 12.5, 5.1 Hz, 1H), 2.57 (d, *J* = 12.4 Hz, 1H), 2.08 (t, *J* = 7.4 Hz, 2H), 1.60 (m, 1H), 1.55 – 1.39 (m, 3H), 1.30 (m, 2H).

$^{13}C$ NMR (101 MHz, DMSO) δ 162.70, 152.43, 150.44, 140.48, 117.16, 115.65, 106.55, 105.06, 101.64, 100.09, 68.56, 60.20, 59.65, 55.64, 38.41, 36.14, 35.09, 28.21, 25.24.

HRMS (ESI), m/z: calculated for $C_{19}H_{27}N_3O_5S$ $[M+H]^+$: 410.1744, found: 410.1747.

#### Synthesis of *N*-(2-(4-hydroxy-3-methoxyphenoxy)ethyl)-5-((3a*S*,4*S*,6a*R*)-2-oxohexahydro-1*H*-thieno[3,4-d]imidazol-4-yl)pentanamide (18, 2MeO-EO-BP):

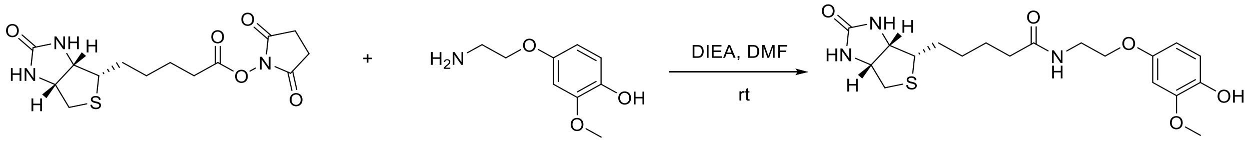

16 17 18, 2MeO-EO-BP

To a solution of 4-(2-aminoethoxy)-2-methoxyphenol (200 mg, 0.91 mmol) in DMF (3 ml) was added BIOTIN-NHS ester (311 mg, 0.91 mmol) and DIEA (0.476 ml, 2.73 mmol). The mixture was stirred at room temperature overnight. LCMS of crude showed product. The mixture was dried with a stream of nitrogen and purified on ISCO using DCM: MeOH (100% to 90% to 50%) gave white solid of the desired product (310 mg, 83%).

$^{1}H$ NMR (400 MHz, DMSO-$d_6$) δ 8.42 (s, 1H), 8.01 (t, *J* = 5.5 Hz, 1H), 6.65 (d, *J* = 8.6 Hz, 1H), 6.53 (d, *J* = 2.8 Hz, 1H), 6.42 (s, 1H), 6.35 (s, 1H), 6.32 (dd, *J* = 8.6, 2.8 Hz, 1H), 4.28 (dd, *J* = 7.5, 5.2 Hz, 1H), 4.13 – 4.05 (m, 1H), 3.87 (t, *J* = 5.7 Hz, 2H), 3.73 (s, 3H), 3.39 – 3.34 (m, 2H), 3.08 – 3.00 (m, 1H), 2.80 (dd, *J* = 12.4, 5.1 Hz, 1H), 2.56 (d, *J* = 12.4 Hz, 1H), 2.09 (t, *J* = 7.4 Hz, 2H), 1.53 (dtdd, *J* = 42.7, 28.4, 12.0, 6.2 Hz, 4H), 1.30 (tt, *J* = 18.5, 9.1 Hz, 2H).

$^{13}C$ NMR (101 MHz, DMSO) δ 172.35, 162.70, 151.66, 148.14, 140.52, 115.34, 105.17, 100.87, 66.81, 61.00, 59.17, 55.56, 38.25, 35.07, 28.02, 25.26.

HRMS (ESI), m/z: calculated for $C_{19}H_{27}N_3O_5S$ $[M+H]^+$: 410.1744, found: 410.1746.

#### Synthesis of *N*-(2-(4-hydroxyphenoxy)ethyl)-5-((3a*S*,4*S*,6a*R*)-2-oxohexahydro-1H-thieno[3,4-d]imidazol-4-yl)pentanamide (20, EO-BP):

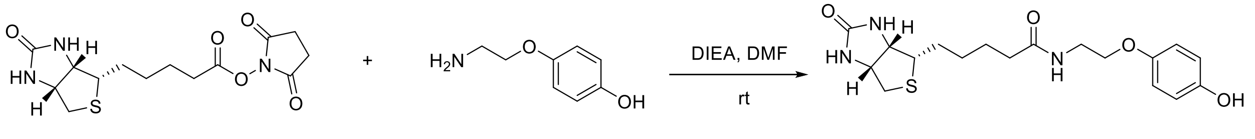

16 19 20, EO-BP

To a solution of 4-(2-aminoethoxy)phenol (140 mg, 0.73 mmol) in DMF (2 ml) was added BIOTIN-NHS ester (252 mg, 0.73 mmol) and DIEA (0.4 ml, 2.21 mmol). The mixture was stirred at room temperature overnight. LCMS of crude showed product. The mixture was dried with a stream of nitrogen and purified on ISCO using DCM: MeOH (100% to 90% to 50%) to give yellow solid of the desired product (275 mg, 98%).

$^{1}H$ NMR (400 MHz, DMSO-$d_6$) δ 8.01 (t, *J* = 5.5 Hz, 1H), 6.77 – 6.71 (m, 2H), 6.68 – 6.62 (m, 2H), 6.42 (s, 1H), 4.29 (dd, *J* = 7.6, 4.6 Hz, 1H), 4.09 (dd, *J* = 7.7, 4.4 Hz, 1H), 3.85 (t, *J* = 5.7 Hz, 2H), 3.35 (q, *J* = 5.6 Hz, 2H), 3.08 – 3.00 (m, 1H), 2.80 (dd, *J* = 12.4, 5.1 Hz, 1H), 2.56 (d, *J* = 12.4 Hz, 1H), 2.08 (t, *J* = 7.4 Hz, 2H), 1.51 (ttd, *J* = 28.2, 13.5, 8.8 Hz, 4H), 1.30 (tq, *J* = 15.6, 8.6, 7.3 Hz, 2H).

$^{13}C$ NMR (101 MHz, DMSO) δ 172.33, 162.70, 151.29, 115.49, 66.88, 61.00, 59.18, 55.40, 38.27, 35.06, 28.12, 28.01, 25.25.

HRMS (ESI), m/z: calculated for $C_{18}H_{25}N_3O_4S$ $[M+H]^+$: 380.1639, found: 380.1631.

### Synthesis of *N*-(3-((4-(dimethylamino)phenyl)amino)-3-oxopropyl)-5-((3a*S*,4*S*,6a*R*)-2-oxohexahydro-1H-thieno[3,4-d]imidazol-4-yl)pentanamide (22, Biotin-DMAA):

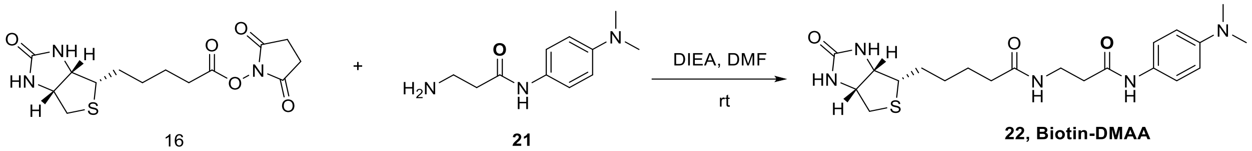

16 + 21 → 22, Biotin-DMAA

To a solution of 3-amino-N-(4-(dimethylamino)phenyl)propanamide-dihydrochloride (140 mg, 0.74 mmol) in DMF (2 ml) was added BIOTIN-NHS ester (252 mg, 0.74 mmol) and DIEA (0.4 mL, 2.21 mmol). The mixture was stirred at room temperature overnight. LC-MS of crude showed product. The mixture was dried with a stream of nitrogen overnight purified on ISCO using DCM: MeOH (100% to 90% to 50%) gave yellow solid of the desired product (256 mg, 54%).

$^{1}H$ NMR (400 MHz, DMSO-$d_6$) δ 11.87 (s, 2H), 10.08 (s, 1H), 7.92 (t, *J* = 5.6 Hz, 1H), 7.66 (d, *J* = 9.0 Hz, 2H), 7.37 (d, *J* = 8.3 Hz, 2H), 6.44 (s, 1H), 4.27 (dd, *J* = 7.7, 4.7 Hz, 1H), 4.08 (dd, *J* = 7.7, 4.4 Hz, 1H), 3.31 (q, *J* = 6.6 Hz, 2H), 3.08 (s, 6H), 3.06 – 3.01 (m, 1H), 2.77 (dd, *J* = 12.5, 5.1 Hz, 1H), 2.58 – 2.53 (m, 1H), 2.47 (d, *J* = 6.8 Hz, 1H), 2.05 (t, *J* = 7.3 Hz, 2H), 1.63 – 1.52 (m, 1H), 1.52 – 1.36 (m, 3H), 1.29 (tp, *J* = 15.3, 8.4, 7.4 Hz, 2H).

$^{13}C$ NMR (101 MHz, DMSO) δ 172.25, 169.61, 162.80, 159.02, 158.66, 158.30, 157.93, 120.17, 119.04, 117.08, 114.18, 61.05, 59.25, 55.43, 44.51, 36.42, 35.14, 28.14, 25.35.

HRMS (ESI), m/z: calculated for $C_{21}H_{31}N_5O_3S$ $[M+H]^+$: 434.2222, found: 434.2222.

#### Synthesis of (2*S*,3*R*,4*R*)-5-(3-(2-(2-((tert-butoxycarbonyl)amino)ethoxy)ethyl)-7,8-dimethyl-2,4-dioxo-3,4-dihydrobenzo[g]pteridin-10(2H)-yl)pentane-1,2,3,4-tetrayl tetraacetate (24)

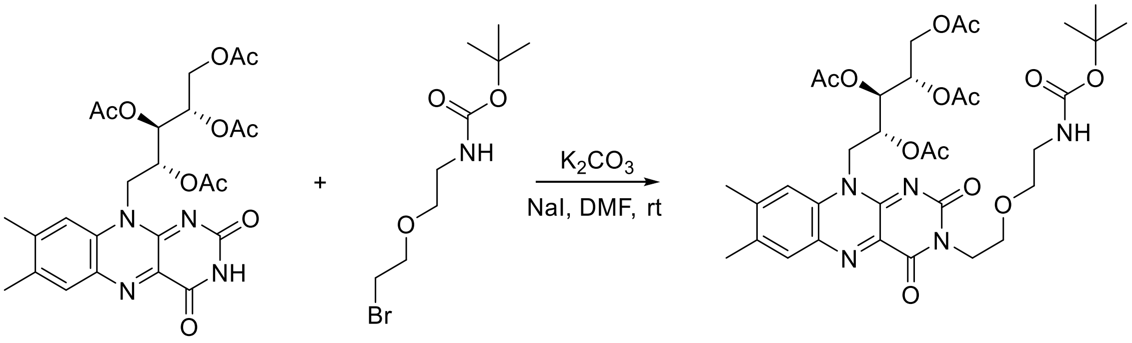

**1, RFT** **23** **24**

A suspension of 2',3',4',5'-tetraacetylriboflavin **1** (1.01 mmol, 550 mg), potassium carbonate (1.515 mmol, 209 mg), and catalytic amounts of sodium iodide in 2 mL of dry *N*,*N*-dimethylformamide was stirred at room temperature for 45 min. Then a solution of *tert*-butyl (2-(2-bromoethoxy)ethyl)carbamate **23** (4.04 mmol, 1083 mg) in 3 mL of of dry N,N-dimethylformamide was added slowly and the stirred for 24 h. Additional 2 equiv of **YY** was added and stirred for another 24 h. The reaction mixture was filtered, dried by stream of nitrogen and purified by column chromatography (DCM : EA 90% to 50% to 100%) to give 120 mg (16% yield) of **24** as orange gum.

$^1$H NMR (400 MHz, DMSO-$d_6$) δ 7.96 (s, 1H), 7.77 (s, 1H), 6.76 (s, 1H), 5.53 – 5.40 (m, 2H), 5.32 (s, 1H), 4.87 (d, *J* = 13.8 Hz, 1H), 4.38 (d, *J* = 10.1 Hz, 1H), 4.22 (dd, *J* = 12.2, 6.3 Hz, 1H), 4.07 (t, *J* = 5.8 Hz, 2H), 3.57 (t, *J* = 6.2 Hz, 2H), 3.41 (t, *J* = 6.0 Hz, 2H), 3.09 – 2.99 (m, 2H), 2.53 (s, 3H), 2.42 (s, 3H), 2.20 (d, *J* = 5.2 Hz, 5H), 2.00 (d, *J* = 7.6 Hz, 4H), 1.59 (s, 3H), 1.36 (s, 9H). $^{13}$C NMR (101 MHz, DMSO) δ 169.76, 169.66, 169.43, 159.33, 154.46, 149.26, 146.57, 139.00, 135.97, 134.00, 131.15, 116.37, 108.69, 77.58, 69.66, 68.75, 66.71, 61.49, 46.75, 43.78, 28.21, 20.76, 20.48, 20.08, 18.76, 14.08.

HRMS (ESI), m/z: calculated for $C_{34}H_{46}N_5O_{13}$ $[M+H]^+$: 732.3092, found: 732.3098.

### Synthesis of RFT-PEG4-DBCO (26)

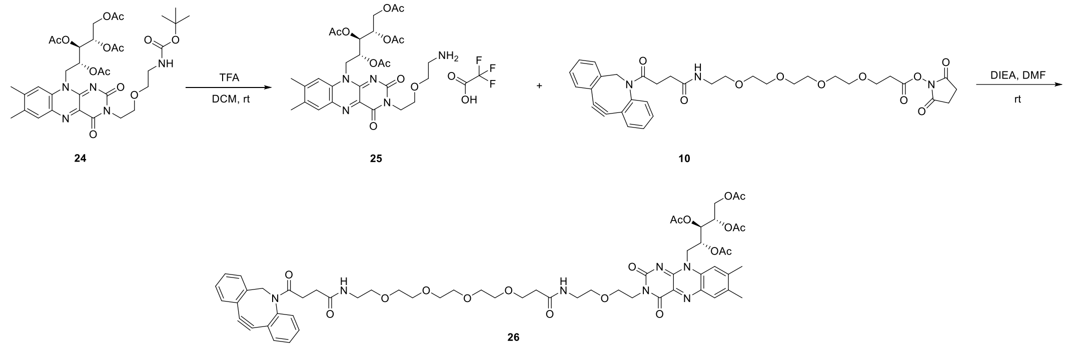

To a solution of (2*S*,3*R*,4*R*)-5-(3-(2-(2-((tert-butoxycarbonyl)amino)ethoxy)ethyl)-7,8-dimethyl-2,4-dioxo-3,4-dihydrobenzo[g]pteridin-10(2H)-yl)pentane-1,2,3,4-tetrayl tetraacetate **24** (0.082 mmol, 60 mg) in DCM (2 mL) was added excess trifluoroacetic acid (2 mL) the mixture was stirred at room temperature for 30 mins. LCMS showed complete conversion. The mixture was concentrated to complete dryness and the crude was used in the next step without purification. To a solution of the crude **25** in DMF (2 mL), DIPEA (0.5 mL) was added DBCO-PEG4-NHS Ester (0.082 mmol, 54 mg). The mixture was stirred overnight and concentrated over a stream of nitrogen. The crude was purified by ISCO using DCM : MeOH (100% to 90% to 50%) to give RFT-PEG4-DBCO **26** as yellow gum (79 mg, 83% over 2-steps).

$^1$H NMR (400 MHz, DMSO-$d_6$) δ 7.94 (s, 1H), 7.84 (t, *J* = 5.4 Hz, 1H), 7.76 (d, *J* = 8.7 Hz, 2H), 7.67 (d, *J* = 7.1 Hz, 1H), 7.61 (d, *J* = 7.3 Hz, 1H), 7.47 (q, *J* = 7.1 Hz, 3H), 7.35 (dt, *J* = 14.7, 7.2 Hz, 2H), 7.28 (d, *J* = 7.4 Hz, 1H), 5.48 (t, *J* = 8.2 Hz, 2H), 5.32 (d, *J* = 2.7 Hz, 1H), 5.02 (d, *J* = 14.0 Hz, 1H), 4.87 (d, *J* = 12.9 Hz, 1H), 4.38 (d, *J* = 10.0 Hz, 1H), 4.22 (dd, *J* = 12.4, 6.2 Hz, 1H), 4.08 (t, *J* = 6.2 Hz, 2H), 3.57 (dt, *J* = 13.8, 7.5 Hz, 5H), 3.51 – 3.39 (m, 14H), 3.33 (bs, 4H), 3.29 (t, *J* = 6.1 Hz, 2H), 3.17 (q, *J* = 5.5 Hz, 2H), 3.11 – 3.03 (m, 2H), 2.59 (s, 4H), 2.52 (s, 3H), 2.41 (s, 3H), 2.30 (t, *J* = 6.5 Hz, 2H), 2.20 (d, *J* = 4.8 Hz, 6H), 2.00 (s, 3H).

$^{13}$C NMR (101 MHz, DMSO) δ 172.76, 171.09, 170.11, 169.76, 159.36, 154.48, 151.60, 148.42, 136.08, 134.00, 132.41, 131.14, 129.61, 128.91, 128.11, 127.97, 127.66, 126.77, 125.13, 122.52, 121.39, 116.38, 114.20, 108.14, 69.75, 69.67, 69.65, 69.52, 69.49, 68.98, 68.76, 66.77, 61.48, 54.91, 43.80, 35.96, 30.33, 29.70, 25.22, 20.76, 20.47, 20.08, 18.76.

HRMS (ESI), m/z: calculated for $C_{59}H_{72}N_7O_{18}$ $[M+H]^+$: 1166.4934, found: 1166.4941.

### Synthesis of RFT-PEG3-Biotin (28) Biotin RFT

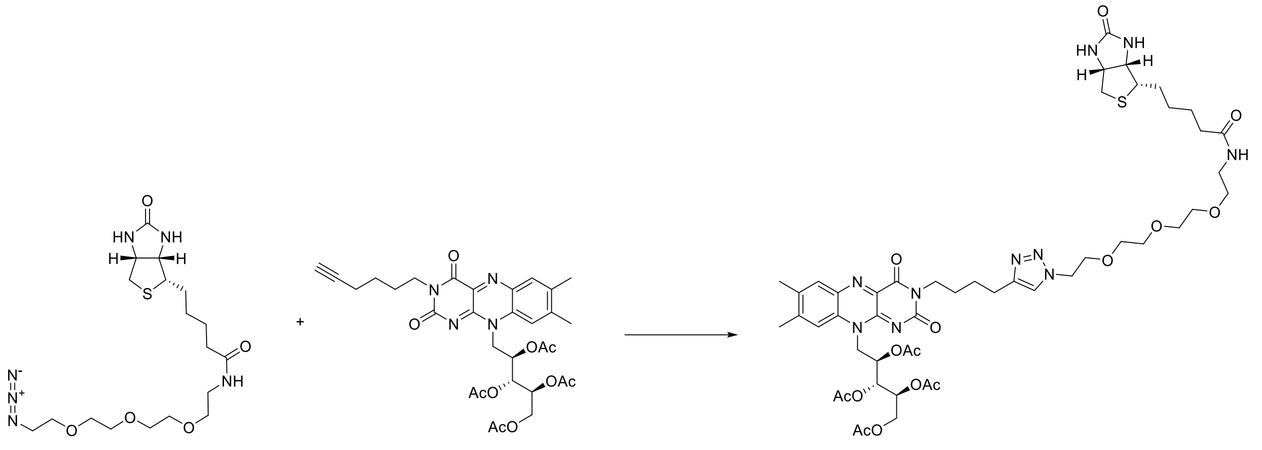

27 9 28, RFT-Biotin

A suspension of 5-(3-(hex-5-yn-1-yl)-7,8-dimethyl-2,4-dioxo-3,4-dihydrobenzo[g]pteridin-10(2H)-yl)pentane-1,2,3,4-tetrayl tetraacetate (1.0 equiv, 40 mg), sodium ascorbate (3 equiv, 38.1 mg) and biotin-PEG3-azide (1.2 equiv, 34.2 mg) in in DMF and water (1ml/0.5 mL) was added a premixed solution of copper sulfate, and tris(3-hydroxypropyltriazolylmethyl)amine in DMF and was stirred at room temperature for overnight. The crude mixture was concentrated by stream of nitrogen and purified by HPLC to give yellow solid of desired product (35 mg; Yield: 51.1%).

$^{1}$H NMR (400 MHz, DMSO-$d_6$) δ 7.95 (s, 1H), 7.80 (d, $J$ = 6.2 Hz, 2H), 7.76 (s, 1H), 6.41 (bs, 2H), 5.52 – 5.42 (m, 2H), 5.35 – 5.28 (m, 1H), 4.87 (d, $J$ = 13.3 Hz, 1H), 4.45 (t, $J$ = 5.1 Hz, 2H), 4.38 (d, $J$ = 10.1 Hz, 1H), 4.32 – 4.26 (m, 2H), 4.22 (dd, $J$ = 12.3, 6.2 Hz, 2H), 4.11 (dd, $J$ = 7.6, 4.5 Hz, 3H), 3.91 (m, 3H), 3.78 (t, $J$ = 5.1 Hz, 3H), 3.53 – 3.42 (m, 8H), 3.37 (t, $J$ = 5.8 Hz, 2H), 3.16 (q, $J$ = 5.6 Hz, 2H), 3.12 – 3.05 (m, 1H), 2.81 (dd, $J$ = 12.4, 4.9 Hz, 1H), 2.64 (s, 2H), 2.56 (d, $J$ = 12.4 Hz, 1H), 2.52 (s, 3H), 2.41 (s, 3H), 2.20 (d, $J$ = 5.4 Hz, 6H), 2.05 (t, $J$ = 7.4 Hz, 2H), 2.00 (s, 3H), 1.53 – 1.38 (m, 4H), 1.28 (dq, $J$ = 14.6, 7.7, 7.1 Hz, 2H).

$^{13}$C NMR (101 MHz, DMSO) δ 172.09, 170.11, 169.76, 169.66, 162.68, 159.26, 154.53, 146.44, 136.06, 133.96, 122.21, 69.68, 69.62, 69.56, 69.15, 68.78, 61.02, 59.18, 55.40, 49.20, 35.08, 28.18, 28.03, 27.04, 26.57, 25.24, 24.81, 20.77, 20.56, 20.47, 20.08, 18.76.

HRMS (ESI), m/z: calculated for $C_{49}H_{69}N_{10}O_{15}S$ $[M+H]^+$: 1069.4665, found: 1069.4670.

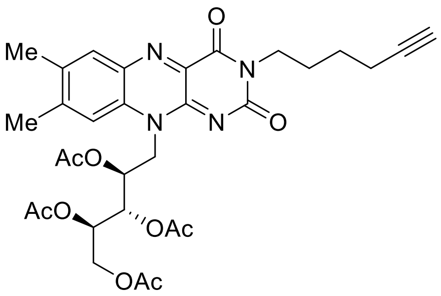

**9, Alkyne-RFT**

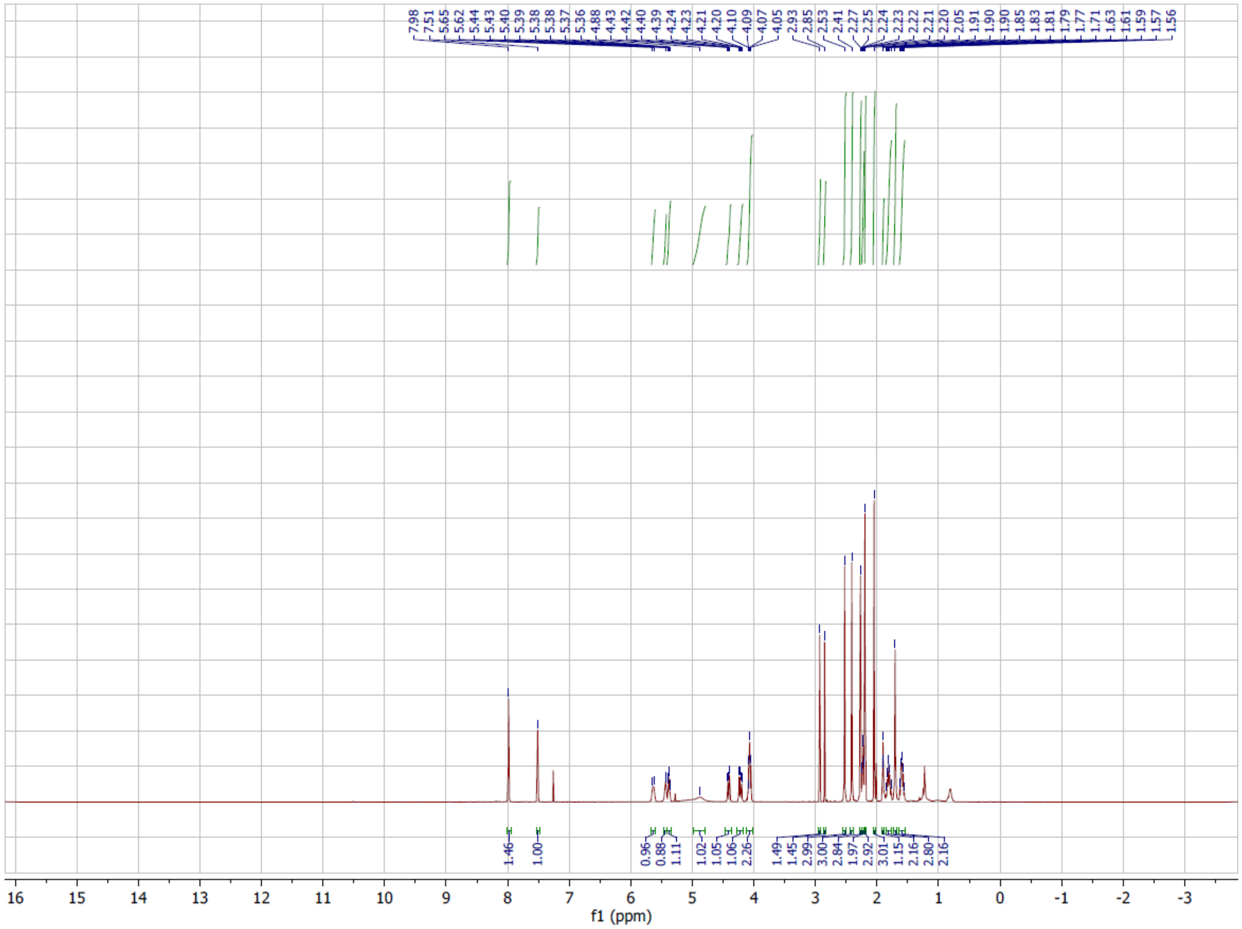

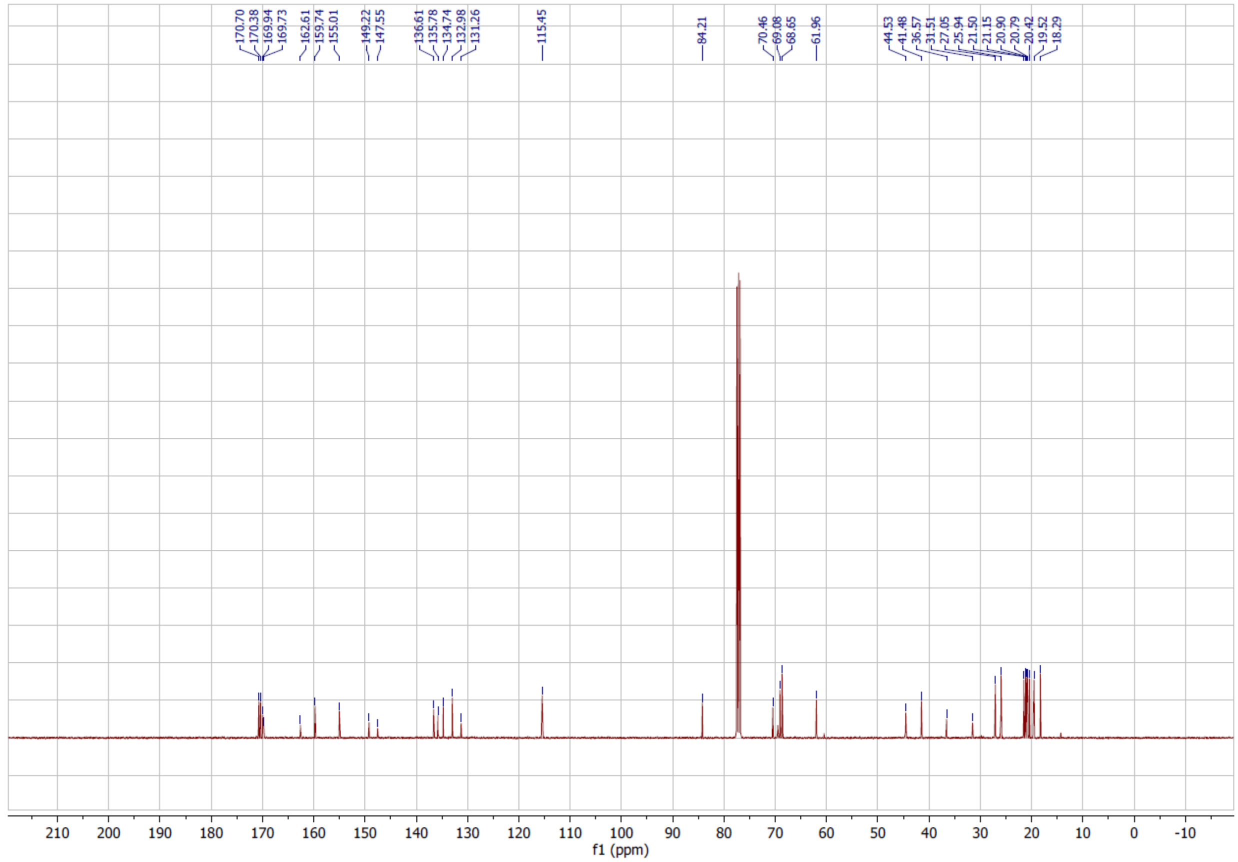
170.70
170.38
169.94
169.73
162.61
159.74
155.01
149.22
147.55
136.61
135.78
134.74
132.98
131.26
115.45
84.21
70.46
69.08
68.65
61.96
44.53
41.48
36.57
31.51
27.05
25.94
21.50
21.15
20.90
20.79
20.42
19.52
18.29
f1 (ppm)

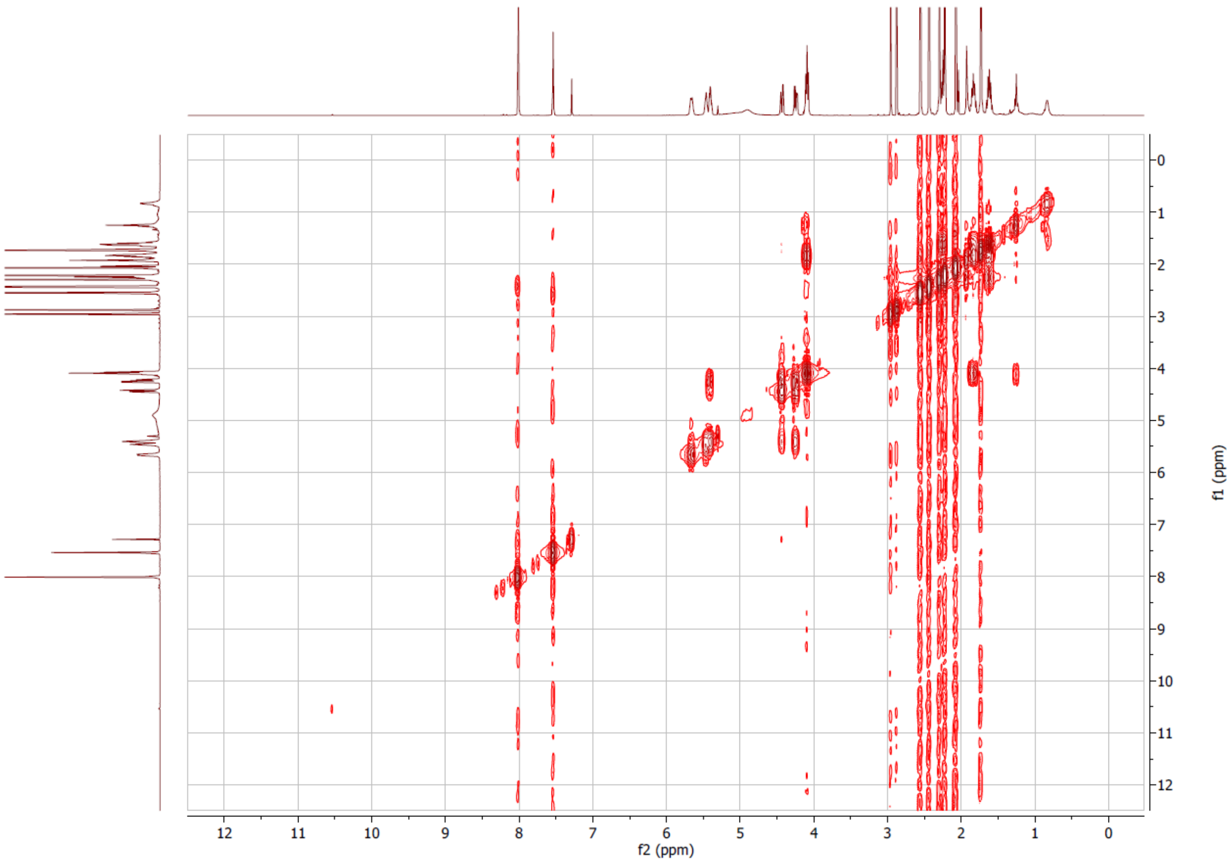
f2 (ppm)
f1 (ppm)

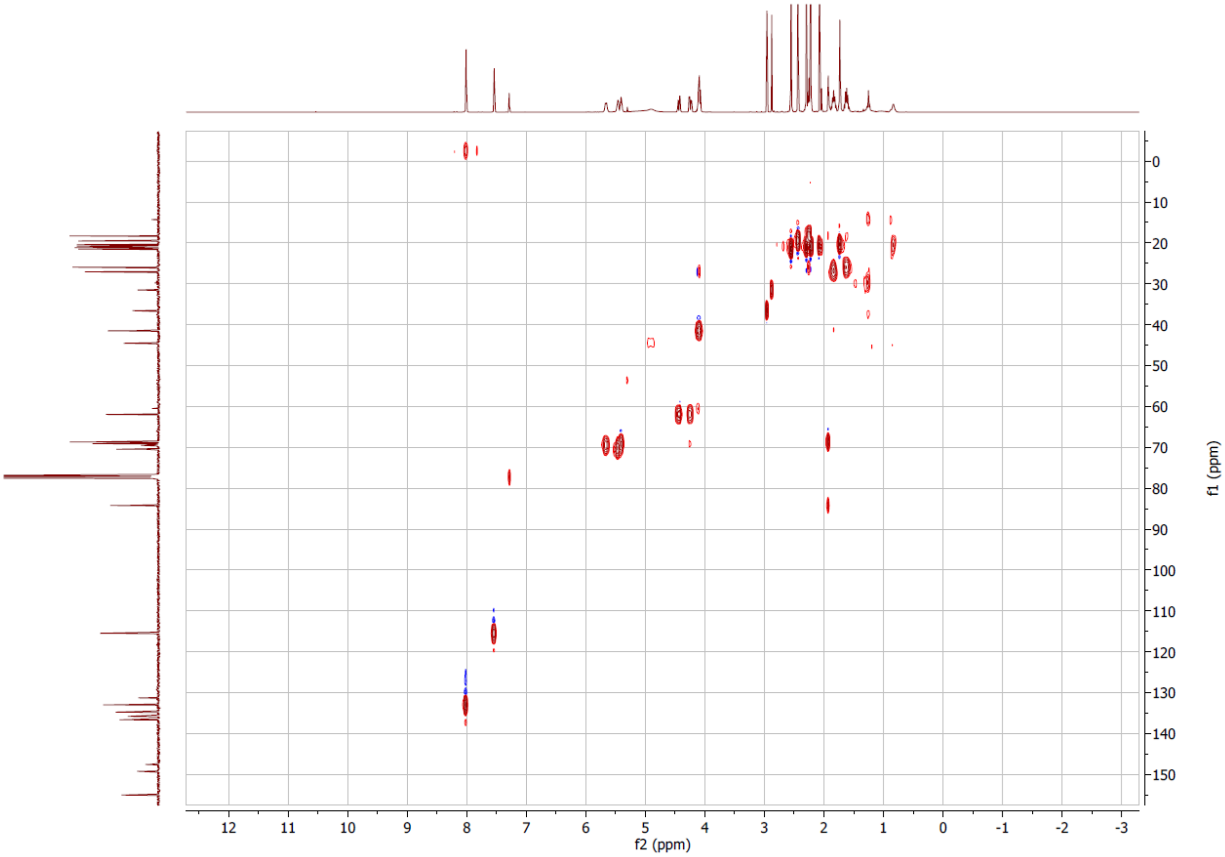
f2 (ppm)
f1 (ppm)

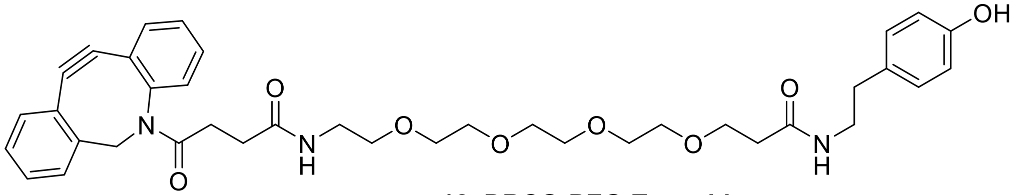

**12, DBCO-PEG-Tyramide**

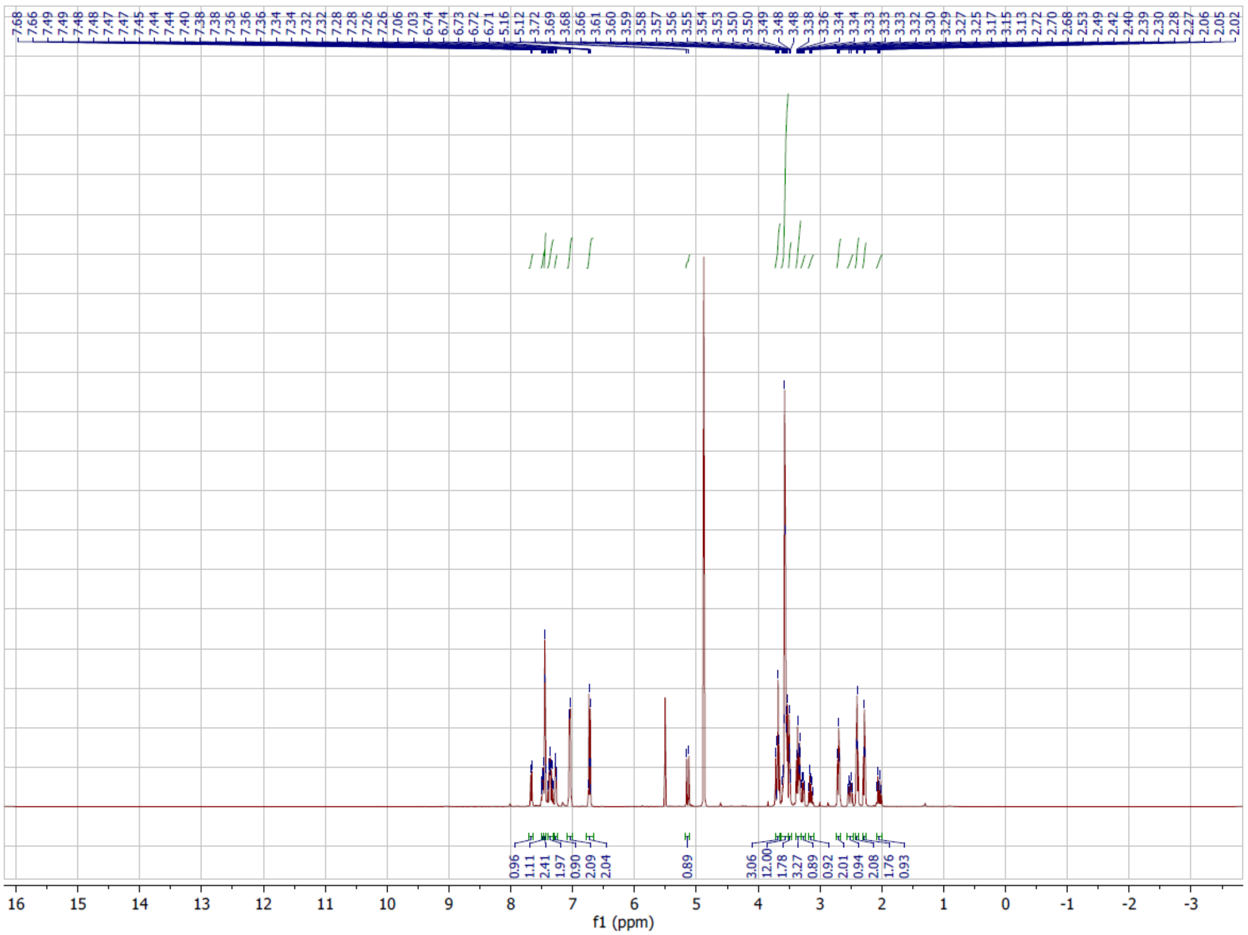

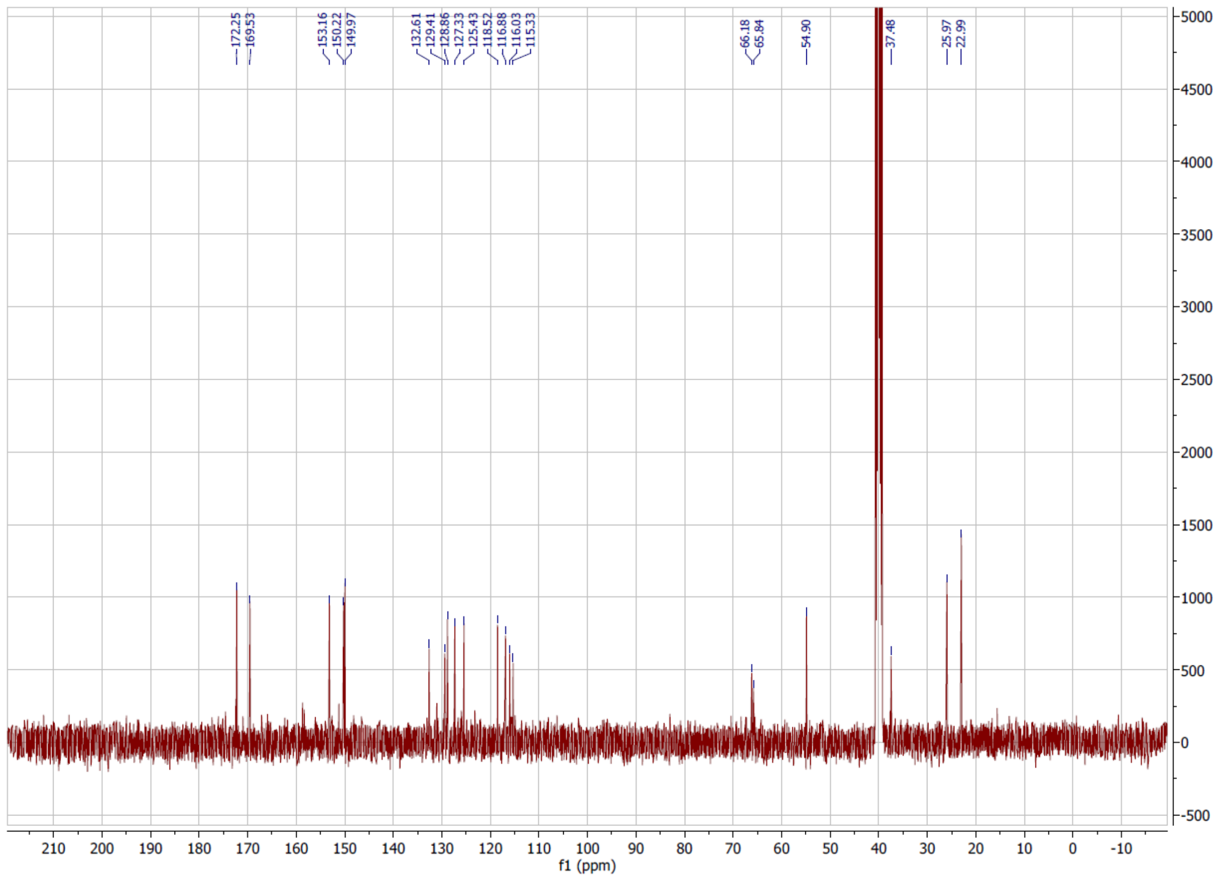
172.25
169.53
153.16
150.22
149.97
132.61
129.41
128.86
127.33
125.43
118.52
116.88
116.03
115.33
66.18
65.84
54.90
37.48
25.97
22.99
f1 (ppm)

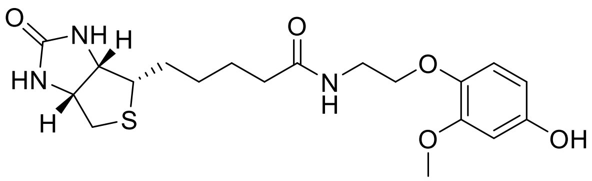

**15, 3MeO-EO-BP**

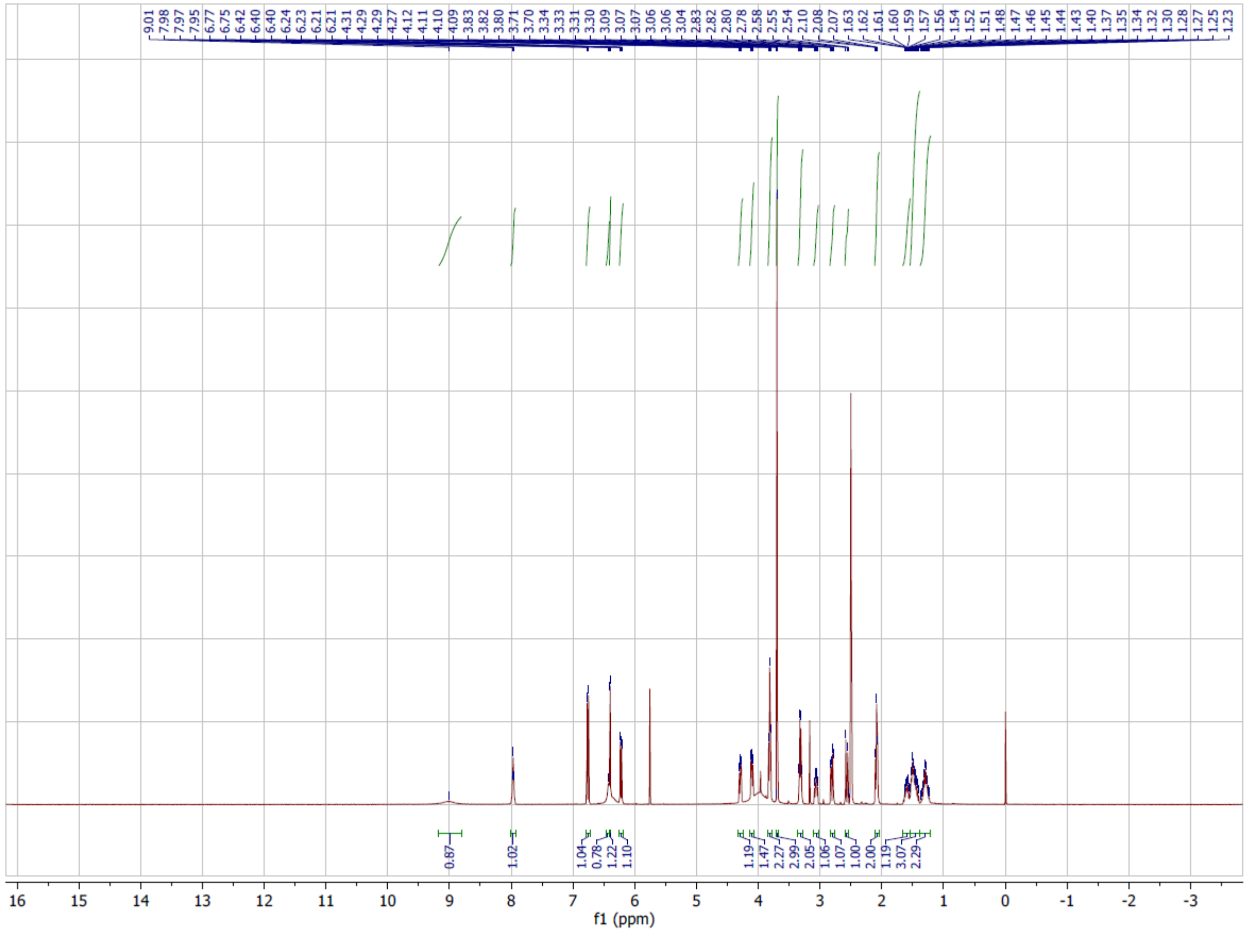

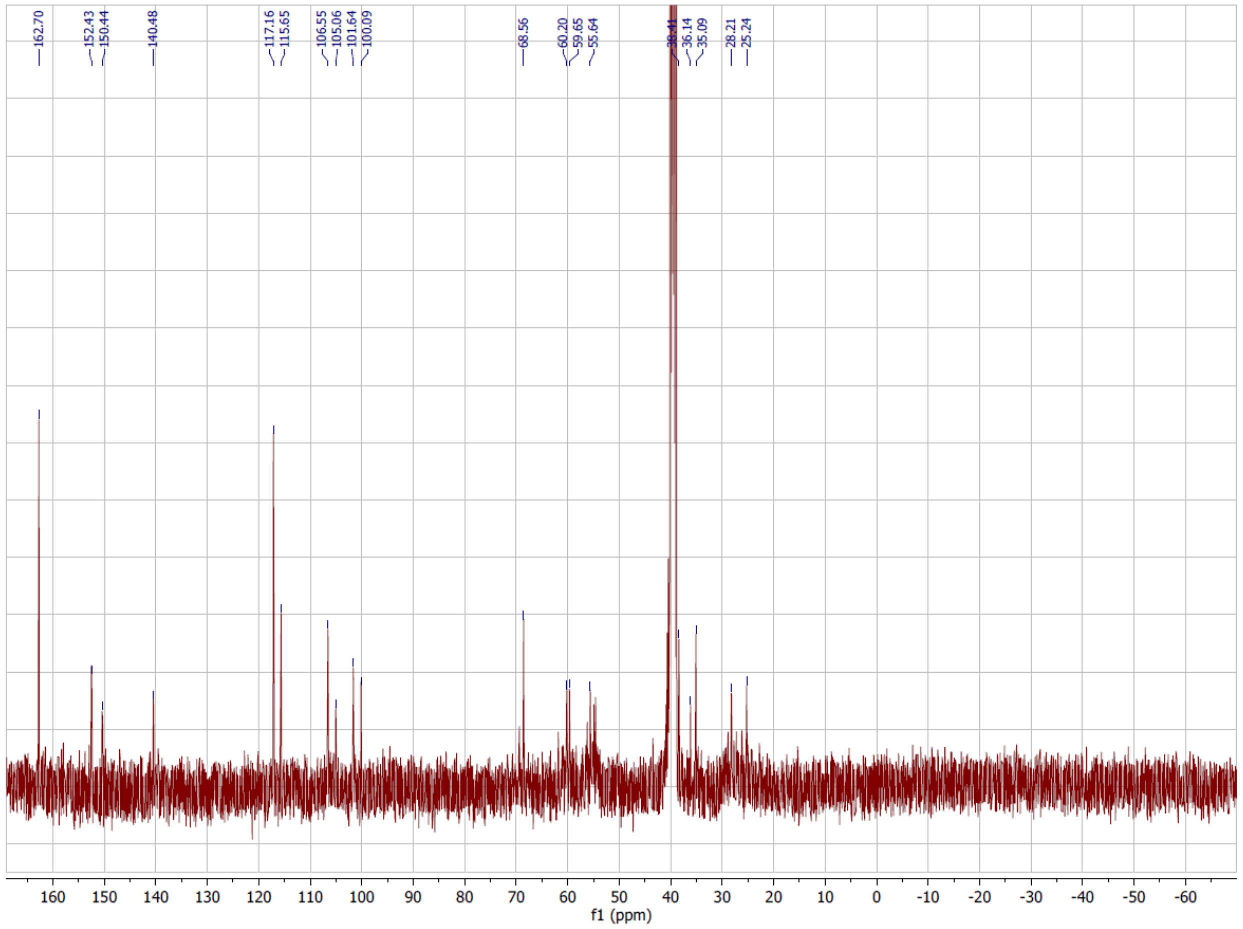
162.70
152.43
150.44
140.48
117.16
115.65
106.55
105.06
101.64
100.09
68.56
60.20
59.65
55.64
38.41
36.14
35.09
28.21
25.24
160 150 140 130 120 110 100 90 80 70 60 50 40 30 20 10 0 -10 -20 -30 -40 -50 -60
f1 (ppm)

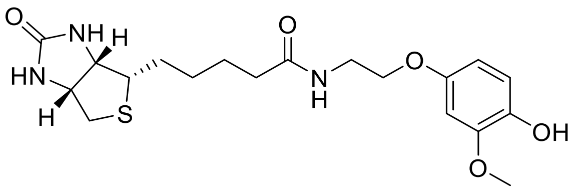

**18, 2MeO-EO-BP**

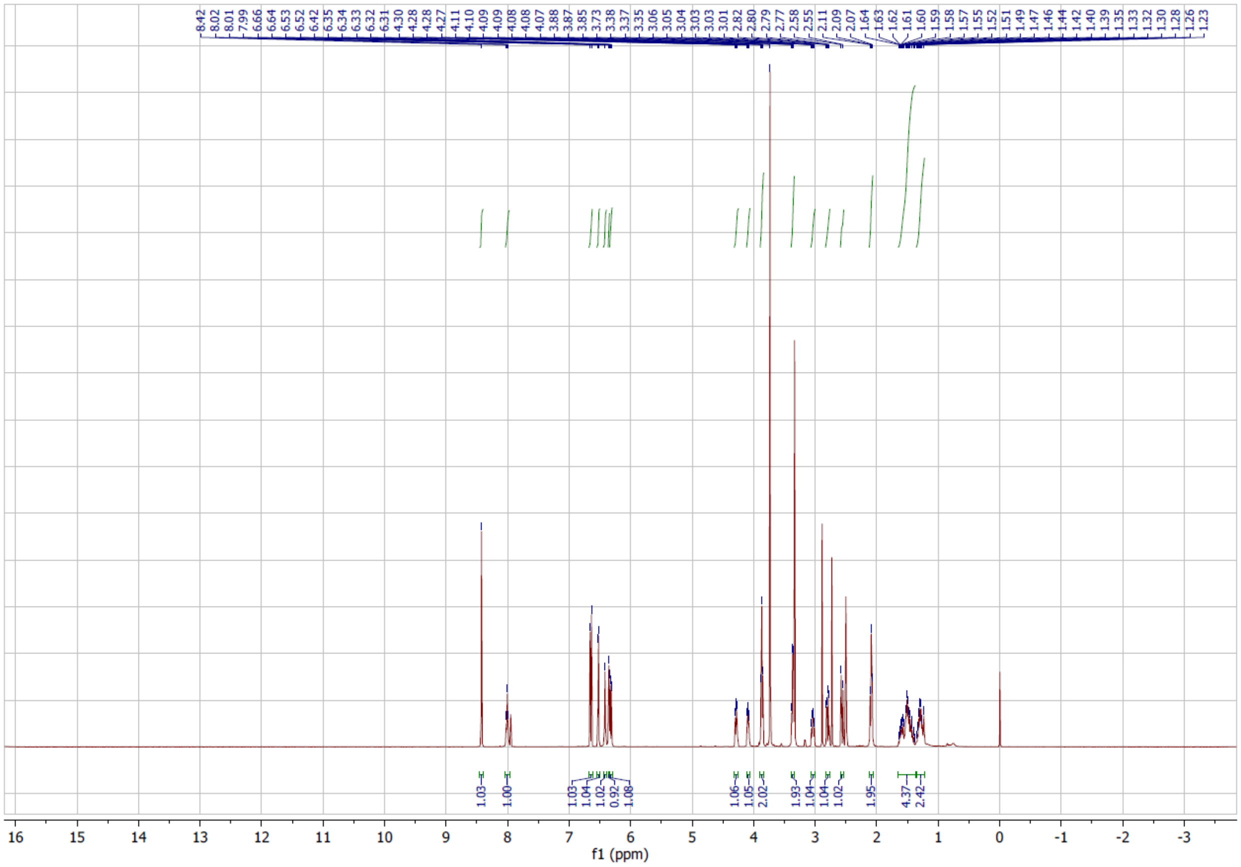

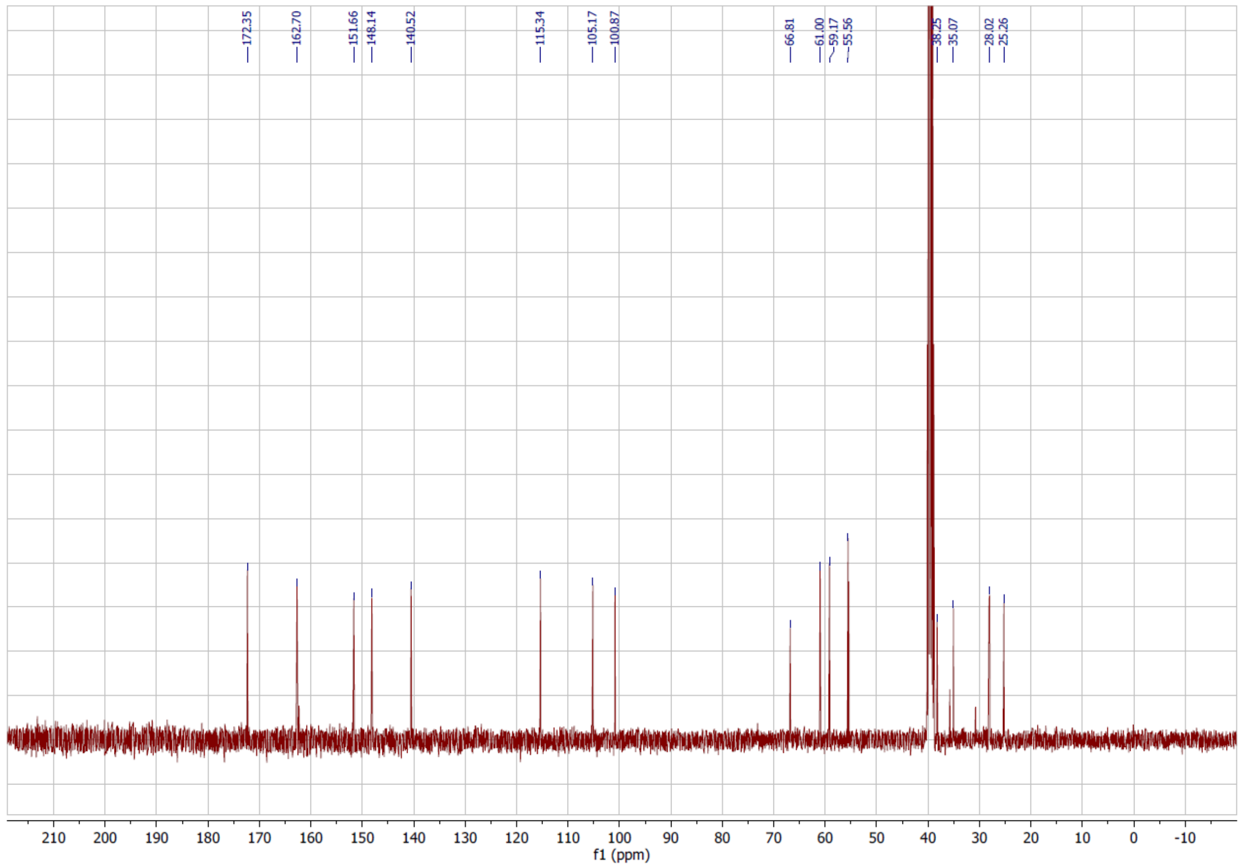
172.35
162.70
151.66
148.14
140.52
115.34
105.17
100.87
66.81
61.00
59.17
55.56
38.25
35.07
28.02
25.26
210 200 190 180 170 160 150 140 130 120 110 100 90 80 70 60 50 40 30 20 10 0 -10
f1 (ppm)

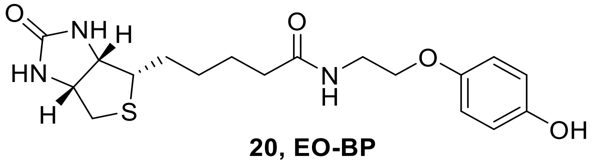

**20, EO-BP**

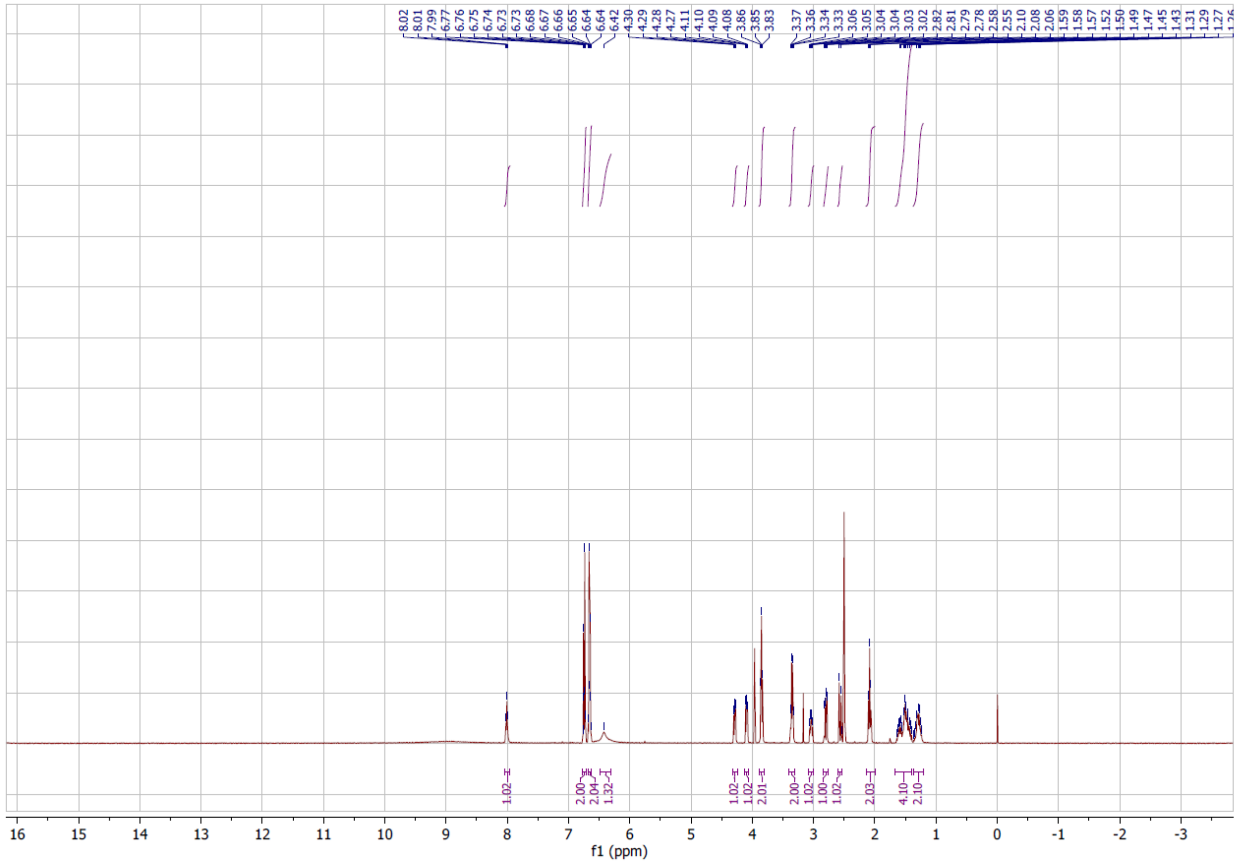

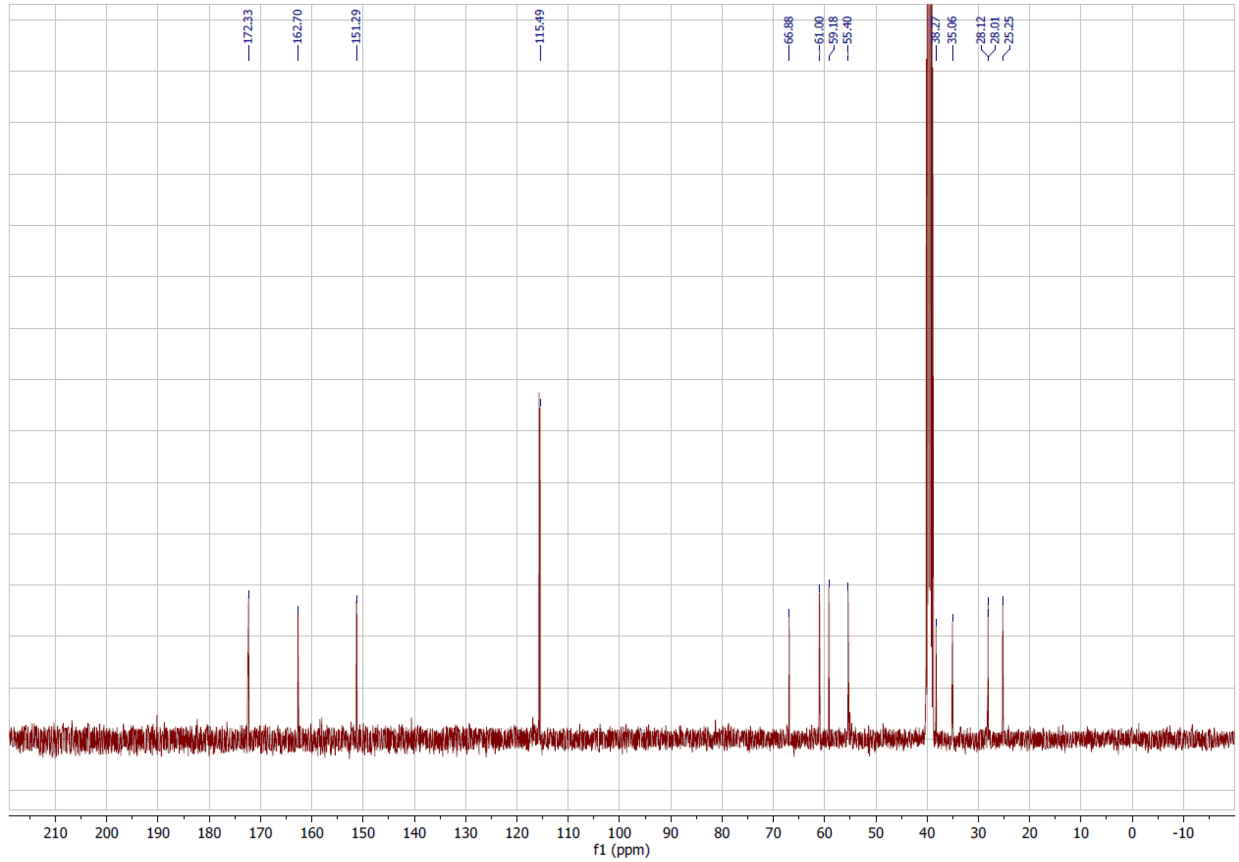

172.33
162.70
151.29
115.49
66.88
61.00
59.18
55.40
38.27
35.06
28.12
28.01
25.25
210 200 190 180 170 160 150 140 130 120 110 100 90 80 70 60 50 40 30 20 10 0 -10
f1 (ppm)

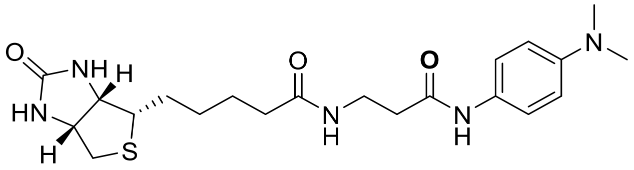

**22, Biotin-DMAA**

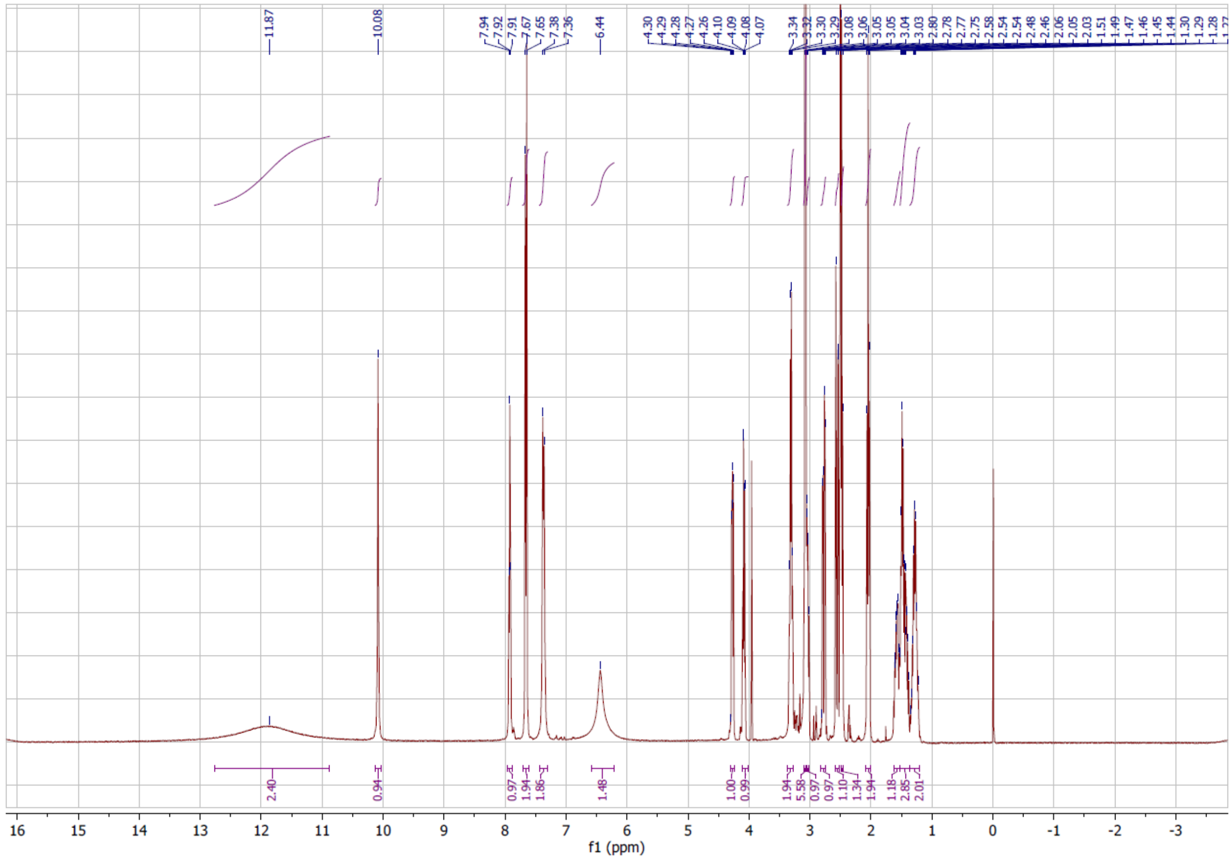

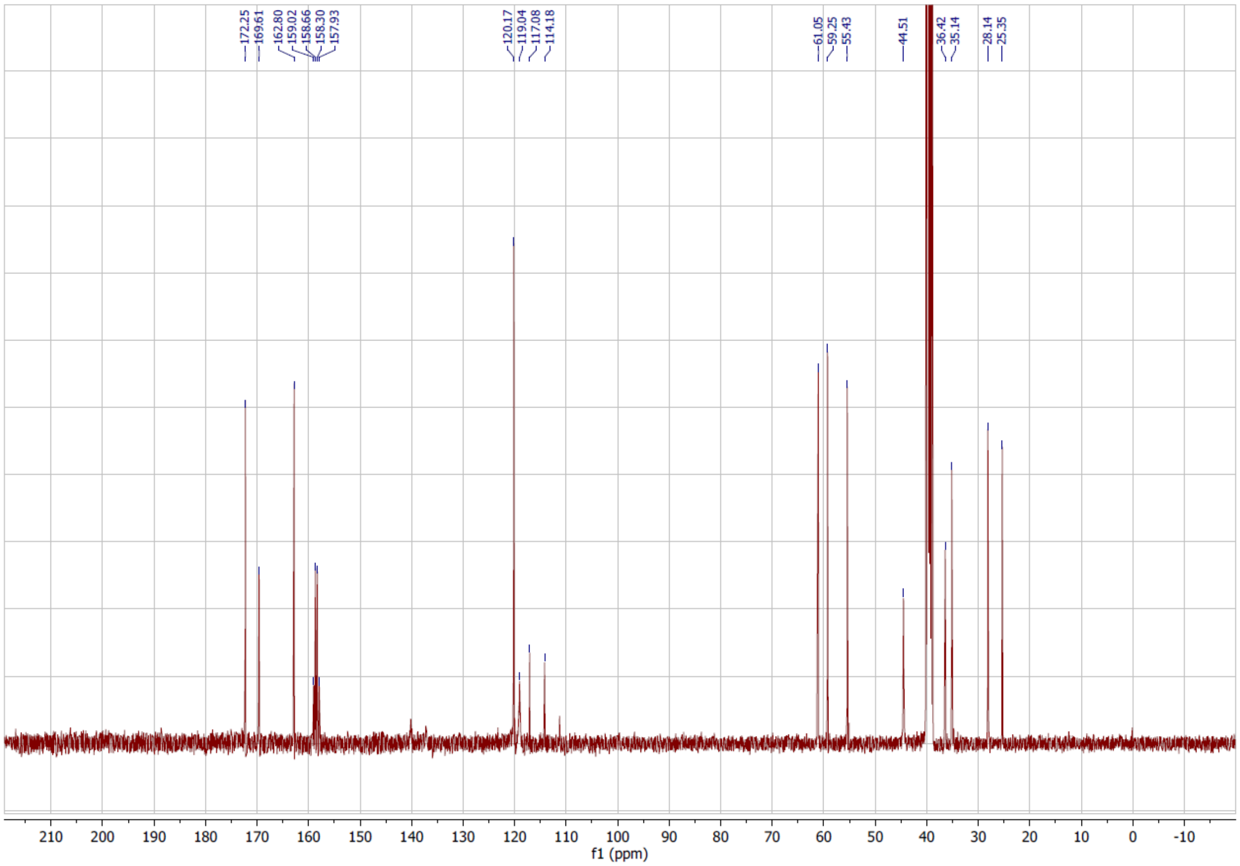
172.25
169.61
162.80
159.02
158.66
158.30
157.93
120.17
119.04
117.08
114.18
61.05
59.25
55.43
44.51
36.42
35.14
28.14
25.35
f1 (ppm)

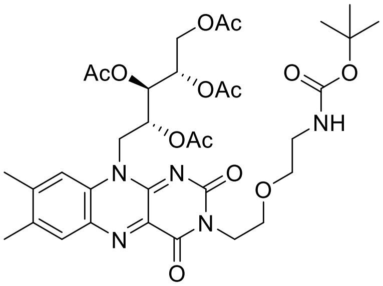

**24**

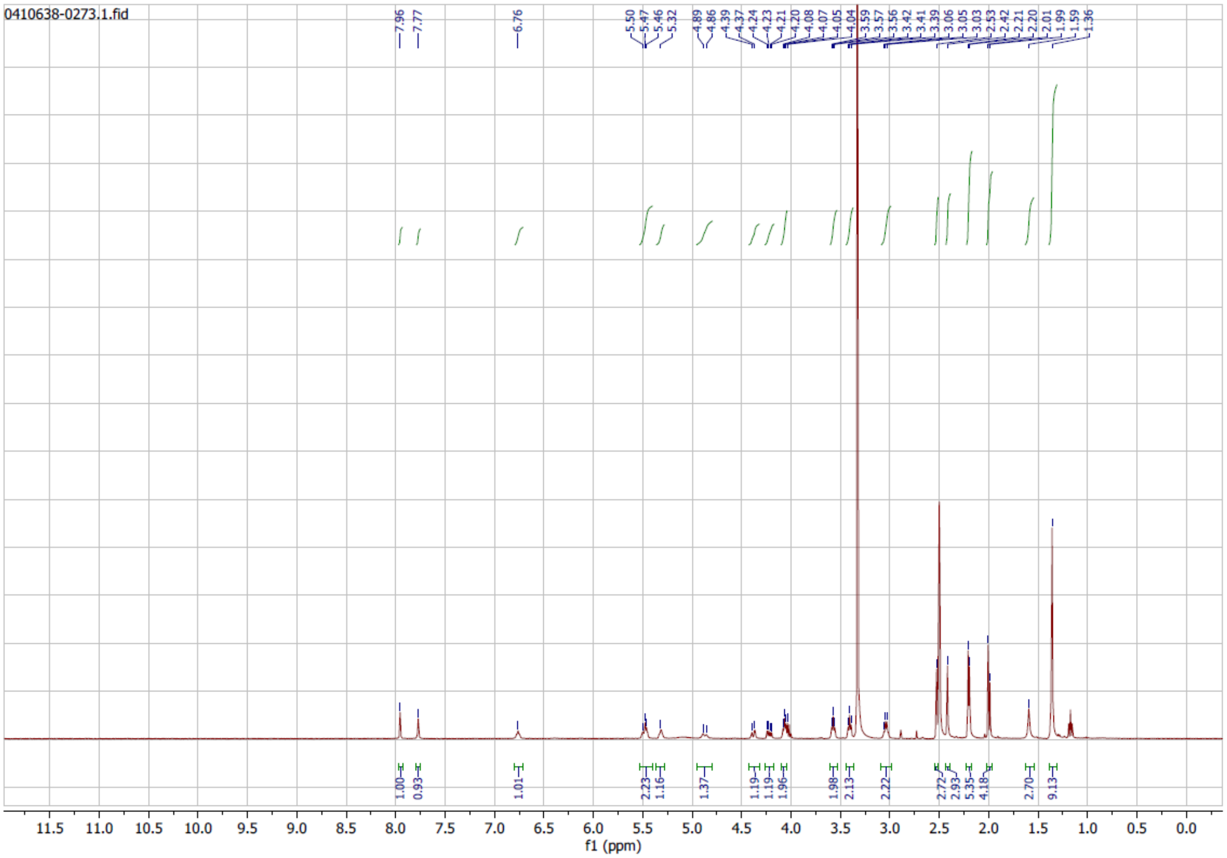

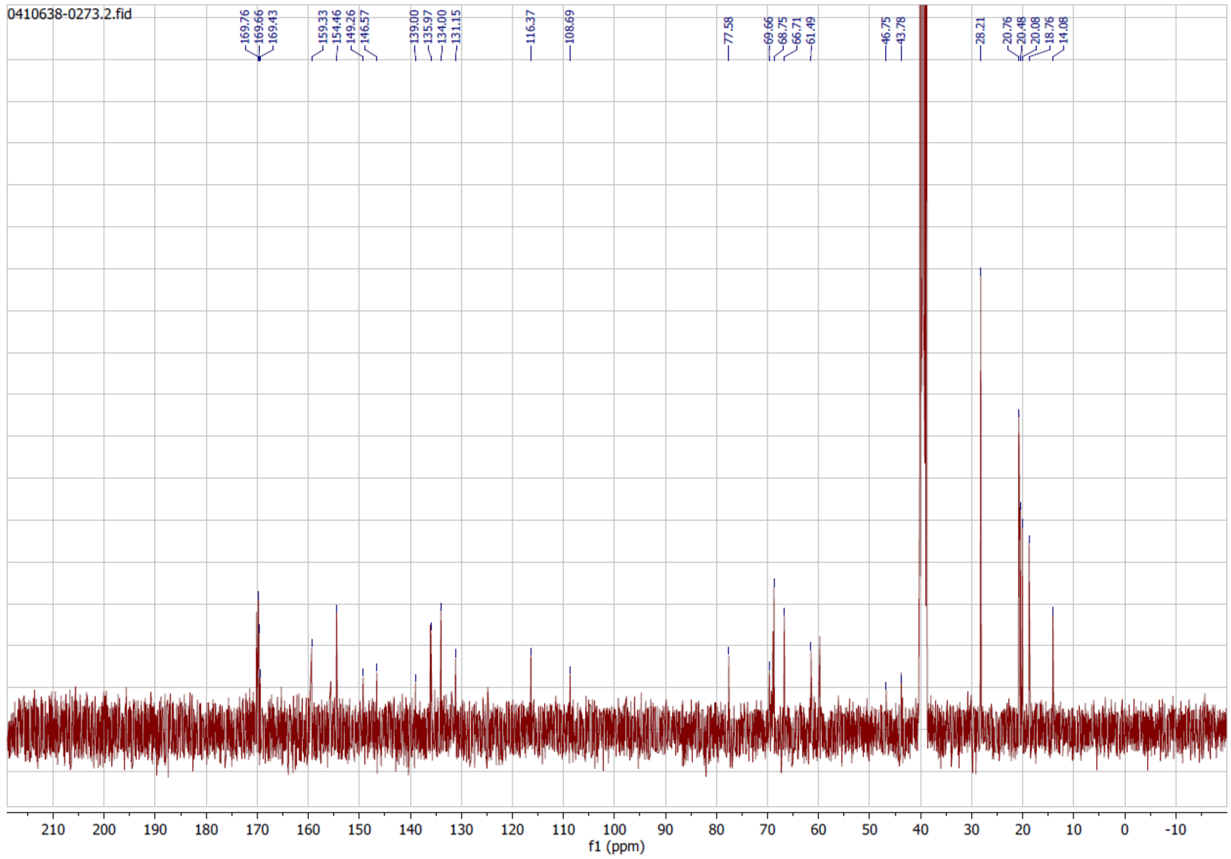
0410638-0273.2.fid
169.76
169.66
169.43
159.33
154.46
149.26
146.57
139.00
135.97
134.00
131.15
116.37
108.69
77.58
69.66
68.75
66.71
61.49
46.75
43.78
28.21
20.76
20.48
20.08
18.76
14.08
210 200 190 180 170 160 150 140 130 120 110 100 90 80 70 60 50 40 30 20 10 0 -10
f1 (ppm)

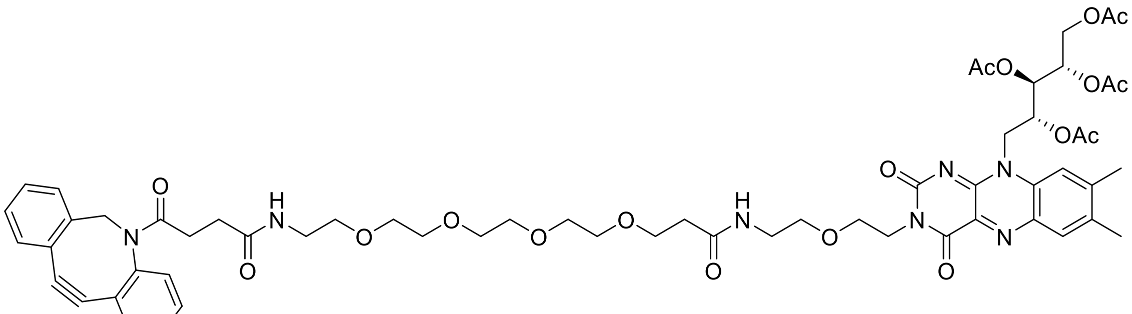

**26**

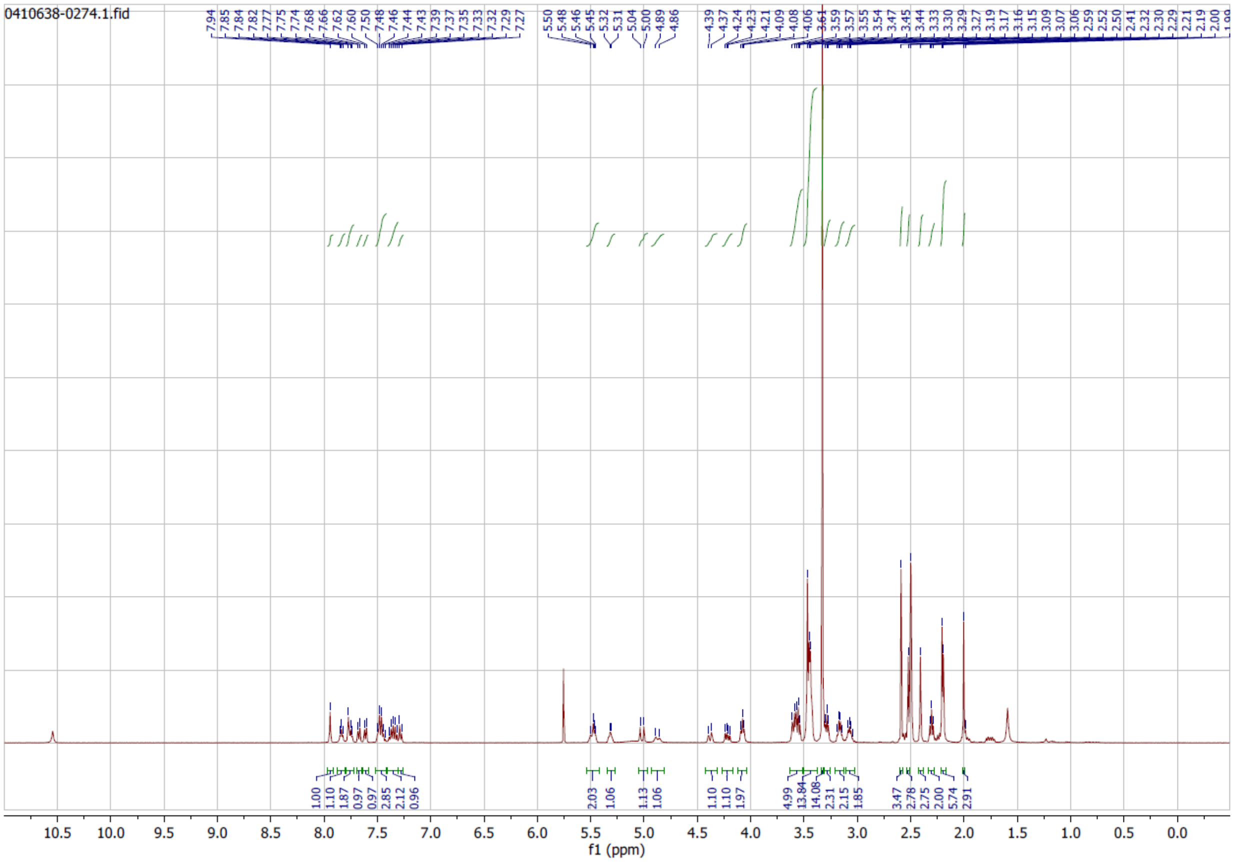

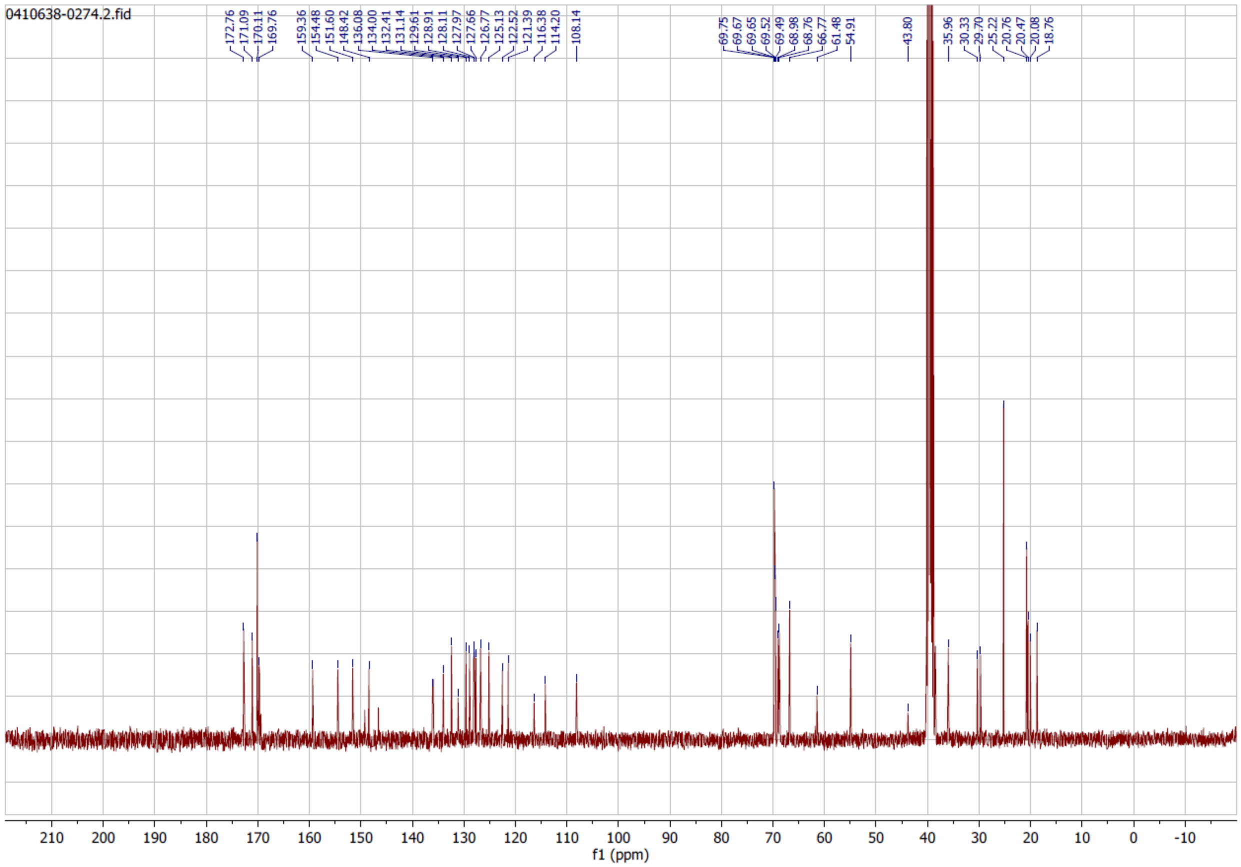
0410638-0274.2.fid
172.76
171.09
170.11
169.76
159.36
154.48
151.60
148.42
136.08
134.00
132.41
131.14
129.61
128.91
128.11
127.97
127.66
126.77
125.13
122.52
121.39
116.38
114.20
108.14
69.75
69.67
69.65
69.52
69.49
68.98
68.76
66.77
61.48
54.91
43.80
35.96
30.33
29.70
25.22
20.76
20.47
20.08
18.76
f1 (ppm)

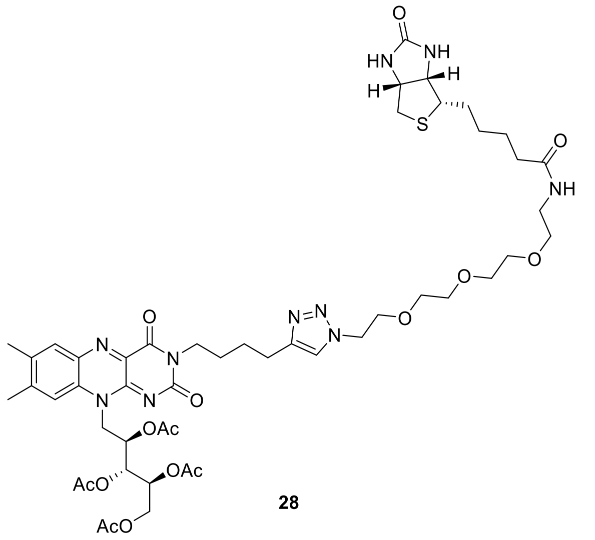

**28**

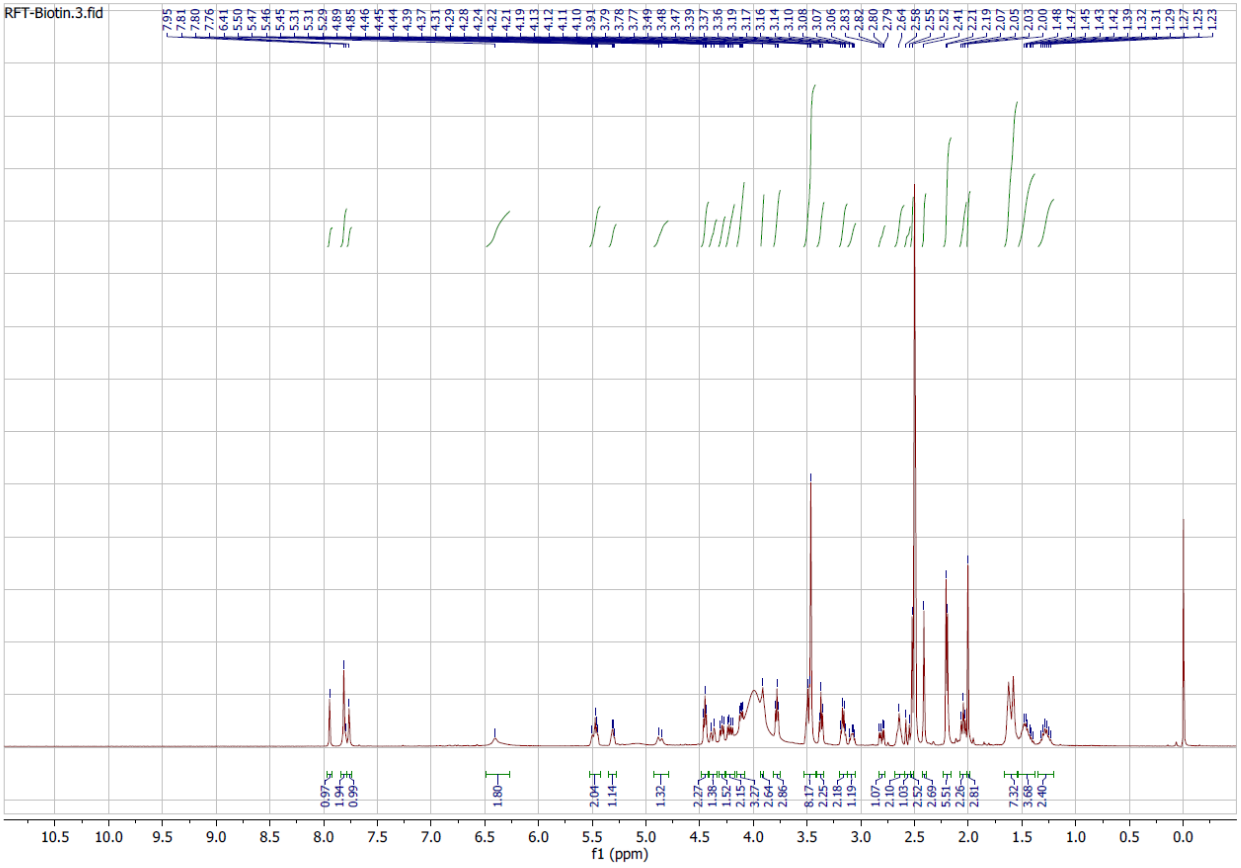

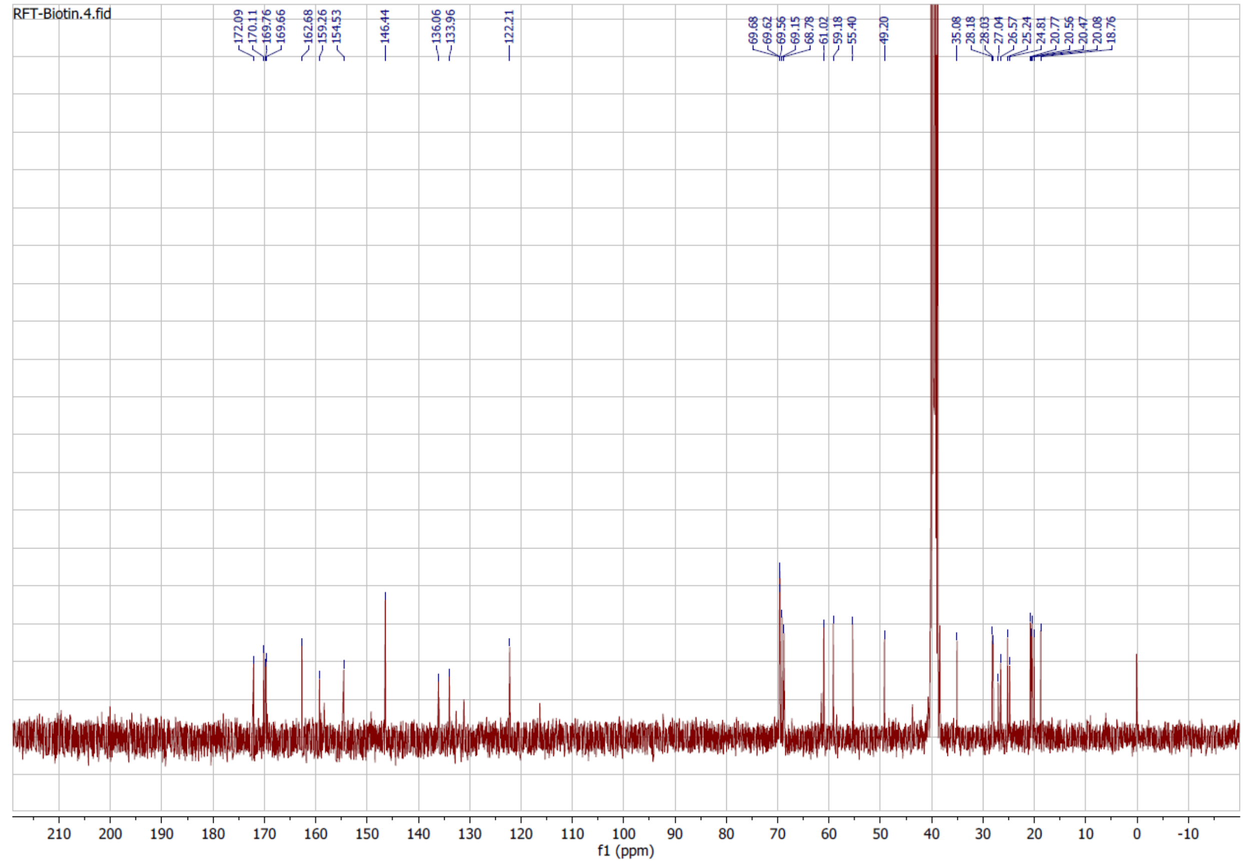
RFT-Biotin.4.fid
172.09
170.11
169.76
169.66
162.68
159.26
154.53
146.44
136.06
133.96
122.21
69.68
69.62
69.56
69.15
68.78
61.02
59.18
55.40
49.20
35.08
28.18
28.03
27.04
26.57
25.24
24.81
20.77
20.56
20.47
20.08
18.76
f1 (ppm)
